# Competing Programs Shape Cortical Sensorimotor-Association Axis Development

**DOI:** 10.1101/2025.06.26.660775

**Authors:** Jeremiah Tsyporin, Menglei Zhang, Cai Qi, Ashlea Segal, Thomas Finn, Hyojin Kim, Sang-Hun Choi, Xinyun Li, Sara Bandiera, Ivan Pavlovic, Suel-Kee Kim, Akemi Shibata, Kohei Onishi, Ziqin Zhang, Elijah Hammarlund, Graham Su, Nikkita Salla, Joy Kachko, Christi Hawley, Shuiyu Li, Daniel Z. Doyle, Xueyan Peng, Timothy Nottoli, Nuria Ruiz-Reig, Fadel Tissir, Yasushi Nakagawa, Erica Herzog, Shaojie Ma, Kevin Gobeske, Kartik Pattabiraman, Tomomi Shimogori, Alvaro Duque, Alex Fornito, Hao Huang, Mikihito Shibata, Bin Chen, Nenad Sestan

## Abstract

The neocortex is organized along a dominant sensorimotor-to-association (S-A) axis, anchored by modality-specific primary sensorimotor areas at one end and transmodal association areas that form distributed networks supporting abstract cognition at the other. The developmental mechanisms shaping this axis remain elusive. Here, we present converging multispecies evidence supporting the Multinodal Induction-Exclusion in Network Development (MIND) model, in which S-A patterning is governed by competing processes of induction and exclusion, driven by opposing transcriptomically-defined identity programs emerging from different nodes. Key molecular and connectional features of association cortices arise through pericentral programs, originating around fronto-temporal poles and partially regulated by retinoic acid. They progress inward toward central territories of the naïve neocortex along fronto-temporally polarized trajectories. Central programs are induced through interactions between topographically separated first-order sensorimotor thalamocortical inputs and the neocortex, promoting the formation of primary areas while excluding pericentral programs. Influenced by SATB2 and ZBTB18, these evolutionarily conserved programs compete for the same territory and create spatial compartmentalization of axon guidance, cell-cell adhesion, retinoic acid signaling, synaptogenesis, Wnt signaling, and autism risk genes. Notably, PLXNC1 and SEMA7A exhibit anti-correlated expression and repulsive functions in shaping cortico-cortical connectivity along the S-A axis. These processes of induction and exclusion establish an S-A equilibrium and topography in which primary sensorimotor areas emerge as focal islands within the broader ocean of distributed associative networks. The MIND model provides a unifying framework for understanding experimental, evolutionary, and clinical phenomena, revealing induction and exclusion as antagonistic complementary principles shaping the S-A axis and processing hierarchies.

## Introduction

A central challenge in neuroscience concerns understanding how large-scale networks of the cerebral cortex develop a modular yet hierarchically organized architecture to support complex functions^1^. Numerous studies have shown that multiple aspects of cortical structure and function are organized along a dominant sensorimotor-to-association (S-A) axis^1–3^. At one end of this axis, modality-specific primary sensory and motor areas form dense local networks that process sensory inputs or execute motor behaviors^4–6^. At the other end, transmodal association hubs in the prefrontal, posterior parietal, and temporal cortices are interconnected by distributed networks of long-range axons, enabling information integration across modalities to support complex and abstract cognition^5–10^. These transmodal networks strategically avoid primary sensorimotor areas while interacting with (para)limbic (peri)allocortical distributed networks processing emotions and memories^1,6,7,11^.

Topographically, primary unimodal areas appear as focal islands within a vast ocean of association cortex, creating the distinct spatial organization of the S-A axis^12^. Along this axis, laminar differentiation and functional specialization progressively decline, while synaptic plasticity and inter-individual variability increase with greater physical distance from unimodal areas^1,8,9,11^. Accordingly, boundaries between primary and adjacent areas are sharply defined but become more gradual in transmodal regions. While this organizational framework is conserved across mammals, the relative proportion of unimodal versus transmodal cortex varies significantly between species, reflecting in part adaptive specializations^12^. In humans, transmodal cortices have undergone the greatest evolutionary expansion relative to non-human primates and exhibit neoteny during postnatal development, which is thought to be linked to both our enhanced cognitive capacities and increased susceptibility to certain neuropsychiatric and neurodegenerative conditions^3^.

Foundational understanding of neocortical arealization and circuit development comes largely from studies of sensorimotor regions that focus on the interplay between intrinsic molecular gradients within progenitor cells and the extrinsic influences of peripheral sensory signals conveyed through first-order (FO) thalamocortical afferents (TCAs)^13–21^. As TCAs navigate toward the cortex, they preserve a topological register, establishing foundational sensorimotor areal maps^15,22^. The FO TCAs drive the patterning of primary cortical areas through a combination of spontaneous patterned activity, originating from sensory organs (e.g., spontaneous retinal waves), and stimulus-evoked activity (e.g., eye opening)^14,15,18,19,23^. These activity-dependent signals interact with immature neocortical neurons to induce key primary area-specific properties, including gene expression, cytoarchitecture, and cortico-cortical connectivity. Their spatially constrained and inductive properties serve as patterning anchors for activity-dependent mechanisms that, in a sensoritopic manner, also shape surrounding prospective association networks across multiple levels of the S-A hierarchy. Furthermore, enhanced peripheral sensory or motor capabilities can expand neocortical sensorimotor representations, while the absence of primary sensory inputs often leads to the adoption of adjacent association-like properties^12–19,21,22,24,25^.

The evolutionary trajectories of S-A patterning are reflected in species differences. Monotremes and marsupials, the most ancient living mammals, possess small neocortices dominated by unimodal areas^12^. In contrast, primates show marked expansion of transmodal cortex, supporting the emergence of unique structural, functional, and developmental adaptations along the S-A axis^3,5^.

Despite this progress, the development and evolution of transmodal cortices remain poorly understood compared to primary and early association areas^1,3,5,8–12^. Several key questions remain: (1) Are transmodal cortices patterned by the same mechanisms that shape unimodal cortex?; (2) How do transmodal neurons establish long-range connections across vast distances while avoiding primary areas?; (3) Why do distributed transmodal networks feature fronto-parieto-temporal transmodal hubs and frequently engage the prefrontal cortex (PFC)?; (4) Why do association areas exhibit graded transitions, in contrast to the sharply defined boundaries delimiting primary areas?; and (5) Has the evolutionary expansion of transmodal cortex in primates necessitated distinct developmental mechanisms?^5^

To address these questions, we present converging lines of evidence to support a new model of S-A axis development, which we named the Multinodal Induction–Exclusion in Network Development (MIND) model. This model posits that competing transcriptomically-defined programs, emerging around the neocortical center and the pericentral fronto-temporal (F-T) poles, differentiate primary unimodal sensorimotor areas from surrounding association networks through complementary processes of induction and exclusion. Together, these dual processes provide a foundational mechanism for establishing the hierarchical organization of the neocortex.

## Results

### Emerging transcriptomic signatures along the developing S-A axis

Prior evidence shows that transcriptomic differences between developing neocortical areas are temporally regulated, with a transient peak in inter-areal gene expression variability occurring during early to mid-fetal development, a crucial period for cell differentiation and neural circuit formation^26–30^. This phase is marked by prominent opposing transcriptomic gradients in the prospective association cortices of the frontal and temporal lobes, alongside more localized patterns in the prospective primary sensorimotor areas, despite the absence of obvious cytoarchitectonic distinctions^26–30^. Based on these findings, we hypothesize that these transcriptomic patterns reflect distinct developmental programs that shape prospective primary sensorimotor and association cortices through separate mechanisms, whose interaction gives rise to the hierarchically organized S-A axis.

To investigate this hypothesis, we generated gene modules (GMs) in an unsupervised manner based on co-expression enrichment in prospective primary sensorimotor or association cortices during fetal periods (p3-7, see **Supplementary Table 1** for developmental periods defined initially in Kang et al., 2011^27^) using data from human and macaque brains. We analyzed three independently generated datasets utilizing two transcriptomic modalities: Kang et al., 2011^27^ (accessed through humanbraintranscriptome.org), human microarray; Li et al., 2018^29^ (accessed through BrainSpan.org), human RNA-seq; Zhu et al., 2018^30^ (accessed through PsychENCODE.org) macaque RNA-seq. These studies sampled four putative primary areas including motor (M1C), somatosensory (S1C), visual (V1C), and auditory (A1C); seven association areas including PFC (medial, MFC; orbital, OFC; dorso-lateral, DFC/dlPFC; ventral, VFC), inferior-posterior parietal (IPC), and superior and inferior temporal cortex (STC, ITC); and limbic paleocortical amygdala (AMY) and archicortical hippocampus (HIP) (**Fig. 1a** and **Extended Data Figs. 1,2a,3a**). A subset of GMs were generated based on the enrichment of gene expression in groups of prospective association or primary sensorimotor areas (**Methods**). Association areas (A) were grouped into A frontal (Af) corresponding to prospective PFC (MFC, OFC, DFC, VFC), and A temporal (At), corresponding to prospective temporal (STC, ITC) association areas during p3-7 (**Supplementary Tables 2-5**). We similarly generated four primary sensorimotor (S)-enriched GM submodules corresponding to primary motor (Sm1), somatosensory (Ss1), visual (Sv1), and auditory (Sa1) areas. We generated GMs using each dataset independently (**Supplementary Tables 2-4**), and curated a stringent subset of genes that were present in all three independent datasets which we refer to as shared GMs (**Supplemental Table 5**).

**Figure 1:**
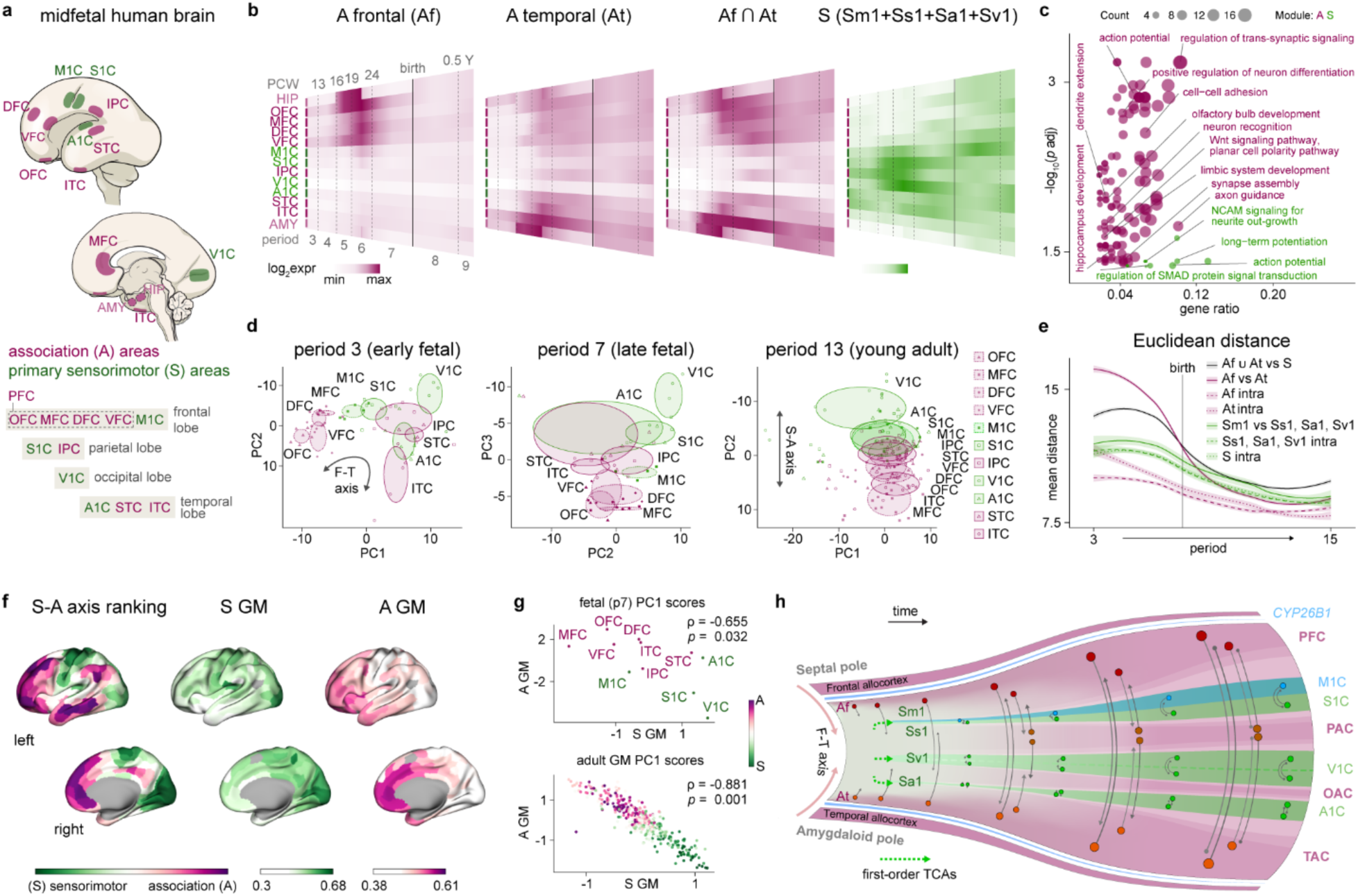
Anti-correlated pericentral and central gene modules define the S-A axis and MIND model. **a,** Depiction of mid-fetal human brain with analyzed regions highlighted. **b,** Shared gene module (GM) heatmaps displaying minimum to maximum median mean value of samples of a given region and period for genes belonging specifically to association frontal (Af) or association temporal (At) modules, genes shared between Af and At (Af ∩ At), or genes present in one of all four sensorimotor (S) modules, across 13 sampled regions and developmental periods (p3–p9). Since samples spanned a range of time points within each period and periods do not represent sharp developmental transitions, we generated a smooth gradient between periods. Given that the medial frontal neocortex borders the rudimentary hippocampus (allocortex), we placed HIP on the frontal side and AMY on the temporal side of the heatmap. **c**, Gene set enrichment analysis (GSEA) bubble plot for shared A (Af + At) and S (Sm1, Ss1, Sa1, Sv1) GMs. **d,** Principal component analysis (PCA) with all genes in the shared S and A GMs plotted against the indicated principal components. Ellipses are centered on the mean of the points of the indicated region and the size of each axis is determined based on the standard deviation of each component. In p7, PC1 reflects sample quality (PMI) due to the smaller sample size and was therefore not plotted. **e,** Euclidean distance based on all genes in the shared S and A GMs between indicated regional groups at each developmental period. Confidence interval level 0.95. **f,** Cortical surface renderings showing individual parcel ranking of the archetypal S-A axis^3^ (S-A axis ranking; green, sensorimotor-like; pink, association-like); and the mean adult gene expression across genes in the S GM or A GM, respectively^32^**. g,** Scatterplots showing the relationship between PC1 in the S GM and the A GM for p7 (top) and adult (bottom). For p7 (top), regions are colored according to predefined S-A axis definitions shown in **a**., for adult (bottom), regions are colored according to position along the archetypal S-A axis. **h,** Graphical depiction of the MIND model.

Gene set enrichment analysis of either shared or individual dataset GMs revealed enrichments for processes including axon guidance (e.g., plexins and semaphorins), cell-cell adhesion (protocadherins), limbic system development, trans-synaptic signaling, positive regulation of neuronal differentiation, retinoic acid signaling, and Wnt signaling represented in the A (combined and/or individual Af and At) GMs; in the S GM (combined Sm1, Ss1, Sa1, and Sv1), action potential generation, long-term potentiation, SMAD signaling, and NCAM-mediated neurite outgrowth processes among others were enriched (**Fig. 1c, Extended Data Figs. 4, and Supplementary Tables 6-9**). Furthermore, we observed a consistent enrichment of autism spectrum disorder (ASD) risk genes across GMs and datasets (**Extended Data Fig. 5 and Supplementary Tables 10-13**). Analysis of single-cell RNA-seq data from prospective primary and association midfetal human neocortical areas^31^ revealed that S and A GM genes are expressed across diverse cell types, including both excitatory projection neurons (ExNs) and inhibitory neurons (InNs), and exhibited distinct laminar and regional/areal patterns (**Extended Data Fig. 6**). Moreover, several S and A GM genes involved in axon guidance and cell-cell adhesion, such as *Pcdh10*, *Pcdh17*, *Plxnc1*, and *Sema7a*, exhibited anti-correlated and complementary expression patterns across layers and areas of the emerging mouse S-A axis (**Fig. 2 and Extended Data Fig. 7**), suggesting that these are largely conserved developmental programs.

**Figure 2:**
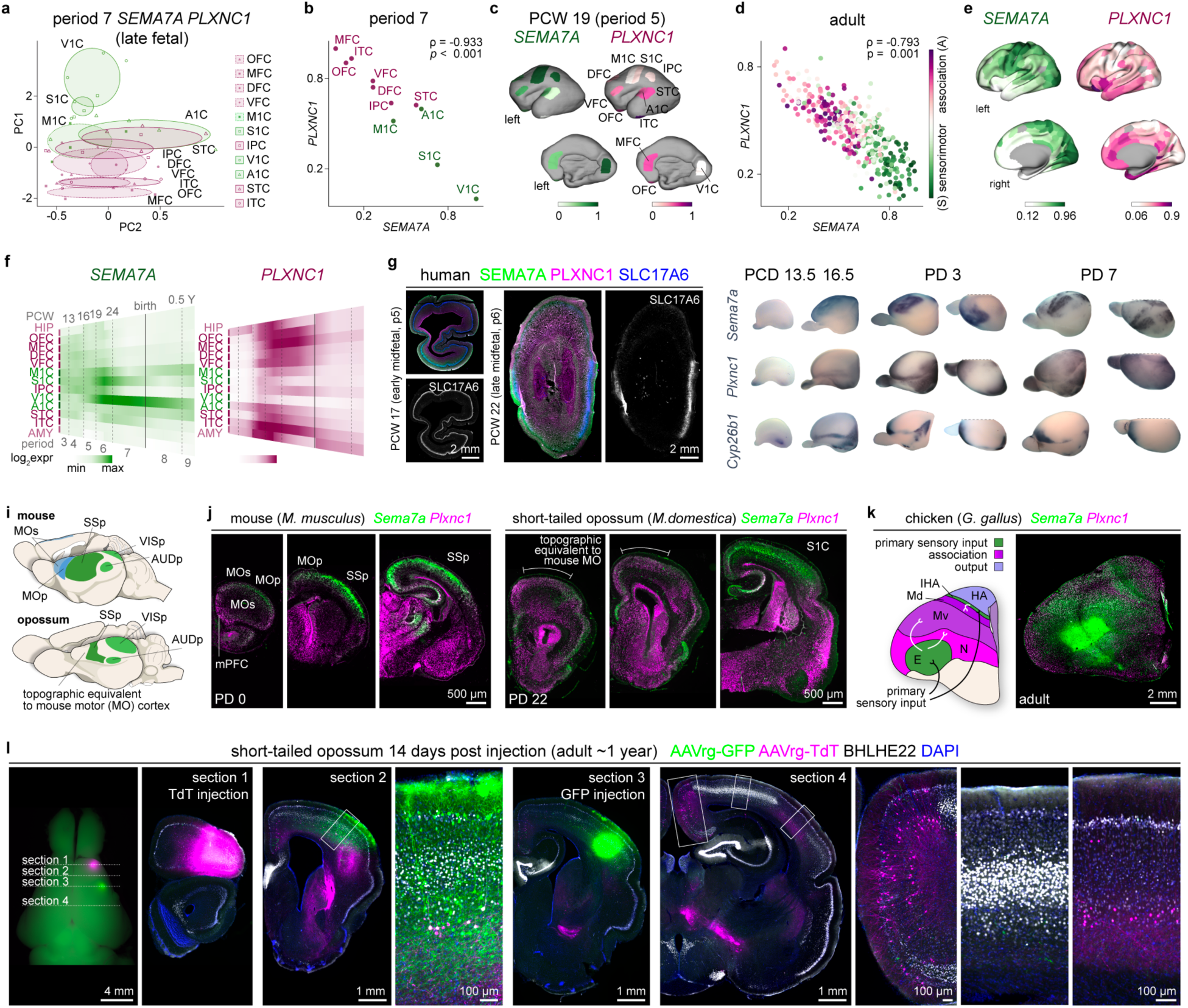
Phylogenetic analysis of the S-A axis reveals conserved and divergent features. **a,** PCA using combined *SEMA7A* and *PLXNC1* expression at period (p) 7. **b,** Scatterplot showing the association between *SEMA7A* and *PLXNC1* expression for p7. Regions are colored according to predefined S-A axis definitions shown in Fig. 1a. **c,** MRI-based cortical surface renderings of a PCW 19 (p5) fetal brain showing gene expression of *SEMA7A* and *PLXNC1* in the sampled neocortical regions. **d,** Scatterplot showing the association between *SEMA7A* and *PLXNC1* expression for adult. Regions are colored according to their position along the archetypal S-A axis of brain organization^3^. **e,** Cortical surface renderings showing gene expression of *SEMA7A* and *PLXNC1* respectively in adult. **f,** Heatmaps showing minimum to maximum median values of individual mean expression for *SEMA7A* or *PLXNC1* in 13 different regions from p3-9. **g,** Human PCW 17 and 22 occipital brain sections stained for SEMA7A, PLXNC1, and SLC17A6. **h,** Whole mount in-situ hybridization for *Sema7a*, *Plxnc1*, and *Cyp26b1* at PCD 13.5, and 16.5, and PD 3, and 7. *Cyp26b1* is initially enriched in the amygdala and spreads to mark the insula and other allocortical regions surrounding the neocortex and motor cortices. **i,** Cortical area map in mouse and short-tailed opossum. **j,** RNA-scope in mouse and opossum at PD 0 or 22, respectively, for *Sema7a* and *Plxnc1*. **k,** Diagram of brain areas and labels in a coronal chicken brain slice adapted from Briscoe et al., 2018^42^ alongside an adult chicken brain section subject to RNA-scope for *Sema7a* and *Plxnc1*. HA, hyperpallium apicale; IHA, interstitial nucleus of the hyperpallium apicale; Md, dorsal mesopallium; Mv, ventral mesopallium; N, nidopallium; E, enteropallium; VT, ventral telencephalon. **l,** Left: image of opossum brain two weeks post-injection of AAVrg-TdT and AAVrg-GFP. Dotted lines indicate the approximate anterior-posterior position from which the coronal images with corresponding labels were obtained.

### The concept of pericentral and central gene programs

To visualize the spatiotemporal progression of association and sensorimotor GMs, we generated heatmaps showing shared or individual dataset GM expression across the analyzed regions of the neocortex and allocortex (paleo- and archi-cortex) throughout pre-natal and post-natal development (**Fig. 1b and Extended Data Figs. 1a,2b,3b**). The spatiotemporal pattern of the shared GM genes is consistent across each of the three datasets we analyzed (**Extended Data Figs. 2 and 3**). When we use GMs generated independently from each of the three datasets, we observed similar spatiotemporal patterns (**Extended Data Fig. 4**). Based on spatiotemporal expression patterns during p3-7, we first observe the emergence of the Af and At modules in prospective F-T association areas adjacent to the allocortex. These modules then progress toward the IPC, which is located centrally in the developing neocortical sheet, while largely avoiding surrounding prospective primary sensorimotor areas associated with the enrichment of S GMs (**Fig. 1b and Extended data Figs. 1,2b,3b,4**). Conversely, S GM expression becomes enriched in the prospective late fetal primary sensorimotor areas (**Fig. 1b and Extended data Figs. 1,2b,3b,4**). Notably, the Af and At GMs share transcriptomic features with HIP and AMY, respectively, suggesting molecular and developmental connections between association neocortex and the allocortex, which may explain their tendency to be highly interconnected^1,6,9^ (**Fig. 1b and Extended data Figs. 1,2b,3b,4**).

Building on this, we propose that S and A GMs represent parts of larger conserved molecular programs regulated by distinct mechanisms that facilitate the patterning of primary sensorimotor and association areas that we define as ‘central’ and ‘pericentral’ programs, respectively. Central, relating to the central location of primary sensorimotor areas along the F-T axis, and pericentral, relating to the surrounding frontally or temporally located, mainly transmodal and paralimbic association areas. V1C is located at what is often considered the posterior, or occipital, pole of the adult neocortex, however, with respect to an F-T axis, it forms in a central region of the prenatal neocortex, in-between the parietal and temporal association regions. The F-T transmodal regions occupy more peripheral, or pericentral, locations aligning with the true topographic poles, the frontal septal and temporal amygdalar poles. The distinctive sets of expressed genes, previously implicated in neural development and autism, and their differing spatiotemporal patterns between pericentral and central programs suggest that transmodal cortices may be under distinct patterning influences from those of primary sensorimotor areas.

### Late fetal F-T to S-A axis molecular shift

To investigate how S and A GM genes distinguish different topographic and functional axes across cortical development, we performed principal component analysis (PCA) of gene expression using the combined genes in the shared S and A GMs. During early fetal development (p3), PCA revealed a polarized organization of gene expression variability aligning with an F-T axis of cortical topography, resembling an adult-like organization with prospective primary sensorimotor areas positioned centrally and intercalated with the prospective transmodal IPC (**Fig. 1d,e and Extended Data Fig. 2c,d**). We observed similar patterns in the macaque cortex (**Extended Data Fig. 3c,d**).

By late-fetal period (p7), the polarization of gene expression along the F-T axis is less apparent and the overall variability of gene expression across the cortex decreases (**Fig. 1d,e and Extended Data Figs. 2c,d and 3c,d**). Beginning in p7 and continuing postnatally, gene expression becomes increasingly polarized along the emerging S-A axis (**Fig. 1d and Extended Data Fig. 2b**), This suggests that, after mid-fetal development, the dominant spatial patterns of gene expression variability align more closely with the functional S-A axis than neocortical topography. Notably, V1C, which represents the dominant sensory modality in primates, consistently shows the most distinctive expression profile along the PC corresponding to the S-A axis (**Fig. 1d and Extended Data Figs. 2c,3c**).

As expected from our analysis of differentially expressed genes in prospective sensorimotor and association areas, we found a significant negative correlation between principal component (PC) scores derived independently from S and A GMs for most developing periods (p3: r = −0.77, *p* = 0.006; p4: r = −0.37, *p* = 0.257; p 5: r = −0.89, *p* = 0.001; p6: r = −0.78, *p* = 0.004; p 7: r = −0.66, *p* = 0.032), indicating that they define opposing anti-correlated expression gradients (i.e., genes in the S GMs are preferentially expressed in prospective sensorimotor while A GM in association areas; **Fig. 1f,g and Extended Data Fig. 4a**). We found a similar negative association between PC scores of the two GMs using whole-brain adult gene expression data^32^ (r = −0.88, *p* = 0.001) (**Fig. 1g**). These findings support the conclusion that gene expression gradually becomes polarized along the emerging functional S-A axis during the mid-to-late fetal transition, ultimately coming to resemble the S-A axis observed in mature brains. These findings suggest that certain genes in pericentral and central modules are the drivers of the archetypal S-A axis development^1–3^.

### The induction-exclusion principle

Our analytical approach identified groups of spatiotemporally regulated transcriptional modules emerging from the F-T poles (A GMs) or as distinct foci within the central region of the F-T neocortical axis (S GMs). We interpreted this pattern as evidence of two pericentral programs emerging from F-T nodes (**Fig. 1h**). We hypothesize that as these pericentral programs advanced toward the neocortical center, transcriptional signatures specific to unimodal primary areas are induced, predominantly within prospective layer 4, the principal thalamo-recipient layer. These signatures emerge within the central neocortex, distinguishing primary areas from the surrounding prospective association cortex by the exclusion of pericentral programs. We interpret the refinement of central programs as a result of processes driven by FO modality-specific thalamic nuclei, each functioning as nodes of induction for primary sensorimotor areas (**Fig. 1h**).

The opposing expression patterns of the S and A GMs suggest antagonistic interactions between prospective primary sensorimotor and association areas. The independent PCA of genes within the S and A GMs respectively, conducted separately across developmental and adult stages described above, revealed genes with complementary functions between pericentral and central GMs. A representative complementary gene pair with high loading PC1 scores is *SEMA7A* and *PLXNC1*. *SEMA7A* exhibited a progressively increasing PC1 loading score relative to other S genes through development, ultimately reaching the highest value in the adult S GM (**Extended Data Fig. 8a**). *SEMA7A* encodes a membrane-bound ligand that binds to the receptor encoded by *PLXNC1*, which also shows a progressively higher PC1 loading score in Af and At GMs with age. Both trajectories align with the increasing refinement of the S–A axis with age^3^. This receptor-ligand pair mediates axonal repulsion through bi-directional signaling and activity-dependent synaptogenesis and dendritogenesis^33–37^.

We hypothesize that the opposing spatio-temporal patterns of S and A GMs, especially the genes encoding axon guidance and cell-cell adhesion genes, facilitate the modular organization of cortical networks and strategic avoidance of primary sensorimotor areas by distributed long-range association projections (**Fig. 1a,b and Extended Data Figs. 1-4**). Consistent with this hypothesis, we observed that key pericentral GM axon guidance and cell– cell adhesion genes are expressed by neurons in both the earlier-generated deep layers and the later-generated upper layers of the F-T (para)limbic and transmodal cortices (**Extended Data Fig. 7**). However, this pattern progressively shifts toward more exclusive expression in deep layers as one approaches the unimodal sensorimotor areas. Previous studies in primates have shown that the laminar origin of fronto-temporally aligned long-range cortico-cortical projections to the PFC shifts from deep to superficial layers when moving from (para)limbic and transmodal areas toward unimodal association areas^6,7,9^. Based on this, we investigated the presence and orientation trajectories of early cortico-cortical axon fibers in the early to mid-fetal human cortex. Using diffusion-weighted imaging (DWI) on ex vivo post-mortem human fetal brains and applying procedures to enrich for putative intra-hemispheric cortico-cortical streamlines (**Methods**), we identified emerging streamlines oriented along the F-T axis in prospective transmodal and para-limbic association areas as early as post-conception week (PCW) 13, corresponding to developmental period p3. These streamlines emerged in parallel with more localized streamlines in prospective primary unimodal areas (**Extended Data Fig. 9**), suggesting that the earliest transmodal streamlines are polarized along the F-T axis and avoid putative primary unimodal regions. These findings complement the spatiotemporal articulation of pericentral transcriptional programs along the F-T axis during the same developmental period.

Given the spatiotemporal progression of central and pericentral programs, and the emergence of primary sensorimotor areas as highly intra-connected regions surrounded by inter-connected association areas^6,7,9^, we propose that the induction of central programs results in the exclusion of pericentral programs in the prospective primary sensorimotor neocortical territories. Specifically, the exclusion of pericentral programs from prospective primary sensorimotor areas, induced by first-order thalamic inputs and activity-dependent mechanisms, results in the archetypical neocortical organizational motif of isolated unimodal islands surrounded by an ocean of associative cortex. We propose that the induction-exclusion principle, a foundation of the MIND model, represents a generalizable core mechanism facilitating the assembly of distributed, predominantly fronto-temporally aligned, connectivity of the association cortices that regulates the balance of primary sensorimotor to association areas required to establish a species-specific S-A equilibrium. The following sections provide further experimental evidence to support this model.

### SEMA7A and PLXNC1 delineate S-A axis

The high PC1 loading scores of *SEMA7A* and *PLXNC1* (**Extended Data Fig. 8a**) suggests that these genes are exemplary markers of primary sensorimotor and association areas, respectively. We examined the pattern of *SEMA7A* and *PLXNC1* expression from early fetal development to adulthood (**Fig. 2a-e and Extended Data Fig. 10a**). PCA at various developmental periods using *PLXNC1* and *SEMA7A* expression across the sampled neocortical regions revealed that, by p7, PC1 corresponded to the S-A axis (**Fig. 2a and Extended Data Fig. 10c**).

We quantified the relative expression of SEMA7A and PLXNC1 along areas of the S-A axis in both fetal and adult brains. This analysis revealed a consistent pattern: low *SEMA7A* and high *PLXNC1* expression in association areas, and high *SEMA7A* and low *PLXNC1* expression in primary sensorimotor areas (**Fig. 2d,e and Extended Data Fig. 10a**). These findings indicate that *SEMA7A* and *PLXNC1* serve as robust markers for primary sensorimotor and association areas, respectively, across both developmental and adult stages of the S-A axis.

Next, we examined the spatiotemporal expression patterns of *SEMA7A* and *PLXNC1*, which correspond well to S and A GMs (**Fig. 2f**). As expected, *SEMA7A* (sensorimotor-enriched gene) and *PLXNC1* (association-enriched gene) were significantly negatively correlated in the adult dataset^32^ (r = −0.79, *p* = 0.001, **Fig. 2d**) and across most development periods (p3: r = −0.63, *p* = 0.073; p4: r = −0.80, *p* = 0.011; p5: r = −0.90, *p* = 0.001; p6: r = −0.93, *p* < 0.001; p7: r = −0.93, *p* < 0.001) (**Fig. 2b**). Notably, the expression of *SEMA7A* becomes refined and increased in V1C between the early mid-fetal and late mid-fetal periods (p5 and 6) (**Fig. 2f**), which we confirmed with staining for SEMA7A, PLXNC1 and the TCA enriched SLC17A6 in human occipital cortex (**Fig. 2g**) and in situ hybridization of *SEMA7A* and *PLXNC1* in macaque cortex (**Extended Data Fig. 10b**).

To investigate the developmental progression of central and pericentral programs, we performed whole mount in situ hybridization at various stages of mouse brain development using *Sema7a*, and *Plxnc1* as representative markers for these programs. We also examined expression of *Cyp26b1*, which encodes an all-trans retinoic acid (RA) degrading enzyme induced by thalamic input.^38^ This gene is enriched in the anterolateral motor cortex and exhibits a ring-like expression pattern in surrounding (peri)-allocortical structures, including the indusium griseum, posterior orbital, anterior insular, perirhinal, and entorhinal cortices, as well as the hippocampus and amygdala^26,27,38–41^. CYP26B1 restricts RA signaling to the fetal PFC, indicating a potential link between RA and pericentral program influence^39^. From post-conception day (PCD) 13.5 to postnatal day (PD) 7, *Plxnc1* expression expands from the F-T poles to the mPFC, insula and temporal association area (**Fig. 2h**). By PD 3, *Plxnc1* expands to the agranular retrosplenial cortex medio-dorsally as well as to central territories surrounding the emerging primary areas (**Fig. 2h**). *Sema7a* expression is initially low but broad across the neocortex at PCD 16.5. Over successive developmental stages, its expression becomes more restricted, upregulated, and refined, predominantly localizing to the primary sensorimotor area, which represents the dominant sensory modality in rodents, and to a lesser extent, the primary auditory and visual areas (**Fig. 2h**). These findings illustrate the progressive localized refinement of central programs along with the inward spread of pericentral programs from the F-T poles and their exclusion from the central regions. Consistent with primate data, the emergence of key Af and At GM genes in mice aligns with the topographic frontal and temporal poles, specifically, the anterior ventromedial frontal and posterior ventrolateral temporal cortices, situated near the septum and amygdala, respectively, and adjacent to other allocortical and ventral pallial structures.

### Evolutionary conservation and divergence in the S-A axis

The conserved emergence and spread of key Af and At GM genes led us to examine if the general organizational principals of the S-A axis are conserved between species. Using *SEMA7A* and *PLXNC1* as markers of primary sensorimotor and association areas, respectively, we analyzed the molecular architecture and connectivity of the S-A axis in eutherian (human, monkey, and mouse), marsupial (short-tailed opossum), and avian (chicken) brains (**Fig. 2i-k and Extended Data Fig. 10c-e**). We observed that the modular expression pattern of *Sema7a* nested within a broadly distributed *Plxnc1* signal is conserved across members of these species (**Fig. 2i-k and Extended Data Fig. 10c-e**). In mouse and opossum, *Sema7a* and *Plxnc1* marked known primary sensorimotor and association regions, respectively. Although the avian dorsal pallium lacks the archetypal six-layered neocortex, it contains neuronal populations homologous to mammalian intra-telencephalic ExNs^42,43^. We observed primary sensory input regions expressing *Sema7a* surrounded by *Plxnc1* expressing association and output regions in chicken dorsal pallium (**Fig. 2k and Extended Data Fig. 10e**). Based on previously published connectivity data^42^, we observed that the regions receiving primary sensory input express *Sema7a* and project directly to other *Sema7a-*expressing regions (**Fig. 2k**). This includes the interstitial apical hyperpallium (IHA), the main sensory input region of the hyperpallium^42^, which contains neuronal types transcriptomically homologous to mammalian neocortical thalamo-recipient layer 4 and 5 ExNs^44^. This indicates that the mechanisms underlying the S-A axis compartmentalization of *Sema7a* and *Plxnc1* are evolutionarily conserved and ancient.

Opossums, unlike eutherians, lack a true primary motor cortex in the frontal region adjacent and rostral to the primary somatosensory cortex. Instead, they exhibit a complete overlap, or “amalgam”, of motor and somatosensory representations within the parietal cortex^12,45,46^ (**Fig. 2i**). According to the induction-exclusion principle of the MIND model, this dorsolateral frontal region, devoid of primary motor cortex, should express *Plxnc1* and exhibit distributed connectivity with higher-order and transmodal association areas. This pattern should arise due to the absence of pericentral program exclusion normally enforced by the presence of a primary motor cortex. Consistent with this prediction, the dorsolateral frontal cortex in opossums expresses *Plxnc1*, whereas the topographically equivalent region in mouse expresses *Sema7a* (**Fig. 2j and Extended Data Fig. 10d**).

To assay the connectivity features of this region, we utilized viral retrograde tracing to label neuronal inputs. We injected retrograde TdT-expressing adeno associated virus (AAVrg-TdT) into the unknown dorsolateral frontal region, and retrograde GFP-expressing AAV (AAVrg-GFP) into the opossum somatosensory cortex, which is predicted to have extensive local connections within somatosensory cortex. We examined TdT and GFP signal two weeks post-surgery (**Fig. 2l**). GFP⁺ cells were confined to the primary somatosensory cortex, as marked by BHLHE22, whereas TdT⁺ cells displayed a broad distribution across what appears to be the association cortex, spanning multiple anterior-posterior positions between the primary sensory domains defined by strong expression of BHLHE22 (**Fig. 2l**). Examining the molecular and connectional features of the dorsolateral frontal region in opossum represents a natural experiment testing the hypothesis that, in the absence of the primary motor cortex, the resulting topographically equivalent region would be overtaken by expanded *Plxnc1* expression and demonstrate association-like connectivity features.

Taken together, the complementary and nested expression patterns of SEMA7A and PLXNC1, representative of central and pericentral gene expression programs, appear to be a conserved feature from avians to primates.

### Evolutionary divergence in the birth order of thalamic nuclei

While the nested expression of central and pericentral programs appears to be a conserved feature across mammals, and possibly birds, there is remarkable diversity between species in the size of primary sensorimotor and association cortices^12^. We propose that a range of intrinsic and extrinsic factors regulate the emergence and progression of the central and pericentral programs. We therefore sought to determine mechanisms which regulate the ratio of primary sensorimotor versus association areas (i.e., S-A equilibrium) across mammals, to uncover why certain species, like primates, have neocortices dominated by association areas.

FO TCAs exert instructive influences through activity-dependent interactions with immature neocortical neurons to drive the formation of primary areas^14,15,18,19,23^. Another driver of S-A axis development may be spatiotemporal gradients of neurogenesis. These gradients have consequential effects on the development of brain structure. For instance, large-brained mammals have exaggerated temporal patterns of neurogenesis compared to rodents^47^. We therefore examined spatiotemporal trends of neurogenesis in the cortex and thalamus in mice, which have a neocortex dominated by primary sensorimotor areas, and in macaques, which have a neocortex dominated by association areas.

We analyzed the birth order of several FO and higher-order (HO) thalamic nuclei and primary sensorimotor and association cortical areas comprising the three major sensory modalities and several prominent association areas in mice and macaques. To determine birth order, which we based on the time of peak neurogenesis in a particular thalamic nucleus, or cortical layer 6, in mice, we used the Neurobirth resource^48^, a database of quantitative 5-ethynyl-2’-deoxyuridine (EdU) birth-dating information spanning the development of the thalamus and cortex. For macaques, we compiled results from previously published tritiated-thymidine birth-dating studies and imaged and quantified tritiated thymidine positive brain sections using archival tissue sections in the MacBrain Resource Center^13,49^ (**Extended Data Fig. 11, Supplemental Table 14**, **and Methods for details**).

We observed that the birth order of thalamic nuclei was not conserved. While FO nuclei were generally born first in mice, the same pattern was not observed in macaques, with the majority HO nuclei born first (**Fig. 3**). Therefore, in addition to an exaggerated sequence of neurogenesis in the thalamus and cortex, FO nuclei emerge relatively later in development in macaques, compared to HO nuclei. The delayed generation of FO nuclei may allow for the prolonged neocortical invasion of pericentral programs and delayed central program induction causing a shift in the S-A equilibrium resulting in a neocortex dominated by association areas. This mechanism based on temporal development of thalamic nuclei, combined with peripheral inputs from sensory organs and neocortical size and geometry^50^ represent generalizable factors regulating S-A equilibria in mammalian clades from rodents to primates.

**Figure 3:**
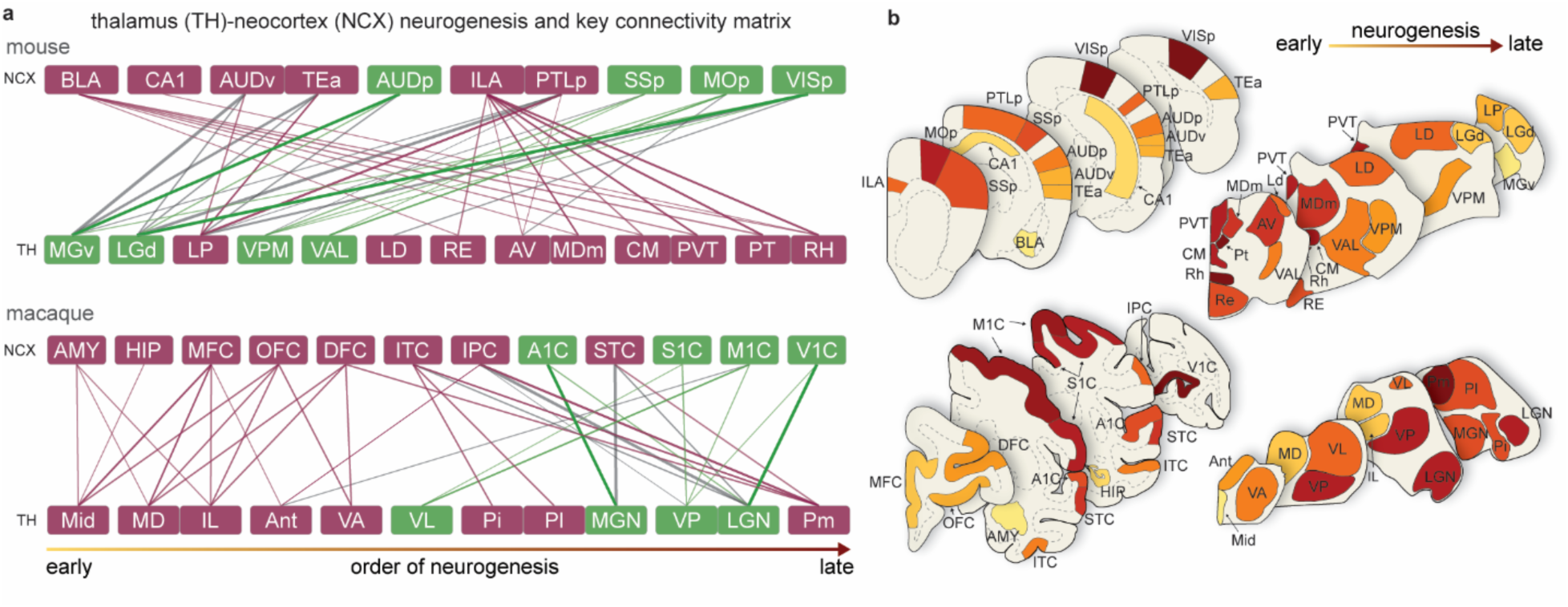
Divergence in thalamic nuclei birth order between mouse and macaque. **a,** Schematic representation of the order of neurogenesis of thalamic nuclei and cortical areas in the mouse (top) and macaque (bottom). Birth order goes from left (early) to right (late). Boxes representing areas that are generated at indistinguishable time-points lack a space. Lines indicate thalamocortical connections and the thickness and opacity are approximately proportional to the connectivity strength. Colors of lines are such that first-order (FO) nuclei to primary sensorimotor area connections are green, higher-order (HO) nuclei to association areas connections are magenta, while the FO nuclei to association cortices or HO to primary area connections are grey. **b,** Representative sections in mouse and macaque with cortical areas or thalamic nuclei colored based on neurogenic order.

### Retinoic acid signaling regulates *PLXNC1* expression

While FO thalamic input drives unimodal/primary sensorimotor cortex development, we do not know how pericentral programs are regulated. Previous studies have demonstrated that RA is enriched in the mid-fetal human frontal and temporal association cortices compared to other centrally located neocortical areas^39^. Moreover, certain genes expressed in our pericentral shared GMs, such as *CBLN2* and *MEIS2*, are enriched in the fetal PFC and regulated by RA^26,27,39,40^, and RA-related GO terms are enriched in the single dataset GMs from Li et al., 2018 (**Extended Data Fig. 4 and Supplementary Table 8**). RA has also been shown to play a critical role in regulating gene expression, neuronal maturation, connectivity, and areal identity of the PFC^38–41^. Collectively, these findings suggest that RA may regulate expression of *PLXNC1,* and other genes enriched in the pericentral GM. To investigate a potential link between RA signaling and *PLXNC1* expression, we immunostained PCD 13.5 mouse brains for PLXNC1 and the RA-synthesizing enzyme ALDH1A3 (**Fig. 4a**). Both proteins were detected in the ventral forebrain and lateral regions of the dorsal pallium (**Fig. 4a**). We also utilized *RARE-lacZ* mice which express *lacZ* under the control of an RA response element^38,39^ and examined lacZ signal at PCD 13.5 and 16.5 in whole-mount preparations (**Fig. 4b**). Activity of lacZ was present in gradients from high in the infero-medial frontal cortex and postero-lateral temporal amygdalohippocampal region to low in the central neocortical territories (**Fig. 4b**). This observation, together with our previous findings, led us to hypothesize that RA signaling contributes to the induction of *PLXNC1* and, more broadly, pericentral programs.

**Figure 4:**
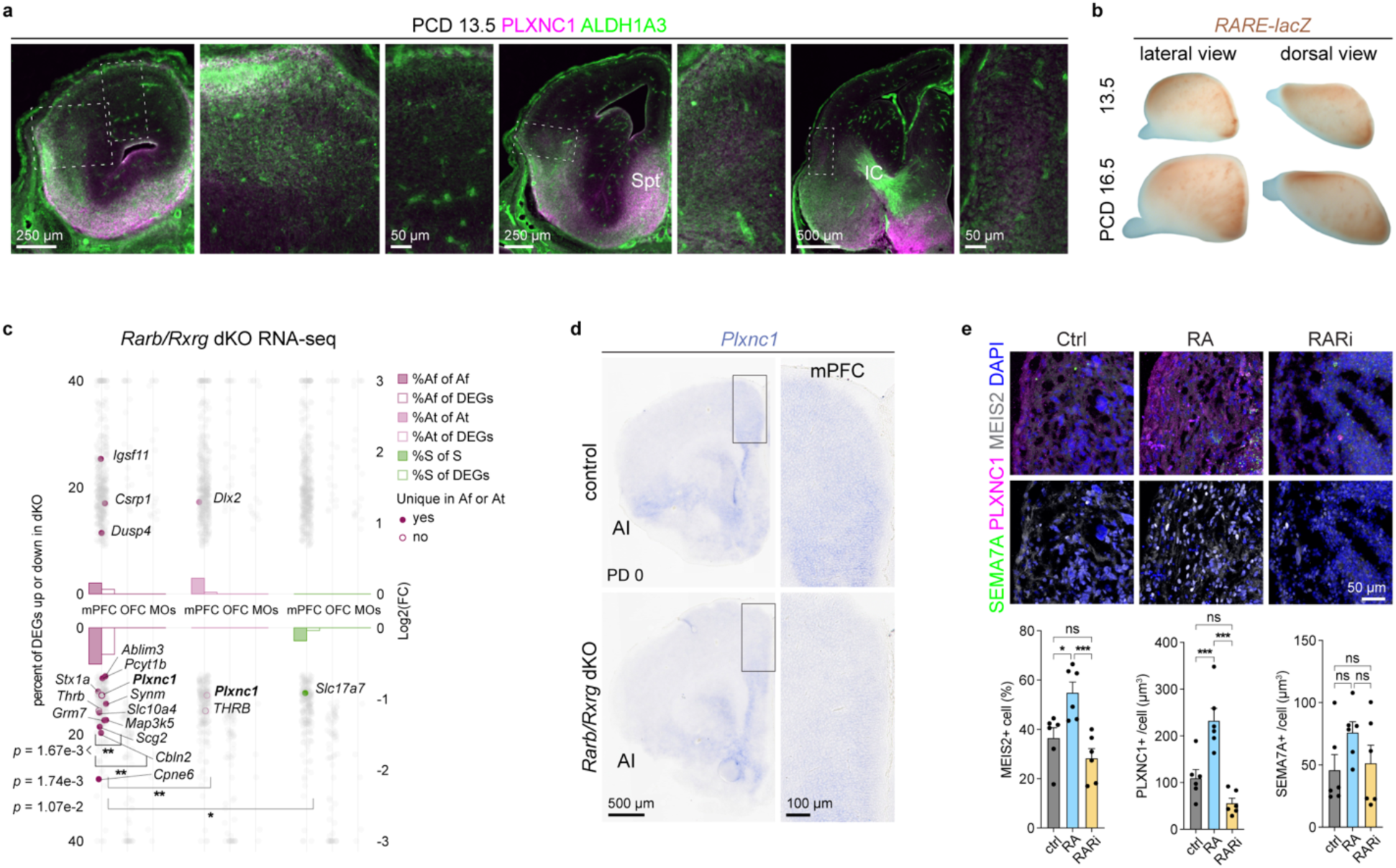
Retinoic acid signaling regulates *PLXNC1* expression. **a,** PCD 13.5 brain sections stained for PLXNC1 and ALDH1A3, revealed co-expression of PLXNC1 and ALDH1A3 in the lateral cortex and the septum (Spt) and high expression of ALDH1A3 in the internal capsule (IC). **b,** *RARE-lacZ* driven lacZ signal in whole-mount preparations of PCD 13.5 and 16.5 brains. **c,** RNA-seq analysis of *Rarb/Rxrg* dKO and wildtype mPFC, OFC, and MOs regions at PD 0. DEGs are labeled that appear in Af, At, or S GMs and the percent of genes represented within a given gene module is shown by solid bars while empty bars represent the percent of all DEGs belonging to a given GM. Fisher’s exact test was applied to test the difference of the percentages both within a GM and between GMs in the same region. *n* = 3 control and *Rarb/Rxrg* dKO. *p*-values indicated in figure. **d,** In-situ hybridization for *Plxnc1* in PD 0 mouse brain sections. AI, agranular insula **e,** Immunostaining for SEMA7A, PLXNC1, and MEIS2 in human telencephalic organoids cultured for 150 days and treated for 48 hours with either RA or a pan RA receptor inhibitor (RARi). The percentage of MEIS2-high cells and the total volume of PLXNC1 and SEMA7A-positive regions were quantified across six organoids (technical replicates). Statistical analysis was performed using ordinary one-way ANOVA with multiple comparisons. MEIS2: ctrl vs. RA, *p* = 0.0161; RA vs. RARi, *p* = 0.0010. PLXNC1: ctrl vs. RA, *p* = 0.0009; RA vs. RARi, <0.0001. For all panels with significance indicated, \**p* < 0.05, \*\**p* ≤ 0.01, \*\*\**p* ≤ 0.001.

To test if *Plxnc1* is regulated by RA signaling, we examined *Rarb*/*Rxrg* double knockout (*Rarb/Rxrg*-dKO) mice, which lack the RA receptors RARB and RXRG and display reduced RA signaling in the mPFC^39^. Using our previously published RNA-seq dataset, we identified differentially expressed genes (DEGs) between *Rarb/Rxrg*-dKO and wildtype mice in frontal areas (mPFC, OFC, and MOs/p) at PD 0 (**Fig. 4c**). Most DEGs overlapping with the shared Af, At, and S GMs belonged to the Af GM, including *Plxnc1,* and were downregulated in the *Rarb/Rxrg*-dKO mPFC (**Fig. 4c**). We performed in situ hybridization for *Plxnc1* in *Rarb/Rxrg* double heterozygous (control) and *Rarb/Rxrg* dKO tissue at PD 0 and found that *Plxnc1* expression was reduced in dKO mPFC (**Fig. 4d**).

To assess whether RA can induce PLXNC1 expression in human cortical neurons, we utilized iPSC-derived cortical organoids which model early cortical development^51^. We generated our own human cortical organoids and also analyzed a publicly available single-cell RNA-seq dataset from human organoids^52^ to examine the expression of *PLXNC1*, *SEMA7A*, and both S and A GM genes. We found that most cells within the organoids corresponded to cortical progenitors and neurons (**Extended Data Fig. 12a**). Notably, *SEMA7A* and select shared S GM genes were expressed by immature ExNs, even after six months in vitro. In contrast, *PLXNC1* and a subset of shared Af and At GM genes were mainly expressed by cortical progenitors, but not by ExNs. This mutually exclusive expression pattern of *SEMA7A* and *PLXNC1* mirrors our in vivo observations (**Extended Data Fig. 12b–d**). These findings suggest that ExNs within the organoids, without the presence of other relevant neural systems, retain a naïve cortical identity. To determine whether RA exposure could induce PLXNC1 in these naïve human cortical organoids, we differentiated the organoids for five months in vitro, allowing the development of major ExN subtypes, and then exposed them for 48 hours to either RA or a pan-RA receptor inhibitor (RARi). We subsequently immunostained control and treated organoids for MEIS2, a protein previously identified as RA-responsive in developing cortical neurons^39^, along with PLXNC1 and SEMA7A. Following RA treatment, MEIS2 and PLXNC1 levels increased, whereas exposure to RARi reduced their expression. SEMA7A levels, however, remained unchanged (**Fig. 4e**). Together, these results indicate that RA regulates PLXNC1 and select genes within the Af GM, at least in part.

### TCA input induces sensorimotor SEMA7A, but not association PLXNC1

To directly test the inductive-exclusionary principle underlying the MIND model, we searched for mouse models exhibiting either absent or topographically altered primary sensorimotor areas, and analyzed the resulting association area changes. We begin by screening mutant mice for defects in primary area development by systematically knocking out *Fezf2*, *Satb2*, and *Zbtb18*, genes known to play key roles in various aspects of cortical ExN projection neuron development^53–59^. To assess changes along the S-A topographical axis, we used SEMA7A and PLXNC1 as markers of primary sensorimotor and association cortices, respectively (**Fig. 5a,b and Extended Data Fig. 14a**). In *Fezf2^flox/flox^*; *Neurod6-Cre* (*Fezf2* cKO) mice, S-A topography appeared largely intact, with a subtle increase in SEMA7A expression in the piriform and motor cortices (**Fig. 5a,b and Extended Data Fig. 14a**). By contrast, in *Satb2^flox/flox^*; *Emx1-Cre* (*Satb2* cKO) and *Zbtb18^flox/flox^*; *Neurod6-Cre* (*Zbtb18* cKO) cortices, SEMA7A was either undetectable or dramatically reduced throughout the neocortex, while the PLXNC1 domain expanded centrally and dorsomedially (**Fig. 5a,b and Extended Data Fig. 14a**). These findings suggest either an expansion of pericentral identity programs within a neocortical context or a fate switch from neocortex to allocortex in the *Satb2* and *Zbtb18* cKO mice. To distinguish between these possibilities, we examined the expression of two genes selectively enriched in allocortical regions by performing RNAscope for *Tfap2d* (paleocortex)^26,60^ and *Zbtb20* (archicortex)^26,61^ (**Fig. 5c and Extended Data Fig. 14b**). While *Zbtb20* expression appeared unchanged, *Tfap2d* exhibited modest expansion in upper layers and decreased expression in deep layers ventrolaterally in both *Satb2* and *Zbtb18* cKOs (**Extended Data Fig. 14c**). Overall, neocortical identity appeared to be preserved in these mutants.

**Figure 5:**
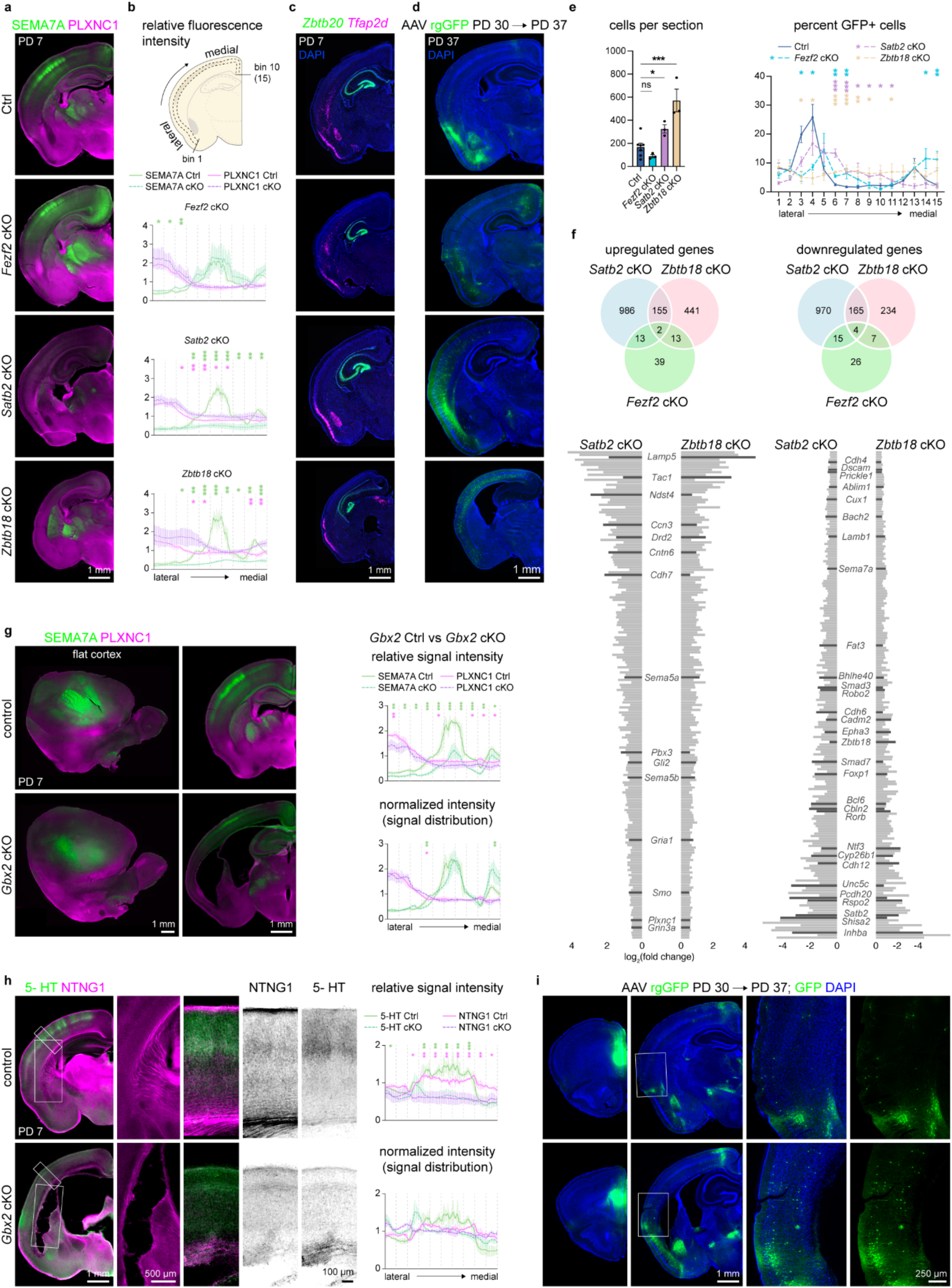
Primary sensorimotor contraction leads to expansion of the transmodal association cortex. **a,** PD 7 control (*Satb2^flox/flox^*), *Fezf2* cKO, *Satb2* cKO, and *Zbtb18* cKO brains stained for SEMA7A and PLXNC1. **b,** Cartoon depicting region quantified (top). Quantifications of signal intensity relative to the mean signal intensity in control versus cKOs. The lateral to medial region quantified was divided into 10 bins, and significance was determined based on the average signal intensity in each bin. **c,** RNA-scope experiments probing for *Zbtb20* and *Tfap2d* in PD 7 control (*Satb2^flox/+^*), *Fezf2* cKO, *Satb2* cKO, and *Zbtb18* cKO brain sections. **d,** GFP signal in sections from PD 37 control, *Fezf2* cKO, *Satb2* cKO, and *Zbtb18* cKO mice in which rgGFP-AAV was injected into the mPFC at PD 30. **e,** quantification of average cells per section in indicated genotype and the percent distribution of cells across 15 equal sized bins from the lateral to medial region of the cortex depicted in panel b. **f,** Venn diagrams representing overlapping and unique DEGs identified from PD 0 *Fezf2*, *Satb2*, and *Zbtb18* cKO neocortical bulk RNA-seq experiments. Bar plots showing shared DEGs between *Satb2* and *Zbtb18* cKO cortices but not in the *Fezf2* cKO. Canonical axon guidance, signaling pathway, area-enriched genes based on Allen brain atlas developing RNA expression data, and other well studied cortical development genes were labeled. All DEGs, genes in Venn diagrams, and bar plots are listed in **Supplementary Table 15**. **g,** PD 7 control (*Gbx2^flox/flox^*), and *Gbx2* cKO (*Gbx2^flox/flox^*; *Olig3-Cre*) brain sections stained for SEMA7A and PLXNC1. The sections imaged in the left column are from dissected and flattened cortices, sectioned tangentially. The right column shows coronal brain sections. **h,** PD 7 control and *Gbx2* cKO brains stained for 5-HT and NTNG1. The relative signal for 5-HT and NTNG1 was determined as in panel **b**, and normalized intensity was determined by dividing each intensity value by the average intensity across the region sampled. For all quantifications, standard error of the mean is shown, and a paired *t*-test was used to determine significance. \**p* < 0.05, \*\**p* < 0.01, \*\*\**p* < 0.001. **See Supplementary Table 16** for statistical information.

To determine if TCAs were reduced or absent, we performed fixed tissue DiI tracing from the thalamus. In *Satb2* cKOs, we observed a loss of DiI signal in L4, though some labeling persisted in L6, likely reflecting cortico-thalamic projection neurons. This was confirmed by cholera toxin B (CtB) injections into the thalamus, which resulted in CtB-positive cell bodies in L6 (**Extended Data Fig. 15a**). In *Zbtb18* cKOs, DiI signal traveled ventrally from the thalamus, away from the internal capsule, and no detectable signal was observed in the neocortex (**Extended Data Fig. 15a**).

We further examined TCAs by staining for 5-HT, a marker of primary sensorimotor TCAs, and the pan-TCA markers NTNG1, and VGLUT2 (**Extended Data Fig. 15b,c**). All three markers were either undetectable or dramatically reduced throughout the neocortex of *Satb2* and *Zbtb18* cKOs cortices, and characteristic barrel fields in somatosensory cortex were not observed (**Extended Data Fig. 15b,c**). Additionally, the central neocortical region topographically corresponding to the prospective primary somatosensory area (SSp or S1C) lacked the archetypal barrel-shaped cytochrome oxidase activity observed in controls (**Extended Data Fig. 15c**). We also found that *Plxnd1* and *Sema3e*, two genes encoding axon guidance molecules required for proper TCA innervation of the cortex^62^, were altered in *Satb2* and *Zbtb18* cKOs cortices (**Extended Data Fig. 15d**). Together, these findings indicate a substantial reduction or complete absence of primary sensorimotor areas, concomitant with the absence of layer 4 TCAs in *Satb2* and *Zbtb18* mutant cortices. Even though we knocked out key neocortical transcription factors, we hypothesize that the observed expansion of PLXNC1 and loss of SEMA7A mainly reflects the failure of pericentral exclusion, which normally results from the FO TCA-dependent induction of SEMA7A and other central identity program genes.

### Sensorimotor contraction leads to transmodal expansion

To determine whether the connectivity of transmodal association cortices is affected by the loss of primary sensorimotor areas, we performed retrograde viral tracing. We injected retrograde adeno-associated virus expressing GFP (rgAAV-GFP) into the mPFC at PD 30 and examined the brains at PD 37 (**Fig. 5d,e and Extended Data Fig. 14d,e**). In *Fezf2* cKO cortices, we observed a subtle medial shift in the distribution of GFP+ neurons, aligning with the increased lateral expression of SEMA7A (**Fig. 5d,e and Extended Data Fig. 14d,e**). Importantly, in *Fezf2* cKO cortices, we did not observe ectopic projections to mPFC from primary sensorimotor areas, despite previously described increases in cortico-cortical projection neurons in *Fezf2* mutants^56^ (**Extended Data Fig. 14d,e**). On the other hand, in *Satb2* and *Zbtb18* cKO mice, there was a dramatic increase in both the overall number of cells projecting to mPFC at multiple anterior/posterior positions of the cortex we examined, and distinct expansions of GFP+ cells medially into the regions typically occupied by primary sensorimotor areas (**Fig. 5d,e and Extended Data Fig. 14d,e**). In posterior regions of the *Satb2* cKOs, the number of cells projecting to mPFC was unchanged, however the domain of GFP+ cells was more dispersed and medially expanded (**Extended Data Fig. 14d,e**).

To further characterize the molecular mechanisms underlying the patterning of the S-A axis, we analyzed shared DEGs between *Satb2* and *Zbtb18* cKO cortices at PD 0 (**Fig. 5f and Supplementary Table 15**). We identified several shared DEGs associated with axon guidance, and cell adhesion, including *Cadm2*, *Cdh4*, *Cdh7*, *Cdh8*, *Cdh12*, *Dscam*, *Epha3*, *Fat3*, *Pcdh20 Plxnc1, Ntf3*, *Robo2*, *Sema5b*, *Sema7a*, and *Cdh8*, some of which were validated by immunostaining and in situ hybridization (**Fig. 5f and Extended Data Fig. 16a**). Additional shared DEGs were related to key developmental signaling pathways, including canonical and non-canonical Wnt signaling (*Prickle1*, *Rspo2*, *Shisa2*), TGFβ (*Inhba*, *Smad3, Smad7*) and sonic hedgehog (*Gli2, Smo*). We also observed that *Cyp26b1,* which is normally enriched in allo-/meso-cortex and motor areas was downregulated in both mice, further supporting the conclusion that neocortex identity is maintained in these mutants (**Fig. 5f**).

To further characterize areal changes, we observed a loss of BCL11A enrichment in primary areas and increased expression of MEIS2, an RA-regulated protein normally enriched in association areas in *Satb2* and *Zbtb18* cKO neocortices. In these mutants, MEIS2 expression expanded into central regions of the neocortex that would normally correspond to primary sensorimotor areas (**Extended Data Fig. 16b**). We also stained for NRP2 (enriched in Kang et al., 2011^27^ Af and At modules), a protein involved in mediating semaphorin and plexin interactions and noted that NRP2 expanded medially into the neocortex which was labeled with TLE4 (**Extended Data Fig. 16c**). Finally, we found that *Satb2* and *Zbtb18* were mutually downregulated, suggesting reciprocal regulatory interactions and indicating that their coordinated activity governs molecular programs underlying TCA ingrowth and, directly and indirectly, the patterning of the S-A axis.

To further test the principles of induction and exclusion, we utilized *Pax6^flox/flox^*; *Emx1-Cre* (*Pax6* cKO) mice in which *Pax6* is deleted from the radial glial cell stage which have been shown to have normal topographic locations, but smaller primary sensorimotor areas^25^. Immunostaining for SEMA7A and PLXNC1 confirmed smaller primary areas and revealed complementary reduced SEMA7A in those areas and expanded surrounding PLXNC1 domain in *Pax6* cKO mice (**Extended Data Fig. 14f**). Consistent with this finding, retrograde tracing indicated increased projections to the mPFC in these mice (**Extended Data Fig. 14f**).

A central principle of cortical arealization is the regulation of primary sensory area development by FO TCAs^12–25^. We attempted to ablate neocortical TCAs through two well-known models, but in both cases observed reduced but not absent TCAs. In *Celsr3^flox/flox^*; *Dlx5/6-Cre* (*Celsr3* cKO) mice^63^, the a-typical protocadherin CELSR3 is absent in the ventral forebrain, resulting in a portion of TCAs failing to cross the pallial-subpallial boundary and enter the cortex. At PD 7, we observed NTNG1-expressing axons projecting through the internal capsule, and SEMA7A expression in layer 4 which lacked clearly defined barrels. (**Extended Data Fig. 17a**). The lack of barrels was also reflected by *Rorb*, *Bhlhe22*, and *Bcl11a* expression (**Extended Data Fig. 17b,c**). Due to the perinatal lethality of this model, we injected retrograde AAV into the mPFC at PD 3 to label mPFC connectivity and observed TdT+ cells in similar neocortical positions in control and *Celsr3* cKO mice. The TdT+ cells in the thalamus confirmed the presence of TCAs in these mice (**Extended data Fig. 17d**).

*Gbx2^flox/flox^*; *Olig3-Cre* (*Gbx2* cKO) mice have TCAs which mostly fail to reach the cortex^64^. *Gbx2* cKOs exhibited reduced SEMA7A and modestly increased PLXNC1 signal in the cortex compared to controls (**Fig. 5g**). SEMA7A and PLXNC1 in flattened cortices as well as quantification of the medial-lateral distribution of SEMA7A and PLXNC1 indicated there was a modest contraction of the SEMA7A expressing domain and a subtle expansion in the PLXNC1 expressing domain towards the neocortical center (**Fig. 5g,h**). We observed a similar reduction of NTNG1 and 5-HT signal, but the presence of NTNG1 expressing axons in the internal capsule, and NTNG1 and 5-HT signal in the cortex, indicated a reduction rather than loss of TCAs (**Fig. 5h**). We performed retrograde tracing at PD 30 and examined the brains at PD 37 and noted an increase in cells projecting to mPFC in piriform cortex as well as a medial expansion of GFP+ cells in *Gbx2* cKO mice (**Fig. 5i**). These results suggest that the reduction of TCAs in *Gbx2* cKO cortices led to a reduction in primary sensorimotor areas with a complementary expansion of transmodal connectivity. Neither the *Celsr3* nor *Gbx2* knockout strategies were successful in abolishing TCAs. While in *Gbx2* cKOs, a reduction in the number of TCAs was accompanied by a decline in SEMA7A expression; the SEMA7A expression domain was mostly normal in size with a modest contraction towards the center.

Using genetic manipulations at multiple levels, including cortical ExNs, progenitors, and the thalamus, to abolish or alter primary sensorimotor areas, consistently led to the expansion of transmodal features into territories typically occupied by primary sensorimotor areas. The loss of TCAs observed in *Satb2* and *Zbtb18* cKOs, combined with increased connectivity between transmodal areas, further suggests that higher-order (HO) thalamic nuclei do not play a primary inductive role in patterning association cortices. Together, these findings support the central principle of induction and exclusion within the MIND model.

### SEMA7A and PLXNC1 repulsion mediates patterning of S-A networks

The shared genes underlying the pericentral and central programs contain axon guidance and cell adhesion genes including *CADM2*, *PCDH7*, *SEMA7A*, and *UNC5C* in S GMs, and *PCDH8*, *PCDH10*, *PCDH17*, *PLXNA1*, *PLXNC1*, and *ROBO1*, in A GMs (**Supplementary Table 5**). These genes and others likely regulate the development of S-A network connectivity. We focused our analysis on the receptor-ligand pair SEMA7A and PLXNC1 enriched in primary sensorimotor and association areas, respectively. Their complementary expression patterns and known repulsive interaction suggest that they play roles in segregating association networks from primary sensorimotor areas.

We utilized a *Sema7a*-null (*Sema7a^-^*) allele and designed and made a *Plxnc1* conditional allele (*Plxnc1^flox^*) (**Extended Data Fig. 18 and Methods**) to generate *Sema7a^-/-^*; *Plxnc1^flox/flox^*; *Neurod6-Cre* (*Sema7a/Plxnc1* dKO) and *Sema7a^+/-^; Plxnc1^flox/flox^* (control) mice. These mice were injected retrograde AAV expressing TdT (AAVrg-TdT) into the primary somatosensory cortex (SSp, S1C) and AAVrg-GFP into mPFC at PD 30 and examined the brains at PD 37 (**Fig. 6a**). Projections to mPFC were not obviously affected, likely reflecting the redundancy in the semaphorin/plexin family and other axon guidance molecules. However, we observed increased nearby projections to SSp from primary auditory cortex (AUDp or A1C) and basolateral amygdala, and a loss of projections from the contralateral entorhinal cortex in the dcKO mice (**Fig. 6a**). This ectopic connectivity between primary sensorimotor areas, entorhinal cortex, and amygdala, a limbic region highly interconnected with neocortical association areas, supports the predicted roles of SEMA7A and PLXNC1 in mediating cortico-cortical network segregation along the S-A axis.

**Figure 6:**
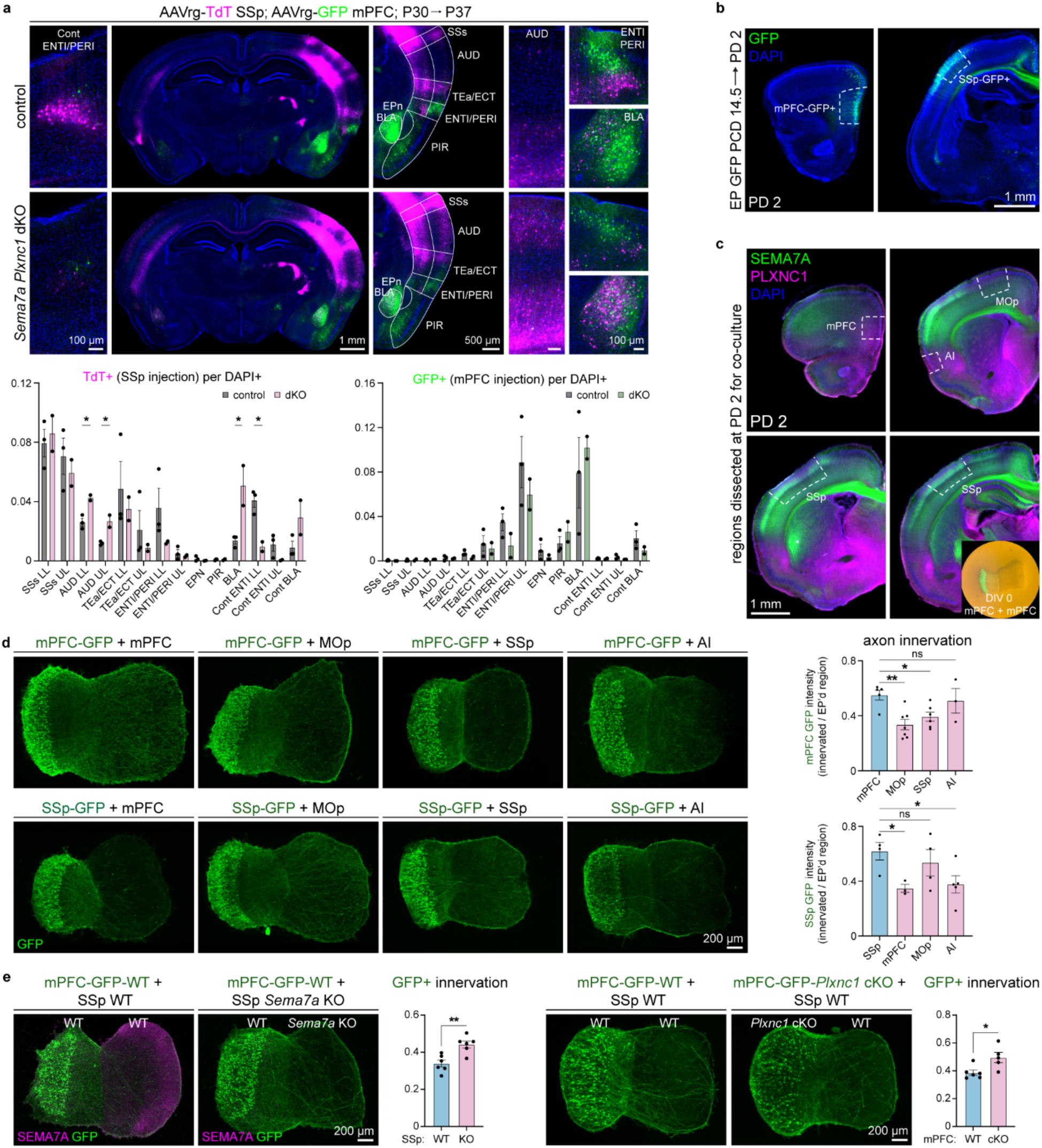
Mutual repulsion of primary and association cortico-cortical connectivity via SEMA7A and PLXNC1. **a,** PD 37 brain sections showing TdT, GFP, and DAPI signal obtained from mice in which AAVrg-TdT and AAVrg-GFP was injected into either the SSp or mPFC, respectively at PD 30. TdT or GFP+ cells per DAPI+ cells were calculated for each region indicated. **b,** Images showing microdissected GFP+ regions for co-culture experiment below. **c,** Images showing micro dissected GFP-regions for the co-culture experiment below and example co-culture at DIV 0. **d,** GFP stained co-cultures generated from PD 2 micro dissected regions and cultured for 2 days in-vitro (DIV), and quantification of GFP+ signal in innervated region relative to GFP+ signal in electroporated region at DIV-2. **e,** Co-cultures generated as described in d, but with either *Sema7a* KO SSp or *Plxnc1* cKO mPFC regions in the indicated combinations. Staining was performed for SEMA7A and shown for cultures involving the *Sema7a* KO experiment. We failed to obtain successful PLXNC1 staining signal in our culturing experiments. For all quantifications, standard error of the mean is shown, and a paired *t*-test was used to determine significance. \**p* < 0.05, \*\**p* < 0.01, \*\*\**p* < 0.001. See **Supplementary Table 16** for detailed statistical information.

### Mutual primary vs association connectivity repulsion

Our model predicts that pericentral and central programs pattern the cortex along an S-A axis through the expression of molecules resulting in mutual repulsion between primary sensorimotor and association areas. We tested this notion by designing an ex vivo culturing strategy in which we challenged the ability of neurons in primary sensorimotor or association areas to innervate different cortical areas (**Fig. 6b-e**). We labeled either mPFC or SSp via in-utero electroporation at PCD 14.5 with *CAG-GFP* plasmids, and co-cultured GFP+ explants at PD 2 with either association areas (i.e., mPFC or agranular insula, AI) or primary areas (i.e., MOp or SSp). When GFP+ mPFC was cultured with mPFC or AI explants, GFP+ axons were readily able to innervate the unlabeled explant, but when mPFC electroporated regions were cultured with MOp or SSp, there was a reduction of GFP+ signal in the unlabeled cortical explants (**Fig. 6d**). Conversely, when electroporated SSp regions were cultured with mPFC or AI, there was a reduction of GFP+ signal compared to when GFP+ SSp was cultured with unlabeled SSp or MOp explants (**Fig. 6d**). These findings suggest that MOp and SSp express molecular signals repelling axons from mPFC, and that mPFC and AI express molecules repelling axons from SSp.

We then utilized this paradigm to test if SEMA7A or PLXNC1 mediate the repulsion between SSp and mPFC. We cultured electroporated wildtype mPFC with either *Sema7a*^+/+^ (*Sema7a* control) or *Sema7a^-/-^* (*Sema7a* KO) SSp (**Fig. 6e**). When GFP+ mPFC was cultured with *Sema7a* KO SSp, we observed increased GFP+ signal in *Sema7a* KO SSp compared to control (**Fig. 6e**). When GFP+ *Plxnc1^flox/flox^*; *Neurod6-Cre* (*Plxnc1* cKO) mPFC explants were cultured with wildtype SSp explants, we observed increased GFP+ signal in wildtype SSp explants compared to those co-cultured with GFP+ *Plxnc1^flox/flox^* (*Plxnc1* control) mPFC (**Fig. 6e**). These findings show that SEMA7A and PLXNC1 can mediate axonal repulsion between mPFC and SSp, which likely reflects their more general role in repulsing axons innervating association and primary areas.

## Discussion

We present experimental findings that integrate with existing knowledge into a unifying MIND model of S-A development. Key molecular and inter-areal connectional features of transmodal and higher-order unimodal association cortices are governed by mechanisms distinct from those implicated in forming primary sensorimotor areas. We term these pericentral identity programs, as they emerge at the neocortical periphery around the septal (frontal) and amygdalar (temporal) poles and gradually propagate inward. As these programs expand towards the center of the naïve neocortex, they interact with focally emerging sensorimotor identity programs. This competitive interplay generates distinct territories through dual processes of induction and exclusion, resulting in early F-T polarization and subsequent spatial compartmentalization of genes previously involved in neural cell differentiation, circuit development, and neurodevelopmental disorders. We demonstrate that the complementary expression of PLXNC1 and SEMA7A, a mutually repulsive axon guidance pair previously implicated in neural circuit and synapse development, across the S-A axis contributes to the antagonistic shaping of cortico-cortical connectivity. The induction-exclusion principles are consistent with certain previous findings^26–28,65–71^.

Due to the gradual inward progression of pericentral programs, centrally emerging sensorimotor programs first define primary territories. As previously shown, these territories then exert activity-dependent influences, anchored in their primary sensoritopic and topological proto-architecture^8,14,15,72,73^, on the surrounding prospective unimodal association cortices, before those regions are further shaped by the later-arriving pericentral programs. This dynamic interplay of antagonistic mechanisms gives rise to a cortical topography in which primary sensorimotor areas emerge as focal “islands” within a broader “ocean” of interconnected association cortex, patterned hierarchically from nearby unimodal association to more distant transmodal distributed networks. This model accounts for the development of transmodal and other higher-order association features regardless of the spatial layout, or even presence, of primary areas. We hypothesize that these developmental processes contribute to specific adult features of areal and laminar connectivity, F-T–polarized transmodal connectivity, gradients, and gyrification, as well as specific evolutionary specializations, exemplified here by our opossum finding. The concept of central programs builds on research emphasizing the sensory-petal activity-dependent, inductive role of FO TCAs in specifying primary areas. Our findings also suggest a mechanistic link between the previously described *Fgf8*- and *Fgf17*-dependent development of the LGE and septum^74^, which are also sources of RA and other factors in the rostral ventral pallium^39,75–78^, and the induction of select Af pericentral genes, such as *Plxnc1*, as well as PFC development. We also elucidated molecular patterning commonalities between transmodal and limbic cortices, helping to reconcile discrepancies observed in cross-transplantation studies between the areal potentials of immature, centrally located presumptive primary sensorimotor cortices and those of peripheral transmodal/limbic cortices^79,80^. Together, our results and model provide a unifying framework for understanding experimental, evolutionary, and clinical phenomena, revealing induction and exclusion as fundamental competitive principles shaping S-A axis and processing hierarchies. The limitations and implications of our findings are discussed in depth in the extended discussion.

## Supporting information

Extended Discussion

## Acknowledgements

The authors thank Suvimal Sindhu for assistance with parcellating the chicken brain and interpreting chicken expression data; Megan Burke for technical support; and Günter Wagner for providing the opossum colony. We are grateful to Ruopeng Wang and Van Wedeen for TrackVis.org (Martinos Center for Biomedical Imaging, Massachusetts General Hospital). We also thank our colleagues at the Yale Genome Editing Center for their help in generating the *Plxnc1-flox* mouse, and the members of the Sestan laboratory for their valuable comments and feedback. Funding for the MacBrain Resource Center was provided by NIH grant MH113257 (to A. Duque). D. Doyle was supported by T32 MH018268. This work was also supported by NIH grants MH129981 (to H. Huang and K. Gobeske), MH094589 (to B. Chen), NS095654, MH106934, MH116488, MH110926, and MH129981, as well as by the Simons Foundation (to N. Sestan).

## Contributions

J.T., M.Z., C.Q., and N. Sestan designed the research. J.T., C.Q., T.F., H.K., S.C., S.B., I.P., S.K., A. Shibata, K.O., N. Salla, E.H., G.S., J.K., C.H., S.L., D.Z.D., N.R-R., and M.S. conducted experimental work. M.Z. and X.L. performed the meta-analysis of primate gene expression data and most bioinformatic analyses, with assistance in initial data processing and analysis steps from S.M. A. Segal generated human brain surface plots, performed PC loading analysis, and created scatterplots. Z.Z. and H.H. contributed DWI data and analyzed it in collaboration with A. Segal. J.T., C.Q., T.F., and O.K. performed retrograde tracing in mice; J.T. and C.H. conducted retrograde tracing in opossum. A. Shibata and G.S. performed immunohistochemistry on human tissue. A. Shibata, J.K., and M.S. completed in situ hybridization experiments. S.K. carried out organoid experiments, including microscopy. J.T., H.K., S.C., and S.L. designed and performed co-culture experiments. S.B. and A.D. analyzed data for thalamic nuclei and cortical area birth dating. E.H., X.P., and T.N. generated the *Plxnc1* flox mouse. J.T., M.Z., A. Segal, and X.L. created supplementary tables. J.T. analyzed microscopy data (excluding organoids). F.T. and Y.N. provided tissue and mice. F.T., K.G., K.K., T.S., B.C., and N. Sestan obtained funding. T.S., B.C., H.H., and N. Sestan oversaw data generation, quality control, and analyses in their respective groups. J.T. and N. Sestan oversaw data quality control and analysis of the data presented in the manuscript. N. Sestan conceived the study and the model, with critical feedback from J.T. J.T. and N. Sestan wrote the manuscript, and J.T. organized all figures. A. Segal and A.F. contributed to interpretation of bioinformatic and imaging results and provided critical revisions to the manuscript. All other authors read and provided comments on the final version of the manuscript.

## Competing interests

The authors declare the following competing interests: Nenad Sestan is a co-founder, board member, and shareholder of Bexorg.

## Methods

### Association and sensorimotor gene module curation

Independently generated spatiotemporal human brain exon microarray data and RNA-seq datasets were obtained from Kang et al., 2011^81^ (accessed through humanbraintranscriptome.org), and Li et al., 2018^29^ (accessed through BrainSpan.org) and the developing macaque brain RNA-seq data across brain regions were obtained from Zhu et al., 2018^30^ (accessed through PsychENCODE.org). The estimated correspondence between macaque developmental time points and equivalent human developmental periods were reported in Zhu et al., 2018^30^. We split association areas representative of pericentral programs into two groups: association frontal (Af) which included orbital prefrontal cortex (OFC), medial prefrontal cortex (MFC), dorsal/dorsolateral prefrontal cortex (DFC), ventrolateral prefrontal cortex (VFC), and association temporal (At) group which included posterior superior temporal cortex (STC) and inferior temporal cortex (ITC). Sensorimotor groups representative of central programs contained primary motor cortex (M1C), primary somatosensory cortex (S1C), primary auditory cortex (A1C), and primary visual cortex (V1C). Dissected areas during fetal development are depicted in Fig. 1a. When defining our gene modules, we used human samples from period (p) 3 to p7 and macaque samples from predicted p5 to predicted p 7 with at least 2 biological replicates for each predicted period (the earliest macaque sample corresponded to approximately human p4 and there were no biological replicates). For each of the 3 sequencing datasets (Kang et al., 2011, human microarray, Li et al., 2018, human RNA-seq, Zhu et al., 2018, macaque RNA-seq), we conducted differential expression analysis using all samples across 10 regions for each period. For RNAseq differential expression analysis, we kept genes with sufficiently large counts using filterByExpr function from the edgeR^82^ package and conducted trimmed mean of M values (TMM) normalization using the normalizeCounts function from tweeDEseq^83^ package. We then applied RNentropy^84^ to identify genes differentially expressed among the above 10 neocortical regions in each developmental period. The resulting DEGs from RNentropy were further selected using criteria adapted from Shibata et al., 2021^39^ in each period: Briefly, a gene to be considered as an up-regulated differentially expressed gene (DEG) in a certain A/S subgroup, (1) there is at least one region in that subgroup where the gene is significantly upregulated; (2) the gene is not upregulated in any region of the opposing (S/A) groups; (3) the gene is under-expressed in at least 30% of the areas in the opposing (S/A) group; and (4) the gene is not in a module gene list of any of the opposing (S/A) subgroups. For exon microarray data in each period, in order to conduct an equivalent analysis to the RNAseq data, we employed limma^85^ to compare each cortical region within a subgroup independently as treatment to all other cortical regions of the opposing group as replicates of control. Differentially expressed genes (DEGs) were selected with FDR < 0.05 and absolute foldchange (FC) > 1 (the same FC threshold used in RNentropy^86^). We then defined the overexpressed DEGs in a certain S/A subgroup in one period and followed the same criteria as we described for the RNA-seq data. Lastly, we merged the genes across all prenatal periods of each gene module within each dataset independently as the final module gene list respectively. To identify shared genes, we found the overlap of each concatenated module gene lists among 3 datasets.

To visualize the module gene expression in heatmap, the module expression value for each region in each period was the median across all individual sample module expression value from that region in that period, where each individual sample module expression value was the mean value of all the genes from the corresponding module in that sample. To generate a smooth gradient change of the module gene expression across developmental time (log_2_(Days)), we interpolated the module expression values to achieve 1000 data points in total based on those of the adjacent discrete period within each region using the interp function from the akima package^87^. The shape distortion of the heatmap is related to the developmental time.

### Principal component and Euclidean distance plots

Principal Component Analysis (PCA) was performed using the prcomp function in R, using the expression matrix of either the combined shared S and A GMs or combined *SEMA7A* and *PLXNC1* expression. Ellipses for each region were centered on the mean of the points within the region, with their axes sized according to the standard deviations on each component. Euclidean distances between different peripheral and central groups were calculated based on the full module gene list for each developmental period. Smooth average lines were generated with a 95% confidence interval.

### Gene set enrichment analysis for GMs

The Gene set enrichment analysis (GSEA) of each GM was conducted using ClusterProfiler^88^ based on reference databases including Gene Ontology (GO), Kyoto Encyclopedia of Genes and Genomes (KEGG), Reactome, and WikiPathways.

### Human diffusion MRI analysis

Diffusion MRI (dMRI) data were acquired from ex vivo human fetal brains at post-conception weeks (PCW) 13, 15, and 17^89–91^. Cortical masks were generated using ITK-Snap, v 4.0.2^92^. Whole-brain tractography was performed using Diffusion Toolkit with a deterministic tracking algorithm, reconstructing fiber pathways based on dMRI data^93^. To isolate cortico-cortical connections, only streamlines with both endpoints located within the manually delineated cerebral wall mask were retained. For ROI-based analysis, tracts were further filtered to include only those with at least one endpoint in a given cortical ROI. Surface ROIs were converted to volumetric space and slightly dilated to compensate for partial volume effects and minor misalignments between the segmentation and underlying diffusion data. Streamlines were visually inspected and cleaned to remove short and anatomically incorrect fibers. Tractograms were visualized in DSI Studio with brains aligned to a consistent orientation for cross-subject comparison^94^. To complement this, whole-brain cortico-cortical tracts were filtered by length using the anterior-posterior extent (L) of each brain as a reference. Thresholds of 1/4L and 1/2L were applied, in addition to unfiltered tracts (0L), to highlight the developmental progression of long-range association pathways. Tractograms were color-coded by principal diffusion orientation (red: left-right, green: anterior-posterior, blue: superior-interior) and visualized using TrackVis (see acknowledgement).

### PC loading scores

For each of the two spatiotemporally distinct, molecularly defined modules (S GM and A GM), we derived the first principal component of gene expression using Principal Component Analysis (PCA) separately. Then we quantified the relationship between these two GM components, separately for 6 developmental periods (developmental periods 3-7, and adult). This allowed us to explore the structure of variation / dominate axis of variation specific to each module.

For periods 3-7, we used the expression data from Kang et al., 2011. We first computed the mean gene expression over replicate samples for each unique period, region and gene. Gene expression values were then normalized column-wise using min-max normalization to ensure comparability across genes. Then using the shared GM gene list (**Supplementary Table 5**), we extracted a 11-region x 43-gene matrix and 11-region x 171-gene matrix for the S GM and A GM respectively.

For adult, we used the Allen Human Brain Atlas (AHBA) dataset. AHBA gene expression data was obtained using the *abagen* toolbox with the default parameters^95–97^. The AHBA is an open-access database containing micro-array gene expression data collected from six human postmortem brains. Samples were assigned to brain regions in the Schaefer 400 region, 17 network atlas^98^. Five regions lacked reliable gene expression so were not included in subsequent analysis (Region ID: 59, Label: lh_17Networks_LH_SomMotB_Cent_5; 173, lh_17Networks_LH_DefaultB_IPL_1; 252, rh_17Networks_RH_SomMotB_S2_5; 299, rh_17Networks_RH_SalVentAttnA_FrMed_2; 303, rh_17Networks_RH_SalVentAttnA_FrMed_4). The resulting gene expression matrix used was 395 regions by 15632 genes. In this matrix, each cell contained the normalized gene expression level for a given region. Using shared S GM and A GM gene list described above, we then extracted a 395-region x 42-gene matrix and a 395-region x 161-gene matrix for the S GM and A GM respectively. Note, there was no gene expression data available for 11 GM genes in the AHBA dataset (*BOC*, *DCHS2*, *DLX5*, *DSC2*, *GCNT2*, *HCRTR2*, *NEUROG2*, *STC1*, *STK32B*, *UBASH3B*, *WBSCR17*).

For each period (3-7, adult), we then decomposed the region x gene matrix using Principal Component Analysis (PCA) to identify the first principal component (PC 1) separately for each GM. Each component is represented by a set of PC scores, one per brain region, defining the dominate axes of variation in gene expression, and a set of PC loadings that capture how strongly expression of particular genes contribute to a component. Note, principal component scores and loadings were sign-aligned for interpretability. Specifically, if the correlation between module PC1 scores and the average expression of the module was negative, we multiplied both the PC1 scores and loadings by −1. This step ensured that positive module PC1s consistently reflected relative gene enrichment within each module.

To evaluate the relationship between the modules, we used Spearman’s rank correlation between the S GM PC 1 and the A GM PC 1. For the fetal data (periods 3-7), at each period, we evaluated statistical significance using a null model where we randomly shuffled gene expression values across the 11 regions for S GM, computed the correlation between the permuted S module and the observed A GM 10,000 times, and then compared to the observed correlation between the S GM and A GM. For the AHBA dataset, we evaluated statistical significance using a spin-permutated Vasa null model (10,000 repetitions)^99,100^.

### Correlation of *SEMA7A* and *PLXNC1* expression in fetal and adult brain regions

We quantified the relationship between *SEMA7A* (sensorimotor-enriched gene) and *PLXNC1* (association-enriched gene). For p3-7, we extracted region-wise normalized gene expression data for *SEMA7A* and *PLXNC1* for each of the 11 cortical regions using Kang et al., 2011. For adult,^32^ we extracted expression data for the marker genes for 395 cortical regions. For each period, we then conducted separate PCAs for each gene and used Spearman’s rank correlation and tested for statistical significance between the PC1 for each of the two marker genes.

### Disease gene enrichment analysis

We conducted a hypergeometric test to examine if the indicated modules were enriched for genes associated with various diseases. We analyzed gene lists associated with neurodegenerative, neurodevelopmental, or psychiatric diseases including Alzheimer’s, ADHD, anorexia nervosa, autism spectrum disorder, bipolar disorder, developmental delay, major depressive disorder, neuroticism, and Parkinson’s disease as well as intelligence quotient described in Mato-Blanco et al., 2024^101^. We also tested if GMs were enriched for diseases that are probably not associated with central nervous system development using genes identified in various genomic analysis and genome wide association studies for coronary artery disease^102^, Crohn’s disease^103^, Lupus^104^, Metabolic syndrome^105^, and Type 2 diabetes^106^.

### Postmortem human and macaque brain tissue

Postmortem human brain samples were collected at PCW 17 and PCW 22. Parental or next of kin consent and approval by institutional review boards was obtained before tissue was collected. Rhesus macaque brain samples were collected postmortem at PCD 165. Whole slabs or whole hemispheres were post-fixed in 4% paraformaldehyde (PFA) for 48 hours and then cryoprotected in an ascending gradient of sucrose 10%, 20%, 30% for one week at each time-point. Tissue was handled in accordance with ethical guidelines and regulations for the research use of human brain tissue set forth by the NIH (http://bioethics.od.nih.gov/humantissue.html) and the World Medical Association Declaration of Helsinki (https://www.wma.net/policies-post/wma-declaration-of-helsinki/).

All experiments using macaque were carried out in accordance with protocols approved by Yale University’s Committee on Animal Research and NIH guidelines.

### Human Tissue Immunohistochemistry

Fixed frozen sections were equilibrated to room temperature (RT) and washed with 1xPBS for 10 minutes. Sections were additionally fixed with 1.6% paraformaldehyde (PFA) for 10 min at RT. To improve tissue adherence, the slides were then baked at 60°C for 30 min and cooled to RT. Next, the samples were incubated in acetone for 10 min at RT. After washing with PBS 3 times, the slides were incubated with a sodium citrate buffer (10 mM citric acid monohydrate, 0.05% Tween-20, pH 6.0) and brought to a boil in a microwave followed by cooling to RT. After washing, the slides were transferred to a slide holder and incubated in autofluorescence quenching buffer (2.25% H_2_O_2_ & 10 mM NaOH in PBS) for 90 min at 4°C. A high intensity broad-spectrum LED light source was placed over the slide holder during this time for additional photobleaching. After washing in PBS, the slides were blocked in blocking buffer containing 5% donkey serum, 1% BSA diluted in staining buffer (2.5 mM EDTA pH 8.0, 0.5x PBS, 0.25% BSA, 0.01% NaN_3_, 0.122 M Na_2_HPO_4_, 0.078 M NaH_2_PO_4_ in ddH_2_O) for 45 min at RT. Next, primary antibodies diluted in blocking buffer was added to the slides and incubated overnight at 4°C. Slides were then washed with staining buffer and post-fixed with 1.6% PFA at RT for 10 min. After washing with 1xPBS, slides were incubated in ice-cold methanol at 4°C for 5 min. Slides were washed with PBST (0.1% Tween-20 in PBS) and secondary antibodies (Jackson Labs) were added (1:500 dilution, diluted in 5% donkey serum, 1% BSA in PBST) and incubated at RT for 2 hours. Next, DAPI nuclear stain was applied to the slides for 10 min at RT. After two washes with PBST and then PBS, the sections were mounted with Fluoromount-G (SouthernBiotech 0100-01). The following antibodies were used, hPLXNC1 (1:250, MAB 544232, R&D Systems), hSEMA7A (1:250, AF2068, R&D Systems), SLC17A6 (1:400, abcam, ab305254).

### Retrograde tracing in the short-tailed opossum

Opossum surgeries were carried out in sterile conditions following all United States Department of Agriculture requirements (USDA) for animals under their oversight, and all procedures were approved by the Yale Institutional Animal Care and Use Committee. Opossums were anesthetized using institutional protocols. Body temperature, heart rate, oxygen saturation, and respiration were closely monitored during the procedure. Anesthetized opossums were placed on a small animal stereotaxic instrument (Kopf, model 940), with a rat head-holder attachment (model 929-B rat gas anesthesia head holder with model 955 ear bars). The skull was exposed, and craniotomies were made using a dental drill above the prospective dorsolateral frontal cortical area and somatosensory area. In the dorsolateral frontal area, a beveled glass needle was inserted 1 mm below the pial surface of the brain. To label all cortical layers, three injections of retrograde AAV carrying p*CAG-tdTomato* (Addgene, 59462-AAVrg) were performed of 50 nl each at 1 mm, 0.6 mm, and 0.3 mm below the surface of the brain. The needle was left at 3 mm for 5 minutes before retracting. To label the somatosensory area, three injections of retrograde AAV carrying p*CAG-EGFP* (37825-AAVrg) were performed of 75 nl each at 1 mm, 0.6 mm, and 0.3 mm below the surface of the brain. The needle was left at 3 mm for 5 minutes before retracting. Two weeks later, opossums were euthanized and subjected to transcardial perfusion with about 200 ml of 1xDPBS followed by 4% PFA in 1xDPBS. Brains were post-fixed in 4% PFA overnight at 4°C and then switched to 30% sucrose 1xDPBS and left at 4°C until equilibrated (brains sank). Brains were sectioned at 60 µm using a Leica cryostat and staining and imaging was performed as described in **Immunohistochemistry using mouse and opossum tissue**.

### Birthdating of thalamic nuclei and cortical areas in the mouse brain

All data on the order of neurogenesis of mouse cortical and thalamic neurons was extrapolated from the collection of EdU birthdating experiments accessible at neurobirth.org (Baumann et al., 2023)^107^. We downloaded the processed quantification of EdU-positive cells within our cortical and thalamic regions of interest. Specifically, the following cortical areas were selected: basolateral amygdala (BLA), as representing the major nucleus of the amygdaloid complex; hippocampal CA1 (CA1) was chosen as representative of the hippocampus; infralimbic area (ILA) for mPFC; primary motor cortex (MOp); primary somatosensory cortex (SSp); primary auditory cortex (AUDp); primary visual cortex (VISp); ventral auditory area (AUDv); temporal association area (TEa); posterior parietal association area (PTLp). These areas were considered the closest correlates of the following human cortical areas analyzed in this work: AMY, HIP, MFC, M1C, S1C, A1C, V1C, STC, ITC, IPC, respectively.

From the same dataset, the following thalamic nuclei projecting to the selected cortical areas were considered: ventral medial geniculate nucleus (MGv), dorsal lateral geniculate (LGd), lateral posterior (LP), ventral postero medial (VPM), ventral anterior lateral (VAL), lateral dorsal (LD), Reuniens (RE), anterior ventral (AV), mediodorsal, medial subdivision (MDm), centromedial (CM), paraventricular PVT, paratenial (PT), rhomboid (RH).

To determine the temporal order of genesis of neocortical areas, we compared the peak of neurogenesis of layer 6a, as it is one of the earliest generated in the mammalian brain^108,109^ and data were available for all areas of our interest. To directly compare the time of neurogenesis of neocortical areas with the genesis of hippocampus, which is not organized into six layers, we consider the peak of neuronal production of the earliest and latest generated laminae of CA1, namely stratum orienses (CA1so) and stratum lacunosum molecolare (CA1slm). To birthdate the amygdala, we selected the peak of neurogenesis of ventral, posterior, and anterior subdivisions of BLA (BLAv; BLAa; BLAp, respectively).

### Birthdating of thalamic nuclei and cortical areas in the rhesus macaque brain

To determine the birth order of thalamic nuclei and cortical (neocortical and subcortical/limbic) areas we performed a meta-analysis of previously published works. In cases in which we could not locate published data for a given area, we obtained images using archival brain sections from tritiated thymidine experiments and quantified tritiated thymidine positive cells in the specified areas. The specimens we used to generate new birth dating data are part of the Rakic collection previously used for various studies of neurogenesis in the non-human primate brain^110,109,111–113,13^. A detailed description of the methods used for the original experiments can be found in the original works. Briefly, in the studies cited, timed-pregnant macaques received an intravenous injection of 10 mCi/kg of thymidine-methyl-^3^H (also known as tritiated thymidine, ^3^H-TdR). The offspring were sacrificed postnatally on the specified day, and autoradiography was used to reveal ^3^H-TdR positive signal on nissl-stained (toluidine blue) brain sections. We integrated the results from previous works that utilized this material with our cell counting analysis on the archival samples to determine the peak time of neurogenesis in all cortical and thalamic regions of interest. Archival material is available and can be requested from the MacBrain-Resource (https://medicine.yale.edu/neuroscience/macbrain/). Metadata on the five samples further analyzed in the current work for integration of the datasets are included in Supplementary Table 3. How we determined the birth order of nuclei and areas is described in the following paragraphs. We obtained information about the neurogenesis of the amygdaloid nuclear complex from Kordower et al., 1992^114^, and hippocampal neurogenesis from Rakic and Nowakowski, 1981^112^. We analyzed the data from CA1 to represent the hippocampus. Neocortical areas including MFC (Broadman Area, BA, 24), OFC (BA 11), DFC (BA 46), and V1C (BA 17) were summarized in Rakic 2002 (see Fig.2)^115^. Rakic identified an early and rapid generation of frontal cortices, that is concluded by E54 with upper layer generation. We therefore considered these three neocortical areas as the earliest generated that we examined, after amygdala and hippocampus. In the same study, the Rakic indicated that neurogenesis in V1C started at the same time as the prefrontal areas but progressed slowly and steadily until much later stages of gestation (PCD 90). We considered this timepoint as the end of neocortical neurogenesis in macaque, and we set V1C as the latest generated cortical area, accordingly.

The order of generation of the remaining neocortical areas was analyzed using archival brain sections from ^3^H-TdR injected macaques and integrated to the above timeline. The following areas were examined: A1C (BA 41, 42); M1C (BA 4); S1C (BA 1-3); STC (caudal superior temporal gyrus, STGc); IPC (BA 7a); ITC (BA 36r). VFC (BA 44, 45) was not included as more rostral sections containing this area were not available for these cases. Sections containing these regions were scanned on an Aperio ScanScope HR CS2scanner at 20x magnification. The resulting images were manually analyzed for quality of the tissue, staining, and imaging. Annotation of the cortical areas of interest was done separately for each individual case based on the anatomical regions defined in the Saleem and Logothetis, 2007 macaque brain atlas^116^. For each area of interest, we selected two serial coronal sections for quantification. ROIs of about 2 x 2 mm were drawn for each area, spanning the cortical depth from pial surface to the cortical white matter. Cells positive for tritiated thymidine were identified by the presence of at least 3-5 silver grains (see examples in Extended Data Fig. 7) and counted manually using the point annotation in QuPath software (v0.5.1). Both heavily and lightly labelled cells were included in the positive count. Total amount of cells revealed by Toluidine Blue were counted by the “Cell Detection” tool in QuPath. By PCD 110, cortical neurogenesis is completed and most ^3^H-TdR - positive cells observed at this age are glial cells which were excluded from the analysis. These were clearly distinguished from neurons by their smaller size, and they are often found physically adjacent to neuronal bodies. After manual counting, positive cells were normalized by area and by total amount of cells for each cortical region. For each area of interest, we calculated the normalized ^3^H-TdR+ cell number at each available injection timepoint (Supplemental Table 3). We then assigned the birthdate of a given region by corresponding it with the injection timepoint that resulted in the greatest normalized ^3^H-TdR+ value compared to all other injection timepoints and plotted it in the respective position in our timeline (Fig. 3).

Birthdating of thalamic nuclei in macaque was determined by performing a meta-analysis of previous studies. The work in Spadory et al., 2022^49^ using similar archival material described above detailed the birth order of the following nuclei used in our timeline: Anterior thalamic nuclear group (Ant); infralaminar nuclei (IL), which includes central medial (CM), central lateral (CL), and parafascicular (Pf) nuclei; mediodorsal nucleus (MD); midline thalamic nuclei (Mid), including reuniens (Re), rhomboid (Rh), paratenial (Pt), and paraventricular (PV) nuclei; ventro-anterior nucleus (VA); ventro-lateral nucleus (VL), ventro-posterior nucleus (VP). These data were integrated with previous birthdating studies of primate thalamic nuclei that included Pulvinar (PUL) subdivisions including lateral, medial, and inferior Pulvinar (PULl, PULm, PULi, respectively), and medial geneiculate nucleus (MGN)^117^. Finally, neurogenesis of macaque lateral geniculate nucleus (LGN) was based on findings in Rakic 1977^118^.

Not all cortical or thalamic areas have exact homologous counterparts between rodents and primates. For example, dorsolateral prefrontal cortex (DFC/dlPFC) was only sampled for the macaque as no equivalent exists in the mouse. We detailed the correspondence between mouse and macaque thalamic nuclei and cortical areas in **Supplementary Table 14**.

### Anatomical connectivity between thalamic nuclei and cortical targets in Fig. 3

To generate a graphical representation of the connectivity strength between various thalamic nuclei and cortical areas, we extracted connectivity measures between areas from previously published anterograde viral tracing experiments. The high-resolution mouse anatomical connectome was derived from a voxel-scale model of Allen Mouse Brain Connectivity Atlas (http://connectivity.brain-map.org/)^119,120^. The connectome data is derived from imaging enhanced green fluorescent protein (eGFP)-labelled axonal projections from 428 viral microinjection experiments in wild type C57BL/6J. We extracted the ipsilateral directed normalized connectivity from thalamic nuclei to regions in the mouse cortex (**Supplementary Table 20**).

In Fig 3a, the thalamocortical connections are represented by lines, the thickness and opacity is derived from connectivity strength. Line thickness was derived by converting the thresholded weighted connection values into thicknesses ranging from 0.25 pt to 1 pt with the more heavily weighted values being represented by lines of higher thickness. Additionally, the line opacity was calculated by assigning opacity values based on the thickness of the lines with the thickest lines having highest opacity. The criteria used for opacity assignment were the following: lines of thickness between 0.25 pt and 0.5 pt were assigned opacity of 60%, thicknesses between 0.5 pt and 0.75 pt were assigned opacity of 80% while thicknesses between 0.75 pt and 1 pt were assigned opacity of 100%. Grey lines indicating connections between FO nuclei and association areas as well as connections between HO nuclei and sensorimotor areas were assigned line opacity of 50%. Connections derived by adjusting mouse connectivity data to reflect accurate macaque connectivity were assigned line thickness of 0.5 pt and opacity of 70%. We removed connections from pulvinar to V1C, S1C, and A1C as the corresponding region in mouse is highly divergent compared to macaque. We also removed MGN-ITC, IPC, and V1C connections as there is extensive primate data charting MGN connectivity, and the coarse nature of tracing experiments in mice, likely contributed to false positives. The conclusions are all based on birth order of thalamic nuclei, while the connectivity between brain regions provides a visual aid and does not alter our conclusions.

### Analysis of human cerebral organoid data

Human cerebral cortex organoid (hCO) data were collected from the Broad Institute’s Single Cell Portal (https://singlecell.broadinstitute.org/single_cell/study/SCP1756)^52^. The Excitatory Neurons without expressing *GAD1* were extracted from each age. Module gene lists of Neocortex (Ncx), hippocampus (Hip), striatum (Str) and thalamus (Thal) were defined in Johnson et al., 2009^26^. Amygdala (Amy) module gene list (containing TFAP2D, TBL1X, and NR2F2) were identified based on in house comparison of the human exon data (Kang et al., 2011) and RNAseq data (Li et al., 2018) used in this study with in situ hybridization (ISH) validation from BrainSpan Atlas^121^. Retinoic acid synthesis (RA_syn) module included *ALDH1A1*, *ALDH1A2*, and *ALDH1A3* genes. The module score was calculated using AddModuleScore function from Seurat V5^122^. A given regional (Ncx, Hip or Amy) cell group was defined with positive module score only of the given region but not the other two regions. To compare the pseudo-bulk gene expression across three brain regions, each regional cell group was randomly split into 3 batches, which were then aggregated as the pseudo-bulk replicates for that region using AggregateExpression function from Seurat V5. Each module expression value was calculated using the median of all genes from the corresponding module for each pseudo-bulk replicate. Regional expression difference of selected genes and modules was tested using t_test function from R with FDR threshold set as 0.05.

### Mice used in this study

All studies using mice (*Mus musculus*) and macaques (*Macaca mulatta*) were performed in accordance with protocols approved by Yale University’s Institutional Animal Care and Use Committee and National Institutes of Health (NIH) guidelines. The mice were housed and kept under consistent environmental conditions at 25°C, 56% relative humidity, and kept on a 12-hour light-dark cycle. Food and water were consumed ad libitum. Experimental cohorts comprised of both sexes. The following mouse lines were used: *Celsr3-flox*^63^, *Dlx5/6-Cre*^123^, *Emx1-Cre*^124^, *Fezf2-flox*^125^, *Gbx2-flox*^126^, *Neurod6-Cre*^127^, *Pax6-flox*^128^, *Olig3-Cre*^129^, *Rarb-null*^39^, *Rxrg-null*^39^, *Satb2-flox*^130^, *Sema7a-null*^33^, and *Zbtb18-flox*^131^.

### Generation of *Plxnc1-flox* line

Mice with a conditional Plxnc1 floxed allele were generated using CRISPR/Cas9 mediated gene editing according to previously described methods^132,133^. A floxed allele of exon 1 was created by using the MIT CRISPR tool (http://crispr.mit.edu) to screen potential Cas9 target guide (protospacer) sequences approximately 1kb upstream of the Plexin C1 5’ UTR and in intron 1 on the reverse strand of chromosome 10. sgRNAs incorporating these protospacers were transcribed in vitro using the MEGAShortscript kit (Invitrogen, Waltham, MA, USA), purified with MEGAclear kit (Invitrogen), and eluted using microinjection buffer. The 156-157 bp repair template oligonucleotides (ssODN) containing loxP sites were synthesized by IDT Technologies. The floxed allele was created in two steps: targeting the 5’ loxP site, followed by a generation of breeding and subsequently targeting the 3’ loxP site. sgRNA/Cas9 RNP and the corresponding template oligo were electroporated into C57Bl/6J (JAX) zygotes^133^, after which embryos were transferred to the oviducts of pseudopregnant CD-1 foster females according to standard methods^134^. Founder animals identified by PCR and sequencing of the loxP target sites. These animals were then mated to C57Bl/6J mice to confirm correct targeting and germline transmission of the conditional knockout allele.

Upstream (5’) guide protospacer + PAM sequence: ATTGAGAACTTGCTCCGTTCTGG Intron 1 (3’) guide protospacer + PAM sequence: CAGCCTGAACCAGCGCGAGGGGG 5’ loxP template oligo: AGTTCTGCTCACTATCTAACCCTAGTGACAAAAGCTAACATTTATTGAGAACTTGCTCCGATAACTTCGTATAATGTATGCTATACGAAGTTATGAATTCTGGGCATAGTTCTAAATGCACCATGTAGATTTTGATTCACTTGATGCTTGGGACAAA 3’ loxP template oligo: TGGTAAATAAATGCTATGGTGCAAAAGCCGCTCGCAGACCGCCCAGCCTGAACCAGCGCGATAACTTCGTATAGCATACATTATACGAAGTTATAAAGGGGGCGGGGCTGTGGCTAGCCAAGGGTGGCTGTGGTCTCAACCTGGTGGAGAGAACCC

### Immunohistochemistry using mouse and opossum tissue

For mouse brain staining, we performed transcardial perfusion with 10 ml 1xDPBS, followed by 10 ml 1xDPBS, 4% paraformaldehyde (PFA) and post-fixed in 4% PFA, 1xDPBS at 4°C overnight. Post-fixed brains were then transferred to 30% sucrose 1xDPBS at 4°C until the brains sank in the solution indicating that they were equilibrated and ready for next steps. The equilibrated brains were immersed and frozen in optimal cutting temperature (OCT) compound (Fisher, 23-730-571) and sectioned at 60 µm using a Leica cryostat (CM3050S). Sections were washed with 1xDPBS at room temperature 3 times for 5 minutes (3 x 5 min) each to remove the OCT and were permeabilized by incubating in 1xDPBS 0.6% triton X-100 for 1 hour. Antigen blocking was performed by incubating sections with blocking buffer (1xDPBS, 5% normal donkey serum, 0.3% Triton X-100) at room temperature for 1 hour. Primary antibodies were incubated with the sections at 4°C overnight, followed by 3 x 5 min wash with DPBS, 0.3% TritonX-100. Individual secondary antibodies (Jackson laboratories) all at a dilution of 1:1000 with a 1:10,000 dilution of 4’,6-diamidino-2-phenylindole (DAPI) to label cell nuclei were incubated with sections for 2 hours at room temperature, followed by 3 x 10 min wash with DPBS, 0.3% TritonX-100. Sections were mounted onto Superfrost Plus Microscope Slides (Fisherbrand, 22-037-246) and sealed with Fluoromount-G mounting medium (Invitrogen, 00-4958-02). Antibody dilutions used for immunostaining were as follows: 5-HT (1:500, Immunostar, 20080), ALDH1A3 (1:250, Proteintech, 25167-1-AP), BCL11A (1:1000, Abcam, ab191491), BHLHE22 (1:500, Sigma, HPA064872), GFP (1:2000, Aves Labs, NC9510598), MEIS2 (1:1000, Santa Cruz Biotechnology, sc-81986), PLXNC1 (1:250, R&D Systems, AF5375), NRP2 (1:250, R&D Systems, AF567), NTNG1 (1:100, R&D Systems, AF1166), SEMA7A (1:250, R&D Systems, AF1835), SLC17A6 (1:500, Synaptic Systems, 135404), TD-TOMATO (1:500, Scigen, AB8181), TLE4 (1:100, Santa Cruz Biotechnology, sc-365406).

SEMA7A and PLXNC1 antibodies were raised in sheep, and goat respectively, which precludes them from individually being recognized by secondary antibodies due to the similarity of goat and sheep IgGs. Therefore, to co-stain for SEMA7A and PLXNC1, we conjugated these antibodies to Alexa fluorophores. Using antibody conjugation kits, SEMA7A was conjugated to Alexa 647 (Invitrogen A2186) and PLXNC1 was conjugated to Alexa 568 (Invitrogen A2184). The resulting primary conjugated antibodies were incubated with no changes to the protocol described above, but the secondary antibody step was omitted.

For some experiments, 5-HT signal was visualized using 3,3’-diaminobenzidine (DAB) staining. Primary and secondary antibody incubations were performed with no changes. A secondary HRP-conjugated donkey anti rabbit IgG (1:2000, Jackson Immuno, 711-035-152) antibody was used. For color development, we used the ImmPACT DAB kit (Vector Laboratories, SK-4105). One drop of ImmPACT DAB reagent was added to 1 ml ImmPACT DAB Diluent and mixed by inversion. Sections were then incubated in the mixture on a shaker. Once sufficient color development was observed by eye, the reaction was stopped by washing with 1 x PBS. Control and experimental groups were incubated at the same time (i.e., one batch) and stopped at identical times Unless otherwise stated in the following methods sections, all microscopy images were obtained with a 10x objective on an Olympus VS-200 Slide Scanner.

### Quantification of immunofluorescence data

PD 7 brain section images obtained using a 10x objective on an Olympus VS-200 Slide Scanner were exported from QuPath, downsample 8 and saved as tifs. The resulting images were opened in FIJI and the segmented line tool with a width of 35 pixels was used to select the quantified region indicated. The straighten tool was then used to generate a rectangular image of the resulting selected area which was subsequently split into 200 equally sized bins and the fluorescence intensity of a given channel was measured in each bin. The values were normalized to the average fluorescence value in the control.

### Preparation of flat mouse cortices

To generate sections from flattened cortices, we performed transcardial perfusion with 10 ml 1xDPBS to remove blood from the tissue. The brains were quickly dropped into ice cold 1xDPBS and cut in half in the sagittal plane. The subcortical structures were carefully removed, resulting in the cortex and olfactory bulb. The cortices were then placed between glass microscope slides with 1 mm spacers. These were then submerged in ice cold 4% PFA, 1xDPBS at 4°C overnight. The following day the microscope slides were carefully removed and the resulting flattened cortices were then transferred to 30% sucrose 1xDPBS at 4°C until the brains sank in the solution indicating that they were equilibrated. The flattened cortices were then section at 60 µm using a Leica cryostat (CM3050S).

### Bulk RNA-seq data analysis for *Satb2*, *Zbtb18*, and *Fezf2* conditional knockout analysis

Fastq files from bulk RNA-seq experiments using dissected neocortices from *Satb2^flox/flox^*; *Emx1-Cre* (*Satb2* cKO) and control mice at PD 0 were downloaded from the GEO database (accession no. GSE68912)^135^. Fastq files of bulk RNA-seq of *Fezf2* homozygous whole-body knockout and wild type mice at PD 0 were downloaded from the GEO database (accession no. GSE160202)^136^. The RNA-seq of the neocortical tissue of the *Zbtb18^flox/flox^*; *Neurod6-Cre* (*Zbtb18* cKO) and wild type mice at PD 0 was obtained from Li et al., 2025^131^. FastQC^137^ was performed before and after trimming adapters using Trimmomatic^138^. The alignment was performed using STAR^139^ against the mouse primary genome assembly GRCm39^140^. Reads with low quality or those that were improperly mapped were filtered out using SAMtools^141^. Duplicated reads were removed using Picard^142^. Then, HTSeq^143^ was applied to annotate features using GENCODE M33 mouse gene annotation^140^ and then we calculated the raw read count mapped to each gene across samples. The differential expression analysis was then conducted using DESeq2^144^. Differential expression of transcripts was detected using a false discovery rate (FDR) < 0.05.

### *In-utero* electroporation

Plasmid mixtures were prepared containing 1 µg/μl *pCAG-GFP* (Addgene # 11150) and 0.1% Fast Green (Sigma) was added to see the plasmid mixture during injections. In-utero electroporation was performed on timed-pregnant PCD 14.5 dams. Timed-pregnant dams were anesthetized using institutional protocols. The uterine horns were exposed. Using a beveled pulled glass needle, about 0.5 µl of the plasmid mixture was injected into the lateral ventricle of each embryo. After visually confirming that the lateral ventricles were sufficiently filled with the plasmid mixture, a handheld platinum coated electrode (Tweezertrode, 5 mm, 45-0489, Harvard Apparatus) attached to an electroporator (BTC, ECM830) was used to deliver five 35V electrical pulses at 50 ms each with a 950 ms interval between pulses to each embryo. The positive end of the electrode was positioned to direct the DNA towards the frontal-medial region of the ventricle to label mPFC and the center of the right or left hemisphere of the dorsal pallium to target SSp. The pups were then delivered without intervention at the expected time.

### Retrograde tracing in mice

For PD 30 injections, animals were anesthetized using institutional protocols. Mice were held in place using a Kopf Stereotaxic instrument (Model 940). The skull was exposed and the bregma point was located. The tip of a beveled glass needle attached to a microinjection unit attached to the stereotaxic instrument was positioned directly atop bregma. This position was considered the zero-point. mPFC was targeted by performing a craniotomy above the planned injection site. The injections were performed at the following position relative to bregma: 2.1 mm anterior, 0.3 mm lateral, and 2.4 mm ventral. Due to the heterogeneity of mutant animals, mPFC was targeted based on anatomical landmarks, and only matching injections determined after processing the brains were used for analysis. 50 nl virus was injected into mPFC, and the needle was retracted after 5 minutes. SSp was targeted by positioning the needle 1.46 mm posterior and 3 mm lateral to bregma. To obtain the maximum precision in the depth of the injection for SSp, a craniotomy was performed to expose the cortex, and the beveled glass needle was moved into the relative posterior and lateral position and placed on the cortical surface, this represented the zero point in the dorsal-ventral axis. The needle was then moved to −0.9 mm relative to the cortical surface and 20nl of virus was injected, the needle was then positioned at −0.6 and −0.3 mm for two additional 20 nl injections to cover the entire depth of the cortex. At the final injection point, the needle was left for 5 minutes before retracting. Targeted regions were injected with 50 nl retrograde AAV carrying p*CAG-EGFP* or retrograde AAV carrying p*CAG-tdTomato* (Addgene, 59462-AAVrg). For all experiments, incubation was for 1 week.

After incubation the brains were collected for histology and stained for the indicated antigens. Brain collection and staining was performed as described in Immunohistochemistry Using Mouse Tissue. Three representative sections at each anterior-posterior position were imaged using the extended focus imaging (EFI) software associated with the Olympus VS200 slide scanner. EFI results in images spanning entire sections with all cells in focus, essentially doing the job of a confocal microscope in a sensible timeframe, facilitating cell counting. All retrograde traced sections for all experiments included in the PD 30 injections were imaged using the identical parameters, EFI with the following exposure times, DAPI 26 ms, GFP 56.5 ms, TdT 56.5 ms.

GFP+ cells were quantified using FIJI. Briefly, the cortex was highlighted using the segmented line tool followed by the straighten function. The resulting rectangular image was split into 15 equal sized bins, and GFP+ cells were counted in each bin. For dual tracer injection experiments, bleed through was observed at the injection sites. Therefore, we manually segmented specific cortical and subcortical areas of interest in QuPath using the DAPI channel as guidance. Cells were quantified using positive cell detection in QuPath.

For PD 3 injections, animals were anesthetized on ice for several minutes. During surgery, the pups were maintained under hypothermia and placed on a stereotaxic apparatus. A purified AAV solution (AAV2-*retro-CAG-tdTomato*; 50 nl per site; injection rate: 50 nl/min) was injected into the right mPFC. The injection site was located 1.0 mm anterior and 0.3 mm lateral to bregma, and 0.5 mm ventral from the surface. Experiments were carried out with KDS Legato 130 (KD Scientific) and stereotaxic frame (Muromachi Kikai). The brains were collected at PD 7. The fluorescent images were acquired using fluorescence microscope (BZ-X810, KEYENCE) with 4x and 10x objectives.

### Fixed-tissue DiI tracing

Brains were quickly dissected at PD 7 and dropped into ice cold 4% PFA, 1xDPBS and left to fix overnight at 4°C. The following day the brains were cut in half coronally, exposing the thalamus. DiI crystals were then carefully placed just below the surface, across the exposed thalamic region. The brains were then embedded in 4% low-melt agarose (invitrogen) and incubated at 37°C for one week. The brains were then sectioned at a thickness of 60 µm using a Leica Vibratome and placed in a solution of DAPI in 1xPBS (1:10,000) for 10 minutes at room temperature before switching back to 1xPBS. Sections were placed on glass slides, sealed with Fluoromount-G and all slides were imaged the same day they were sealed.

### Human telencephalic organoid culture

Human induced pluripotent stem cell lines (line HSB311) were authenticated by morphology or genotyping and confirmed to be free of mycoplasma contamination using the MycoAlert Mycoplasma Detection Kit (Lonza). For maintenance of pluripotency, cells were dissociated into single cells using Accutase (Thermo Fisher Scientific, 00-4555-56) and plated at a density of 1 × 10^5^ cells per cm^2^ on Matrigel (BD)-coated 6-well plates (Falcon) in mTeSR1 (STEMCELL Technologies, 85850) supplemented with 5 μM Y27632 and a ROCK inhibitor (Sigma-Aldrich, SCM075). ROCK inhibitor was removed after 24 hours, and cells were cultured for an additional four days before passaging.

Telencephalic organoids were generated using a directed differentiation protocol as previously described^39^. Cells were dissociated into single cells with Accutase (Thermo Fisher Scientific, 00-4555-56) and resuspended in neural induction medium containing 100 nM LDN193189 (STEMCELL Technologies, 72147), 10 μM SB431542 (Selleck Chemicals, S1067), and 2 μM XAV939 (Sigma-Aldrich, X3004-5MG) to achieve dual SMAD and WNT inhibition. Cells were plated at 10,000 cells per well in 96-well v-bottom ultra-low-attachment plate (Sumitomo Bakelite). To promote cell survival and aggregation, 10 μM Y-27632 (Sigma-Aldrich, SCM075) was added for the first 24 hours. After 10 days in stationary culture, organoids were transferred to 6-well ultra-low-attachment plates (Millipore Sigma) and maintained on an orbital shaker at 90 rpm to improve nutrient and gas exchanges. Starting on day 18, organoids were cultured with maturation medium supplemented with 1× CD lipid concentrate (Thermo Fisher Scientific, 11905031), 5 μg ml^−1^ heparin (STEMCELL Technologies, 07980), 20 ng ml^−1^ BDNF (Abcam, 9794), 20 ng ml^−1^ GDNF (R&D Systems, 212-GD), 200 μM cAMP (Sigma-Aldrich, 20–198) and 200 μM ascorbic acid (Sigma-Aldrich, A92902) to support neuronal maturation. On day 148, all-trans RA (Sigma-Aldrich, R2625) or the pan-RA receptor antagonist (RA inhibitor) AGN193109 (Tocris, 5758) was added to the medium for 48 hours prior to sample collection.

For the preparation of organoids for staining, telencephalic organoids were fixed in 4% paraformaldehyde at 4 °C and cryoprotected in 20% sucrose, and embedded in OCT compound (Thermo Fisher Scientific, 23-730-572). Embedded samples were sectioned at 10 µm using a cryostat (Leica, CM3050S). The organoid sections were washed in PBS (3 x 5 min), then incubated in a blocking solution containing 0.5% (v/v) Triton X-100 and 10% (v/v) donkey serum (Jackson ImmunoResearch Laboratories, 017-000-121) in PBS for 2 h. Primary antibodies were added in 10% (v/v) donkey serum and applied at 4 °C overnight. Sections were washed in PBS (3 × 5 min) and incubated with fluorescent secondary antibodies for 2 h at room temperature in 10% (v/v) donkey serum. After a final PBS wash (3 × 5 min), sections were coverslipped with Vectashield mounting medium (Vector Laboratories, H-1000). The primary antibodies used targeted hMEIS2 (1:200, Abcam, ab244267), hPLXNC1 (1:250, MAB 544232, R&D Systems), hSEMA7A (1:250, AF2068, R&D Systems). Images were acquired using an LSM800 confocal microscope (Zeiss) and processed with Zeiss ZEN and ImageJ software. Z-stack images were analyzed using Volocity (v.6.3.1) and Spotfire (v.11.2.0).

### In situ hybridization in brain sections and whole-mount preparations

Whole-mount and section *in situ* hybridization using antisense digoxigenin (DIG)-labelled RNA probes was performed as described previously^39^, except that 5% dextran sulfate was added in the hybridization buffer for whole-mount in situ hybridization. Brains from the embryonic mouse, Rhesus macaque (M.mulatta) and human were fixed in 4% PFA at 4 °C and sectioned at 20 μm and slides were stored in −80 °C until use.

The following commercially available probes (Horizon Discovery) were used: mouse *Cyp26b1* (MMM1013-202798233), *Plxnd1* (MMM1013-202765536), *Sema3e* (MMM1013-202769580), and *Sema7a* (MMM11013-202769010). The *Plxnc1* probe was made from the following primers, forward (fwd): CAGCCAATCAAACCTTGAGCAC, reverse (rev): GTTGTTGAATAGAGGCCCAGTGAC. Template for mouse *Ntf3* was a gift from Dr. John L.R. Rubenstein (University of California, San Francisco, San Francisco, CA). Template for mouse *Sema5b, Cdh8* probes were gifts from Dr. Elizabeth A. Grove (University of Chicago, Chicago, *IL*).

To detect Rhesus macaque *PLXNC1* and *SEMA7A*, DNA fragments containing single exon of each gene (1057, 1043, 718 bp, respectively) were PCR-amplified from the genomic DNA. DNA fragments were subcloned into pCRII vector using TA-cloning kit (Invitrogen, K202020). All plasmids containing cDNAs and genomic DNA fragments were used as templates to generate probes after linearization by restriction enzymes. Primer sets used for PCR amplification were as follows: *PLXNC1* fwd: ATGGAGGTCTCCCGGAGGA, rev: GGCTCTCGGCCGTCTTGAAG. *SEMA7A* fwd: ACAAGGCCCCACTGCAGAA, rev: TTCCCAGCCCCTCCCTTTC. The digoxigenin-labelled probes were synthesized from the linearized templates either by SP6 (NEB, M0207S), T7 (NEB, M0251S) and T3 (NEB, M0378S) RNA polymerases, respectively and RNA labelling mix (Roche, 11277073910) according to the manufacturer’s instructions. Images of brain sections taken using a VS200 microscope (Olympus Microscopy).

Whole-mount in situ hybridization preparations were imaged and color balance was manually adjusted to normalize the background hue across images without altering signal intensity.

### LacZ signal development in whole-mount preparations

Brains were dissected from PCD13.5 and PCD 16.5 *RARE-lacZ* mouse embryos and drop-fixed in 4% paraformaldehyde for 90 min at 4°C, followed by rinsing with 1xPBS. The enzyme activity of b-gal was visualized using Rad-Gal (Sigma-Aldrich, RES1364C) as substrate for chromogenic reaction.

### RNAscope analysis on chicken, opossum, and mouse brain tissue

Brains from the chicken, opossum (Monodelphis. domestica), and mouse were fixed by quickly dissecting and immersing in ice cold 4% PFA, 1xDPBS overnight, followed by equilibration in 30% sucrose, 1xDPBS. The brains were then sectioned at 20 µm using a leica cryostat and placed on microscope slides and allowed to dry overnight at room temperature to enhance tissue adhesion. We used the following probes: chicken *Sema7a* (1276971, ACD) and *Plxnc1* (1276981, ACD); opossum *Sema7a* (1570351, ACD) and *Plxnc1* (1570361, ACD); mouse *Sema7a* (437261, ACD), *Plxnc1* (495481, ACD), *Zbtb20* (837641, ACD), and *Tfap2d* (551631, ACD). RNAscope was performed according to RNAscope Multiplex FL v2 protocol and kit (323270) from ACD (Advanced Cell Diagnostics bio). Briefly, slides were incubated at 60°C for 30 min, then further fixed in 4% PFA, 1xDPBS on ice for 30 min and then dehydrated with graded ethanol immersions (50%, 70%, 100%). To inactivate endogenous peroxidases, slides were incubated with hydrogen peroxide for 10 minutes and washed with distilled water. Protease treatment was performed with protease III for 10 minutes. Target retrieval was performed by immersing slides in boiling RNAscope target retrieval buffer for 5 minutes. The samples were then washed with RNAscope wash buffer and incubated with probes for 2 hours at 40°C. Following probe incubation, the samples were incubated sequentially with RNAscope Multiplex FL v1 Amp1, Amp 2, and Amp 3 for 30 minutes each. The slides were then incubated with horseradish peroxidase (HRP) C1, and then subject to tyramide signal amplification. This was repeated for each additional channel. All slides were imaged using a 10x objective with a VS200 Slide Scanner (Olympus).

### Organotypic slice co-culture

GFP+ regions were labeled via in-utero electroporation as described in In-Utero Electroporation. The brains of PD 2 control (non-electroporated) and GFP+ mice were extracted and immediately transferred to ice-cold (<4 °C), carbogen (95% O_2_:5% CO_2_) saturated artificial cerebrospinal fluid (aCSF, 92 mM NMDG, 20 mM HEPES, 5.5 mM glucose, 30 mM NaHCO_3_, 5 mM sodium L-ascorbate, 2.5 mM KCl, 1.25 mM NaH_2_PO_4_, 2 mM thiourea, 3 mM sodium pyruvate, 5.5 mM urea, 10 mM MgSO_4_, 0.5 mM CaCl_2_-2H_2_O). Coronal 200 µm-thick slices were obtained using a vibratome (Leica, VT1200S) in ice-cold aCSF continuously aerated by carbogen (95% O_2_:5% CO_2_). Region of interests were dissected from coronal brain slices using scissors with tiny blades (FST, 15000-03). Dissected brain areas were transferred to 50 μg/ml rat tail collagen-I (Corning, 354236) coated culture inserts (Falcon, 353090) in a 6-well plate containing 1.5 ml of slice culture medium (Neurobasal medium (Gibco, 21103049), supplemented with 1X B-27 supplement (Gibco, 17504044), 1% (v/v) GlutaMAX (Gibco, 35050061), 5 µg/ml human insulin (Sigma, I9278), 20% (v/v) horse serum (Gibco, 26050088), and 1% (v/v) penicillin-streptomycin (Gibco, 15070063). The orientation of two dissected slices were arranged such that the ventral sides were in contact, while the dorsal sides (upper-layers) were at opposing ends. Organotypic brain slices were cultured at 37°C for 2 days.

For immunofluorescence staining, co-cultured organotypic slices were fixed on days-in-vitro 2 using 4% paraformaldehyde for 20 min at room temperature. Slices were washed in PBS (3 x 10min), and incubated in blocking solution containing 5% (v/v) normal donkey serum (Jackson ImmunoResearch Laboratories, 017-000-021) and 0.3% (v/v) Triton X-100 in PBS at 4°C overnight. Slices were incubated with primary antibodies diluted in blocking solution overnight at 4°C and washed with PBST (0.1% (v/v) Tween-20 in PBS, 3 x 10min). Slices were incubated in the appropriate fluorescent secondary antibodies (all raised in donkey, Jackson ImmunoResearch Laboratories) diluted at 1:1000 in blocking solution at 4°C, overnight, and washed with PBST (3 x 10min). Slices were then counter stained with DAPI for 10 min at room temperature, and finally washed with PBST (3 x 10min). 2 by 2 tile scan images of eight serial optical sections at 5.8-μm intervals over a total depth of 40.8 μm were acquired with an LSM880 confocal microscope (Zeiss). Orthogonal projection images were used for the quantification of axonal innervation. Mean fluorescence intensity of the electroporated and non-electroporated regions were measured in Fiji.

---

**Extended Data Figure 1:**
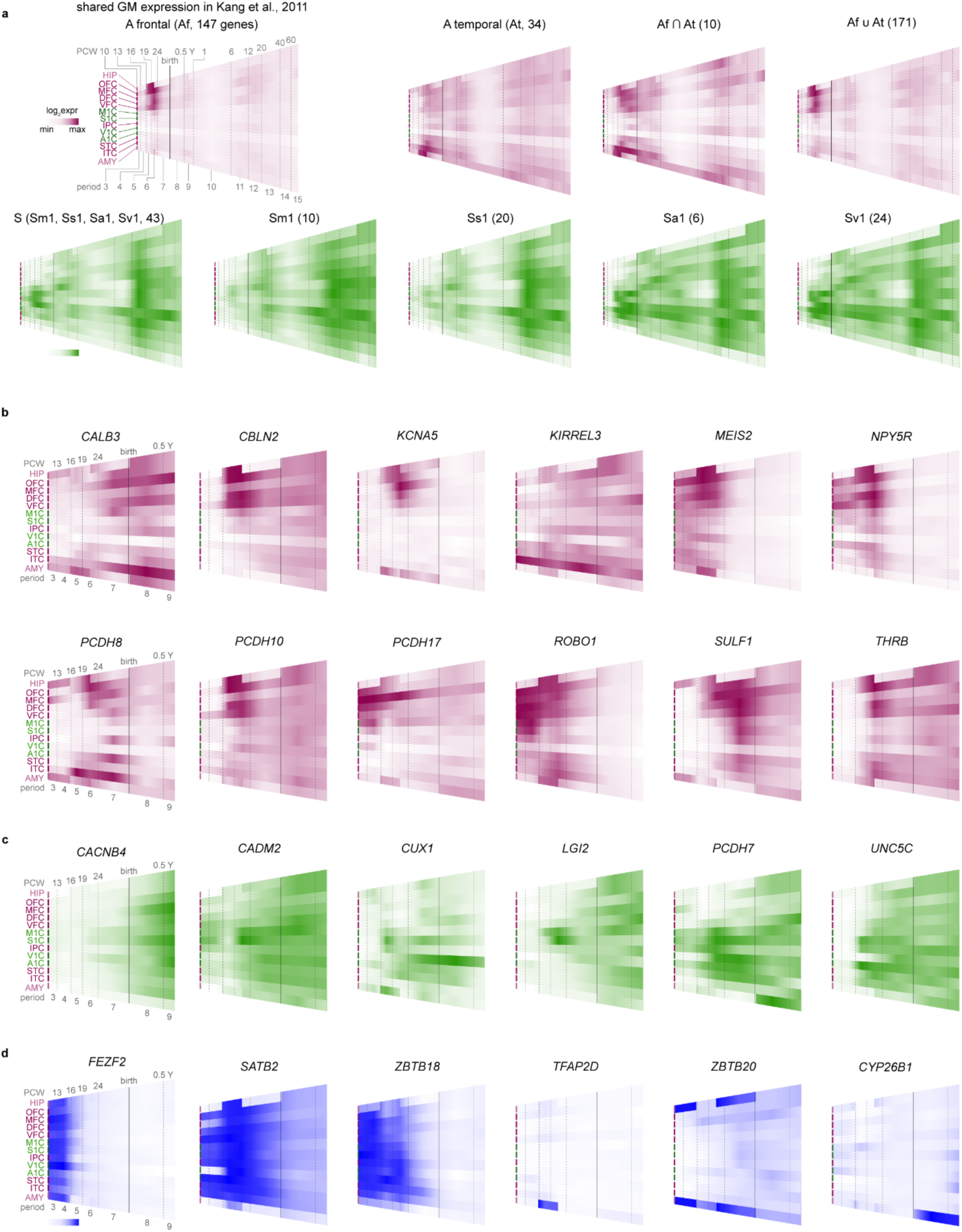
Spatiotemporal progression of GMs and relevant example genes. **a,** Heatmaps generated using the shared GMs. Indicated sampled regions plotted from p3-p15 displaying minimum to maximum median mean value of samples of a given region and period. **b,** Example gene heatmaps from the shared A (Af + At) module. **c,** Example gene heatmaps from shared S (combined Sm1, Ss1, Sa1, Sv1) GM. **d,** Gene heatmaps for frequently referenced genes in our study. All heatmaps in this figure were generated using expression data from Kang et al., 2011.

**Extended Data Figure 2:**
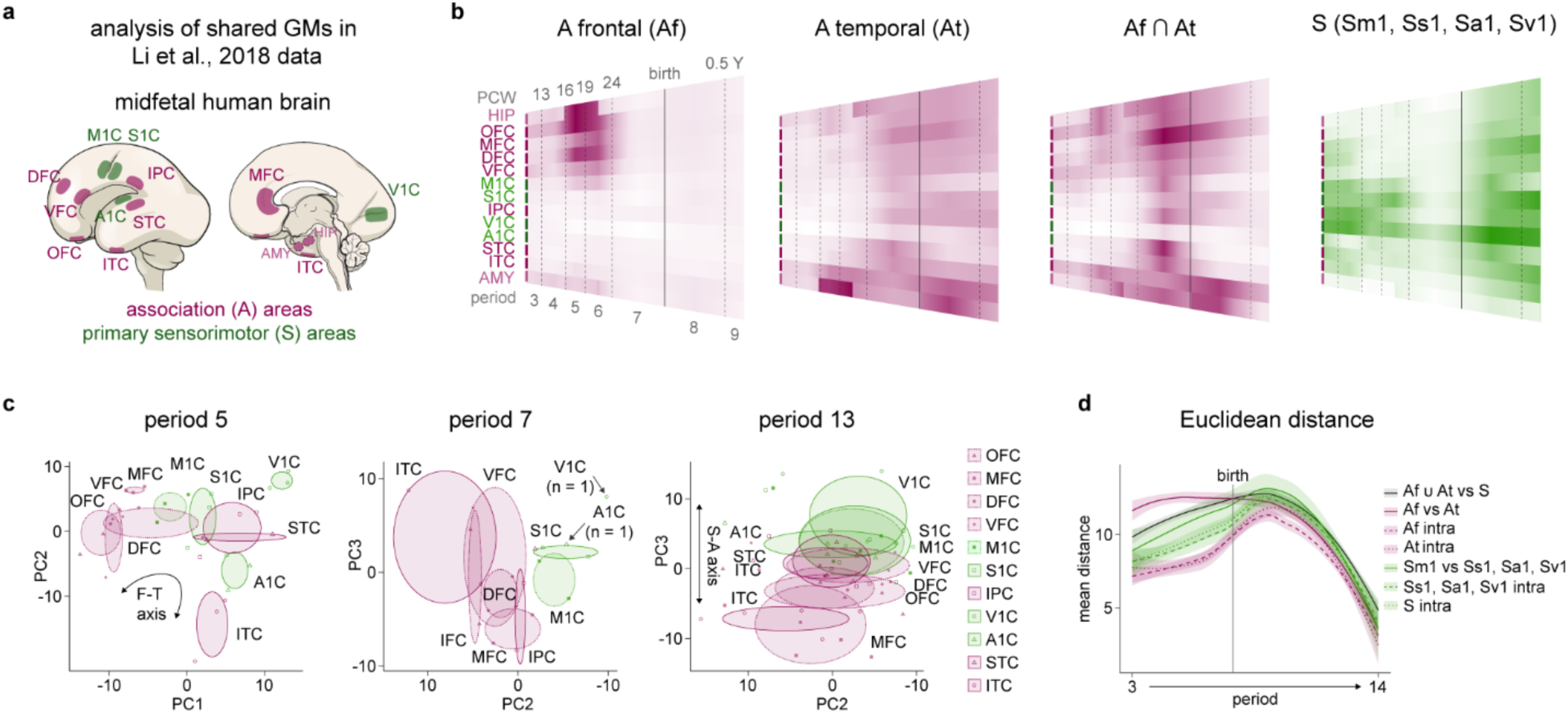
Analysis of shared GMs in Li et al., 2018^29^ human data. **a,** Midfetal human brain schematic with analyzed regions highlighted. **b,** Shared gene module heatmaps with 13 indicated sampled regions across p3-9 displaying minimum to maximum median mean gene value of samples of a given region and period. **c,** Principal component analysis (PCA) plots using combined genes in shared S and A GMs plotted against the indicated principal components. Ellipses are centered on the mean of the points of the indicated region and the size of each axis is determined based on the standard deviation of each component. For period 7 and 13, PC1 appeared to be related to sample quality or another unknown variable and was thus omitted. **d,** Euclidean distance based on all genes in shared S and A GMs between indicated regional groups at each developmental time point. Confidence interval level 0.95.

**Extended Data Figure 3:**
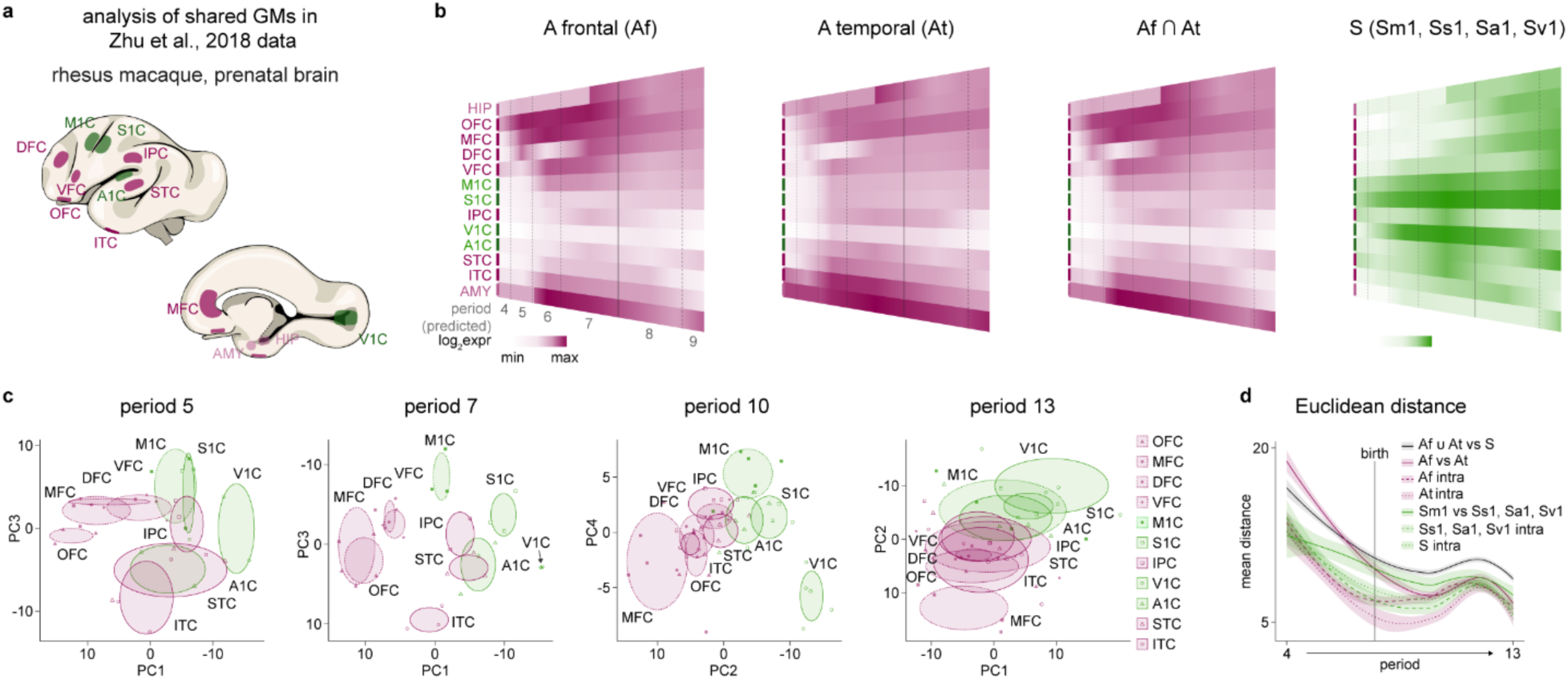
Analysis of shared GMs using Zhu et al., 2018^30^ macaque data. **a,** Depiction of mid-fetal macaque brain with analyzed regions highlighted. **b,** Gene module heatmaps with 13 indicated sampled regions across developmental timepoints estimated to be equivalent to human periods 4-9 showing the median of all individual mean gene expression values in each module for each equivalent period. **c,** Principal component analysis (PCA) with genes in S and A GMs plotted against the indicated principal components. Ellipses are centered on the mean of the points of the indicated region and the size of each axis is determined based on the standard deviation of each component. A p4 PCA plot was not included because only one sample per region was collected at the equivalent p4 stage in macaque. The first two PCs that reflect topographical locations of areas were shown, and those that corresponded to either sample quality or other unknown variables were omitted. **d,** Euclidean distance based on shared genes in shared S and A GMs between indicated reginal groups at each developmental time point. Confidence interval level 0.95.

**Extended Data Figure 4:**
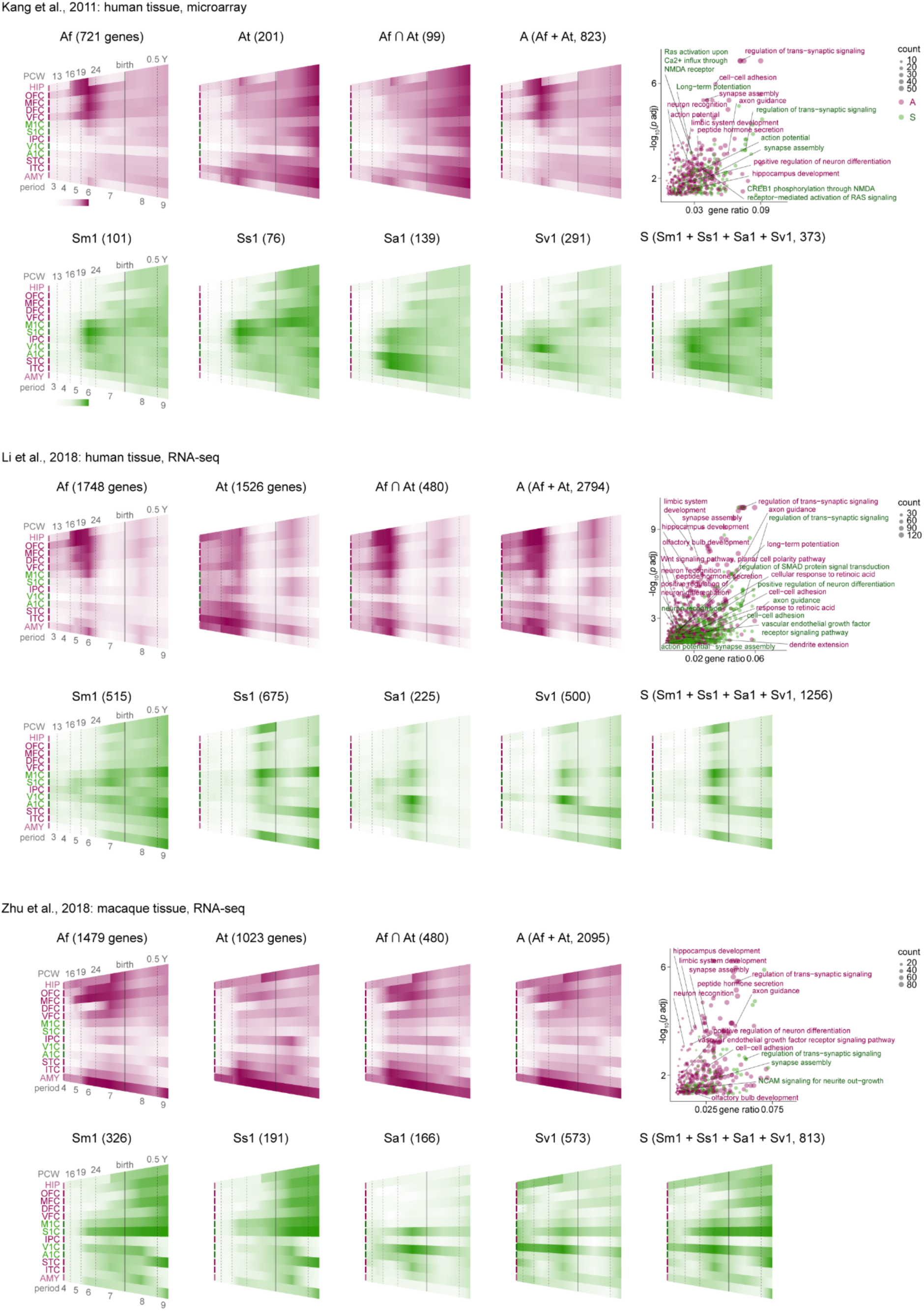
Spatiotemporal expression of GMs generated from three independent datasets. Gene modules were generated using three independent datasets (Kang et al., 2011^27^, Li et al., 2018^29^, Zhu et al., 2018^30^) and the spatiotemporal expression patterns for each dataset-specific set of GMs were shown in heatmaps displaying minimum to maximum median mean value of samples of a given region and period. There were no p3 equivalent samples in macaque (Zhu et al., 2018^30^). The bubble plots are representative of gene set enrichment analysis (GSEA) bubble plot for A (Af, At) and S (Sm1, Ss1, Sa1, Sv1) genes. Complete GSEA in **Supplementary Tables 7-9**.

**Extended Data Figure 5:**
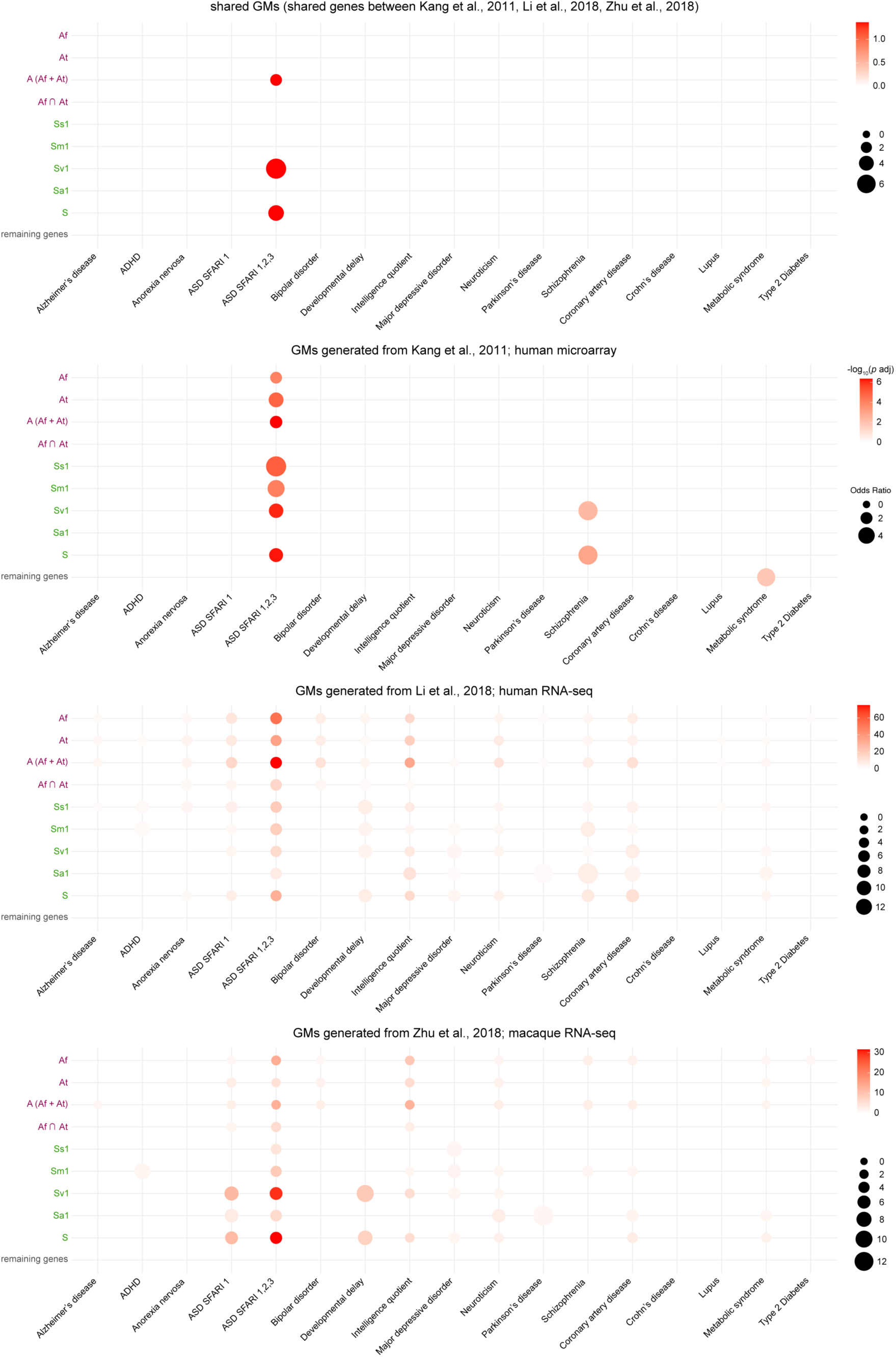
Disease gene enrichment analysis of GMs. Dot plots representative of gene enrichment test for GMs derived from three independent datasets or using shared GMs. A hypergeometric test was used and enrichments with a *p*-value > 0.05 adjusted for multiple comparisons using the Benjamini-Hochberg false discovery rate were considered statistically significant and shown in the plot. We also tested for enrichment associated with disease modules using all remaining genes (i.e., all genes not included in indicated GMs). See **Supplementary Tables 10-13** for genes lists for genes lists in each group (dot). See **Methods** for details regarding disease gene modules. ADHD, Attention-Deficit/Hyperactivity Disorder; ASD Autism spectrum disorder.

**Extended Data Figure 6:**
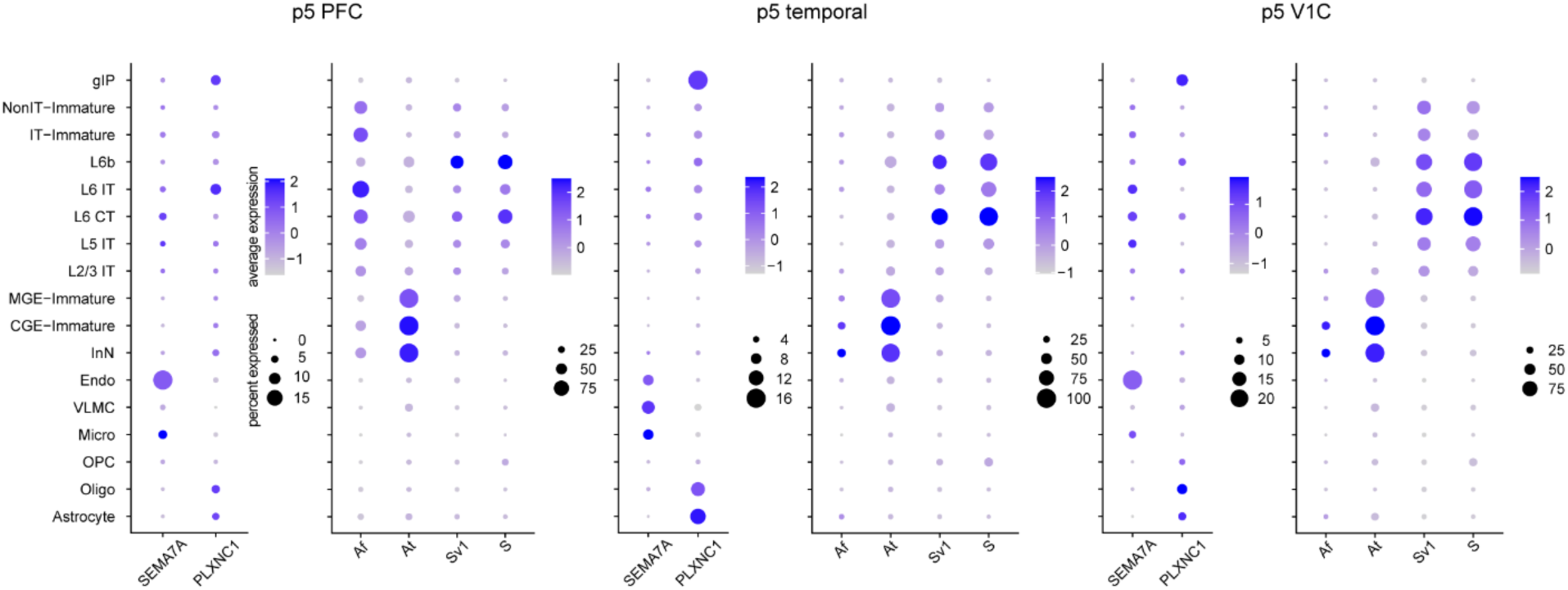
Regional single cell analysis of *SEMA7A*, *PLXNC1* and shared association and sensorimotor GMs. Analysis of cell-type and region-specific expression of *SEMA7A*, *PLXNC1* and shared Af, At, Sv1, and S (Sm1, Ss1, Sa1, Sv1) GMs during mid-fetal (p5) development. Data from Bhaduri et al., 2021^31^. PFC, prefrontal cortex; V1C, primary visual cortex; gIP, glutamatergic intermediate progenitor; IT, intratelencephalic; L, layer; MGE-Immature, medial ganglionic eminence-derived immature interneurons; CGE-Immagure, caudal ganglionic eminence-derived immature interneurons; InN, interneurons; Endo, endothelial cells; Micro, microglia; OPC, oligodendrocyte precursor cells; Oligo, oligodendrocytes.

**Extended Data Figure 7:**
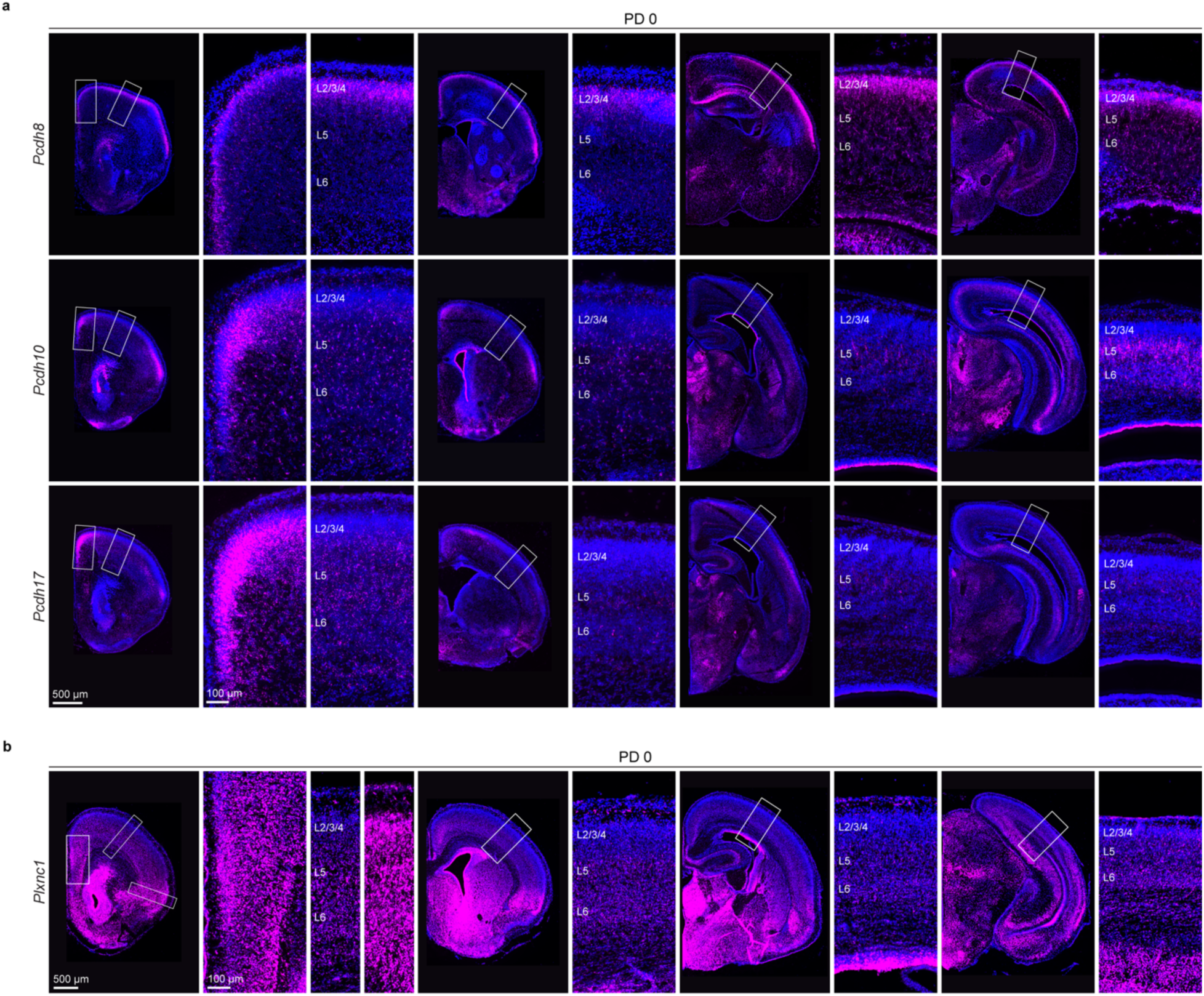
Expression of *Pcdh8*, *Pcdh10*, *Pcdh17*, and *Plxnc1.* **a,** RNA-scope of *Pcdh8, Pcdh10, and Pcdh17* identified in our shared association GMs, in PD 0 cortices. Unlike the other analyzed genes, *Pcdh8*, appears to exhibit non-conserved expression between mice and primates. **b,** RNA-scope probing for *Plxnc1* in PD 0 mouse brains sections. All genes analyzed here are broadly expressed by neurons in both the earlier-generated deep layers and the later-generated upper layers of the developing fronto-temporal (para)limbic and transmodal cortices. For *Pcdh10*, *Pcdh17* and *Plxnc1* this pattern progressively shifts toward more exclusive expression in deep layers as one approaches the unimodal sensorimotor areas.

**Extended Data Figure 8:**
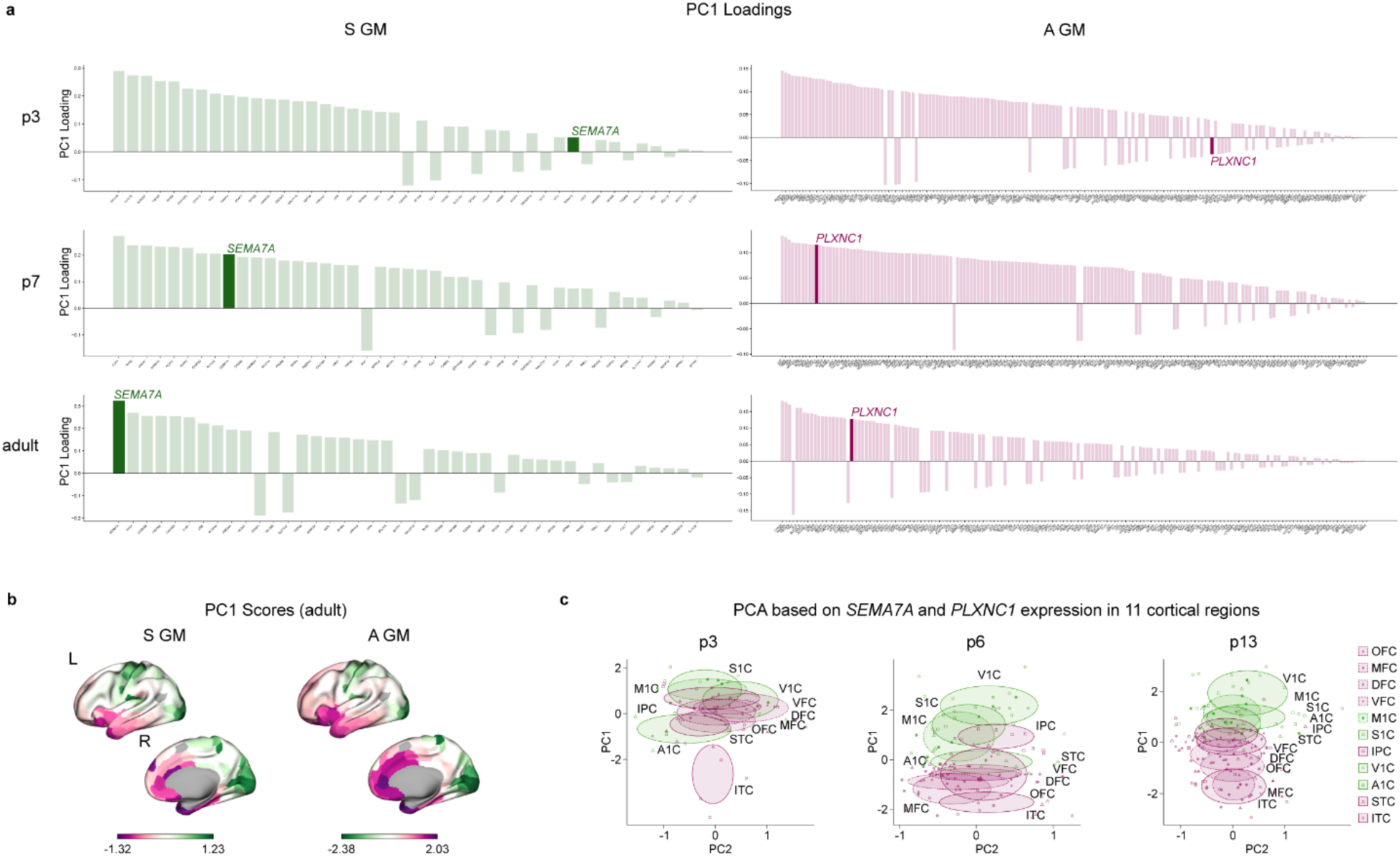
PC1 loadings and SEMA7A/PLXNC1 PCA. **a,** PC1 Loadings for S GM genes (left) and A GM genes (right). **b,** Projection of the PC1 scores of the adult data onto cortical surface renderings for S GM (left) and A GM (right). **c,** PCA using *SEMA7A* and *PLXNC1* combined expression at p3, 6, and 13. We observed that A1C at post-conception week 3 (p3) exhibits the highest variation in gene expression among prospective fetal primary area samples. We suspect that this variability may, in part, stem from challenges in accurately identifying A1C based on anatomical landmarks and precisely dissecting this region in younger and smaller fetal brains, compared to other prospective areas.

**Extended Data Figure 9:**
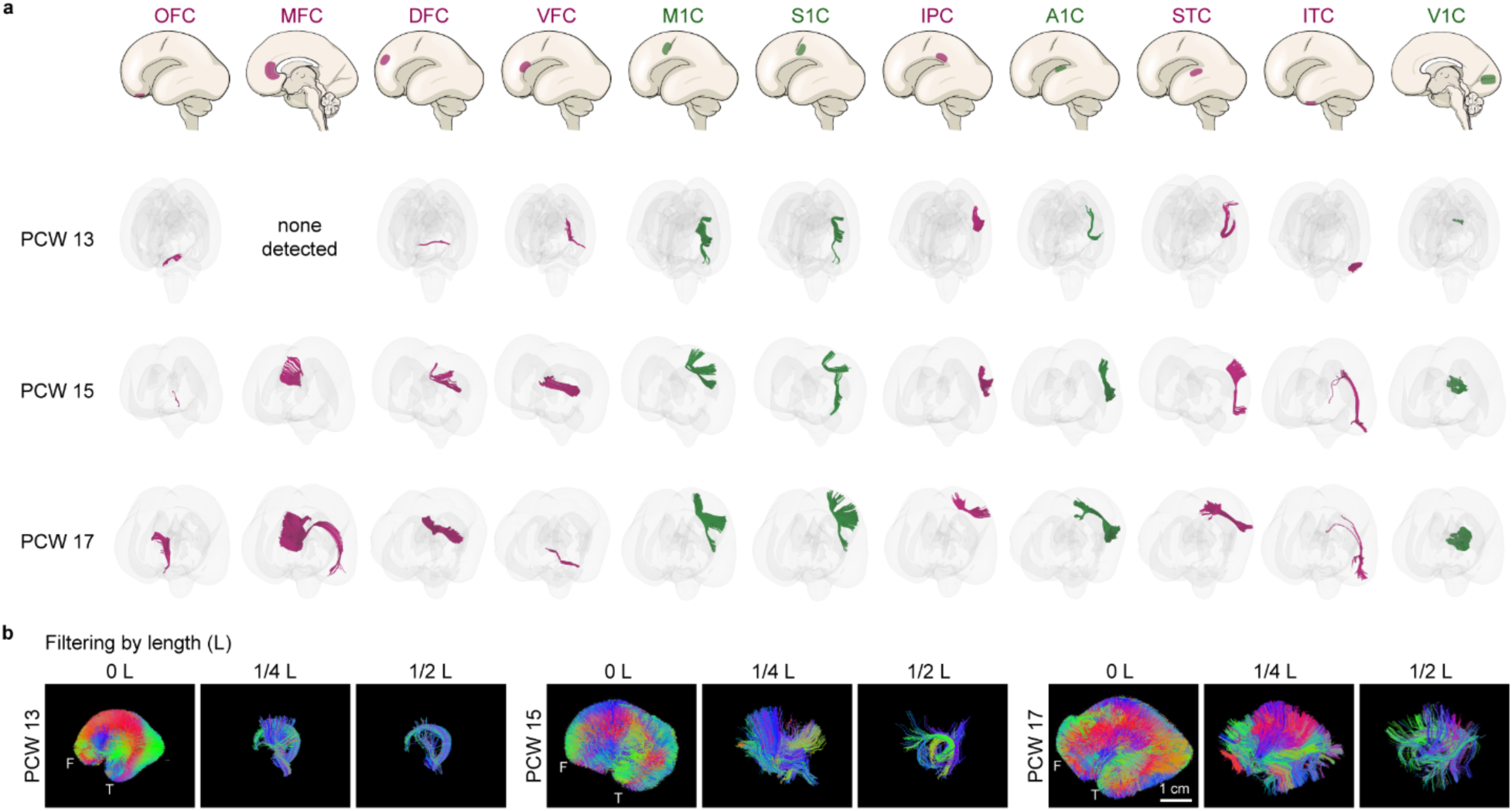
DWI in early to mid-fetal human brains. **a,** ROI-based tractography of cortico-cortical fibers across three developmental time points (PCW 13, PCW 15, and PCW 17). Fibers were traced from 11 cortical ROIs, with association areas shown in magenta and primary sensorimotor areas in green. Cortico-cortical tracts were extracted using logical filtering, retaining only streamlines with one or both endpoints located within volumetric ROI labels and confined to the cerebral wall segmentation. **b,** Lateral views of whole-brain cortico-cortical fiber tracts for fetal brains at PCW 13, PCW 15, and PCW 17. Fiber tracts were filtered by length, using the anterior-posterior extent (L) of each brain as a reference. Three thresholds were applied: unfiltered tracts (0L), tracts longer than 1/4L, and tracts longer than 1/2L. F, frontal; T, temporal.

**Extended Data Figure 10:**
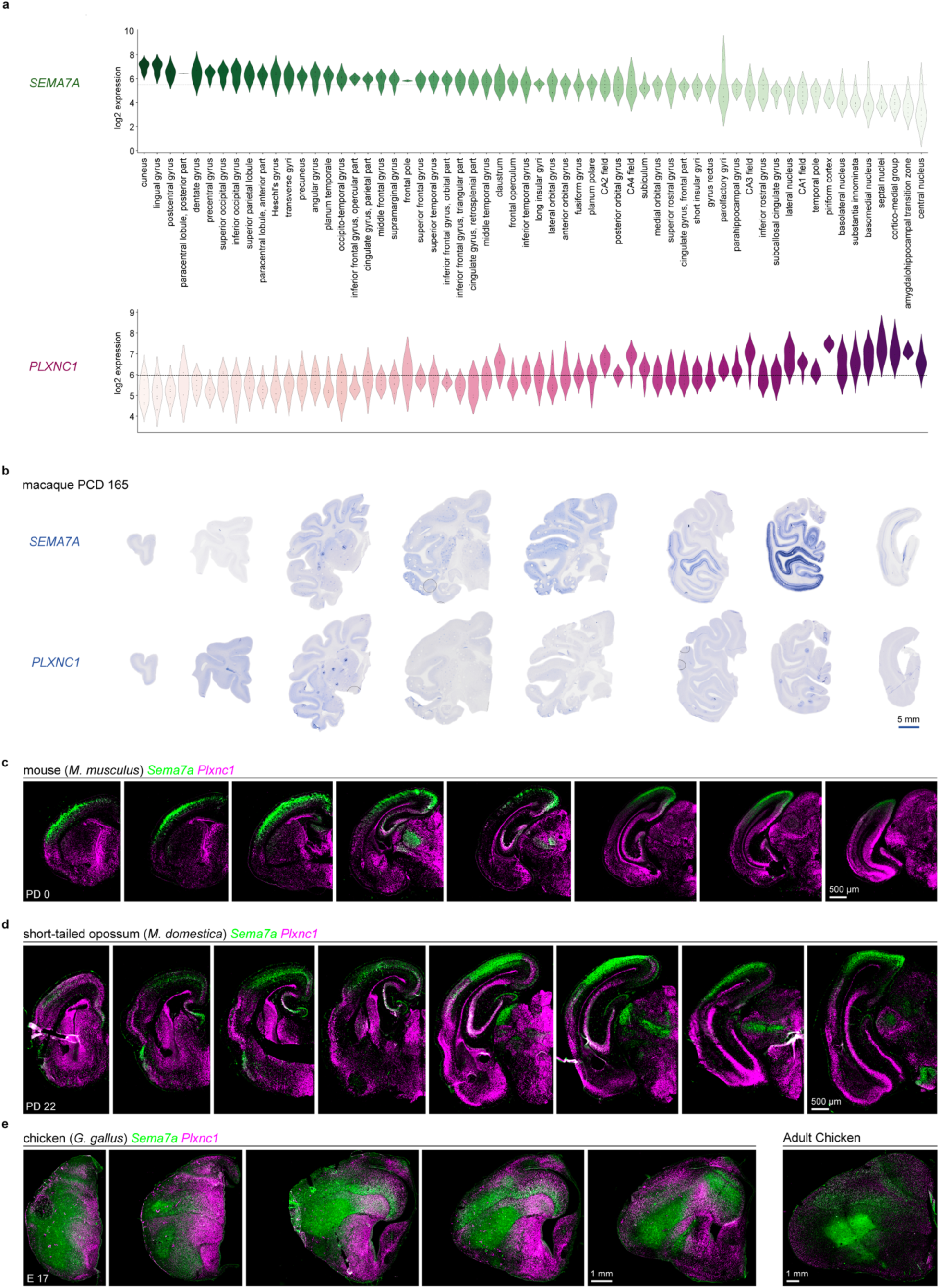
*Plxnc1* and *Sema7* expression in eutherian, marsupial, and avian brains. **a,** *SEMA7A* and *PLXNC1* expression across human neocortical and paleocortical regions based on microarray data generated from adult donors^32^. **b,** In-situ hybridization using anti-sense probes for *SEMA7A* and *PLXNC1* with post-conception-day (PCD) 165 (birth) macaque brain sections. **c,** RNA-scope using *Sema7a* and *Plxnc1* probes on PD 0 mouse (eutherian) brain sections. **d,** RNA-scope with *Sema7a* and *Plxnc1* probes on PD 22 (roughly developmentally equivalent to mouse PD 0) Short-tailed opossum (marsupial) brain sections. **e,** RNA-scope with *Sema7a* and *Plxnc1* probes on Chicken (avian) embryonic day (E) 17 and adult brain sections.

**Extended Data Figure 11:**
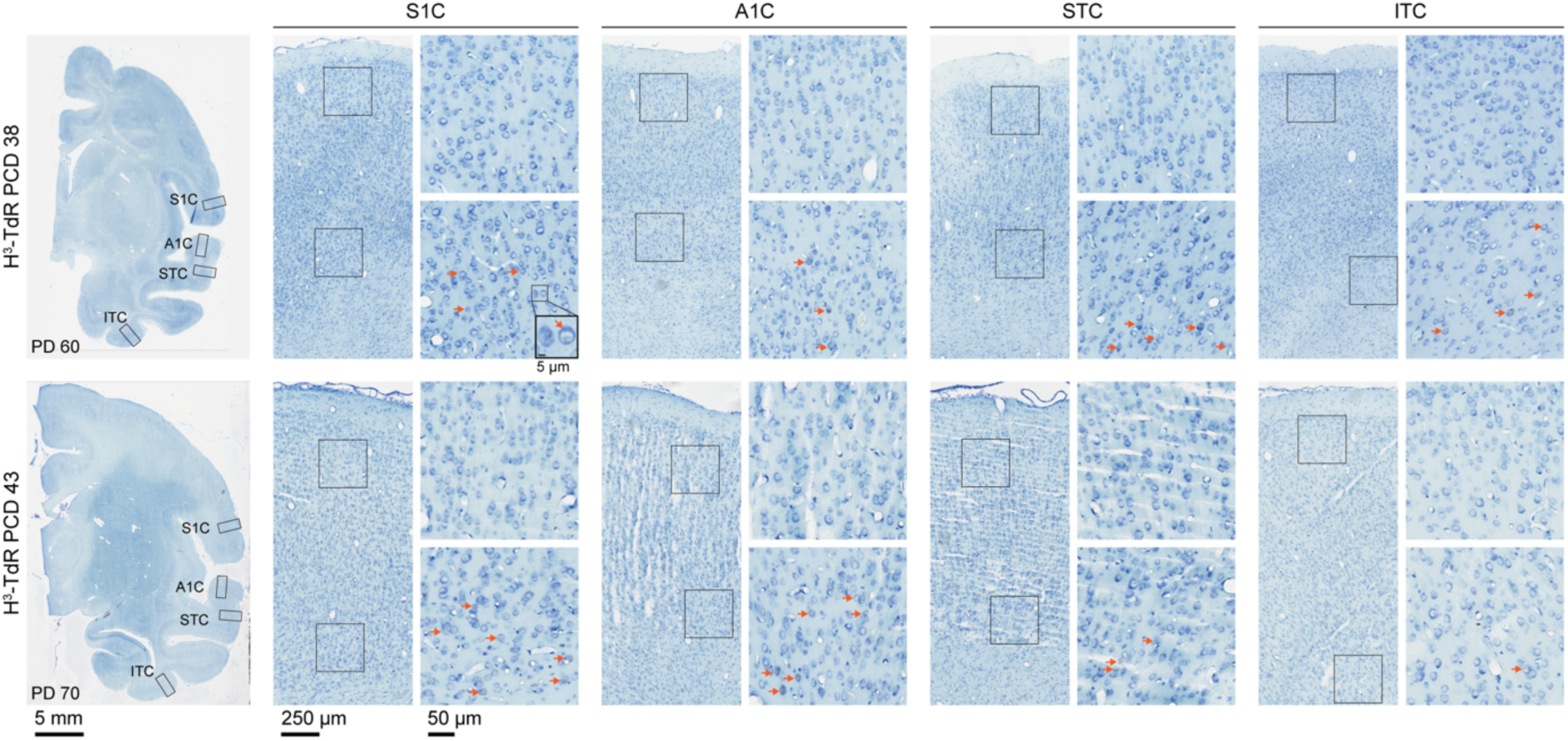
Representative sections from tritiated thymidine injected macaque brains. Brightfield images of representative archival brain sections from macaques injected with thymidine-methyl-H^3^ (also known as tritiated thymidine, H^3^-TdR) at PCD 38 or PCD 43. Low magnification representative images of 8 µm thick coronal brain sections stained with toluidine blue are shown with indicated regions examined. Orange arrows indicate example H^3^-TdR positive cells.

**Extended Data Figure 12:**
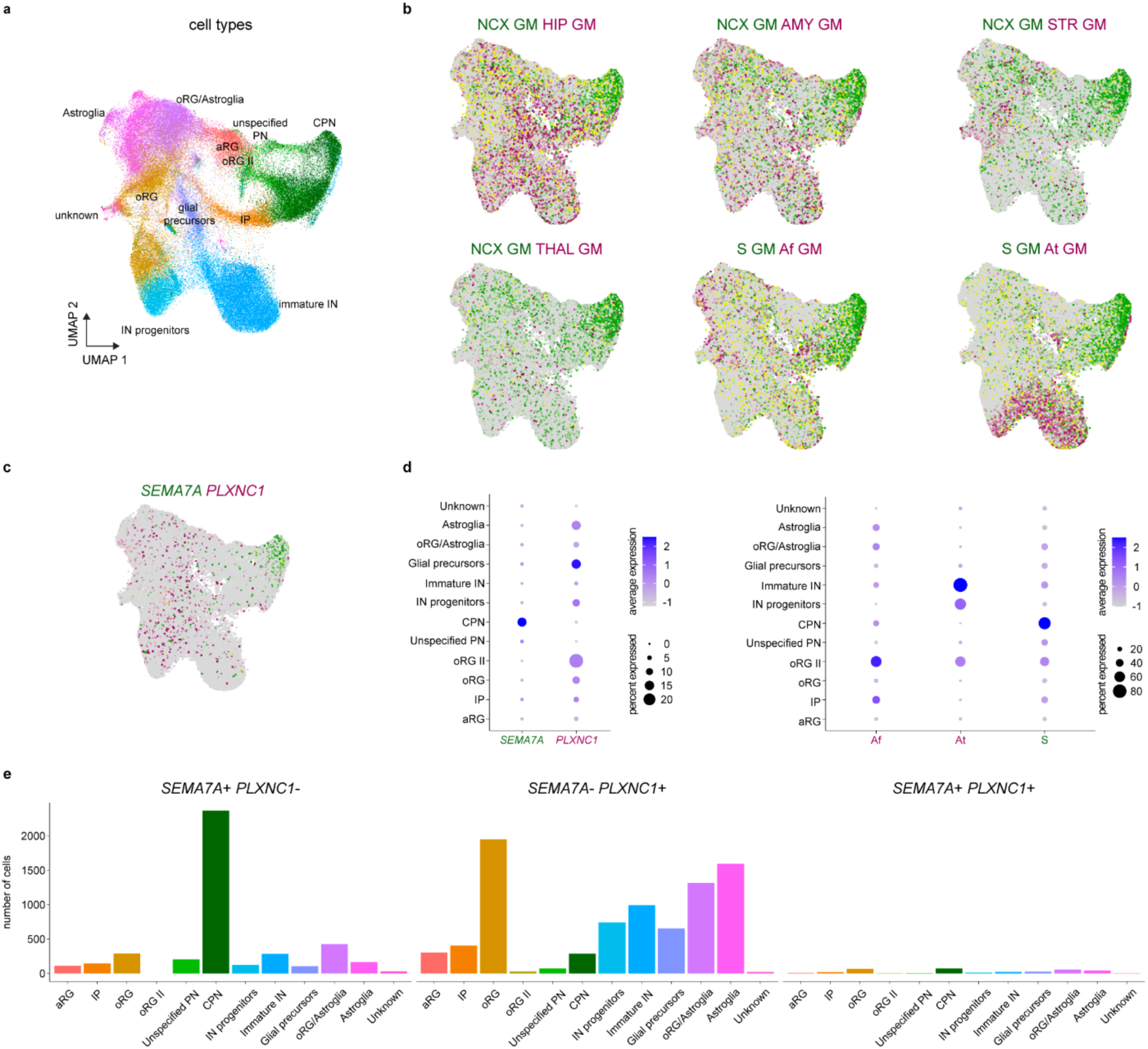
Human cerebral organoids model naïve neocortex. **a,** UMAP of 6-month-old cerebral organoids using a published single cell sequencing dataset with cell types annotated in Uzquiano et al., 2022^52^. **b,** Expression of GMs in magenta or green, yellow indicates co-expression. GM, gene module; Ncx, neocortex; Hip, hippocampus; Amy, amygdala; Str, striatum; Thal, thalamus; S, sensorimotor; Af, association frontal; At, association temporal. **c,** UMAP showing *SEMA7A* and *PLXNC1* expression. **d,** Dotplots indicating gene or gene module expression across cell types. **e,** Number of cells expressing either *SEMA7A* (*SEMA7A*+/*PLXNC1*), *PLXNC1* (*SEMA7A*-/*PLXNC1*+), or *SEMA7A* and *PLXNC1* (*SEMA7A*+/*PLXNC1*+). In cerebral organoids, the expression of *SEMA7A* and *PLXNC1* is highly segregated.

**Extended Data Figure 13:**
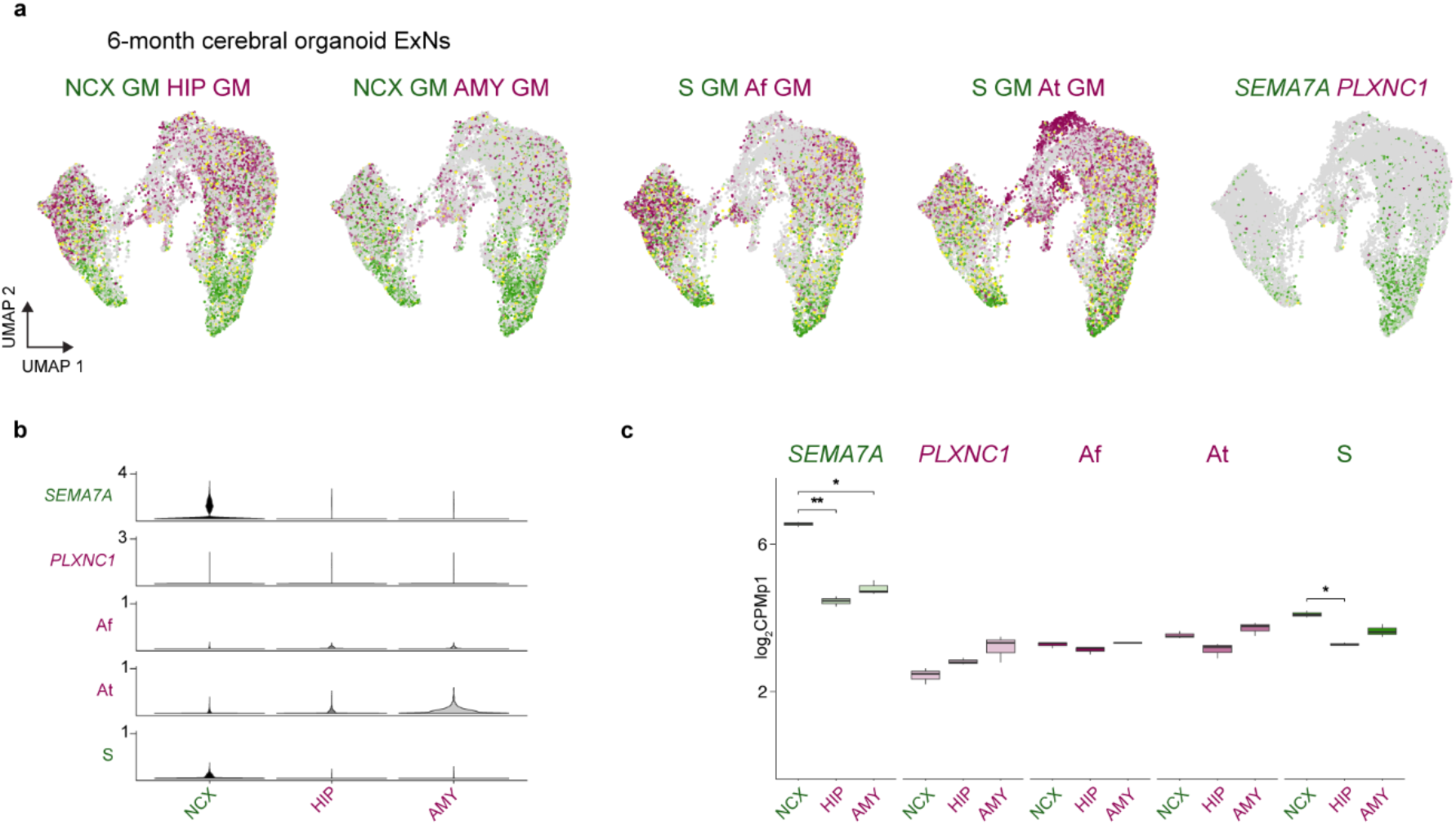
Intrinsic SEMA7A expression in 6-month human cerebral organoid ExNs. **a,** GMs or gene expression feature UMAPs of the ExNs subset from 6-month-old cerebral human organoids^52^. We identified cells with neocortex (NCX), hippocampus (HIP), and amygdala (AMY) molecular identities. Expression of GMs or genes in magenta or green, yellow indicates co-expression. GM, gene module; S, sensorimotor; Af, association frontal; At, association temporal. **b,** Violin plots showing distribution of gene expression or module score across the cells categorized as NCX, HIP, or AMY. **c,** Boxplot illustrating pseudo-bulk log_2_(CPM + 1) expression levels of genes or gene modules. The cells expressing NCX module genes showed signatures of *SEMA7A* and S GM expression enrichment but not *PLXNC1* and A GMs similar to our observations in the early fetal neocortex. This suggests that projection neurons in these organoids, which lack ventral patterning centers, are of naïve cortical identity. We used a *t*-test with FDR < 0.05 to determine significance. See **Supplementary Table 18** for detailed statistical information.

**Extended Data Figure 14:**
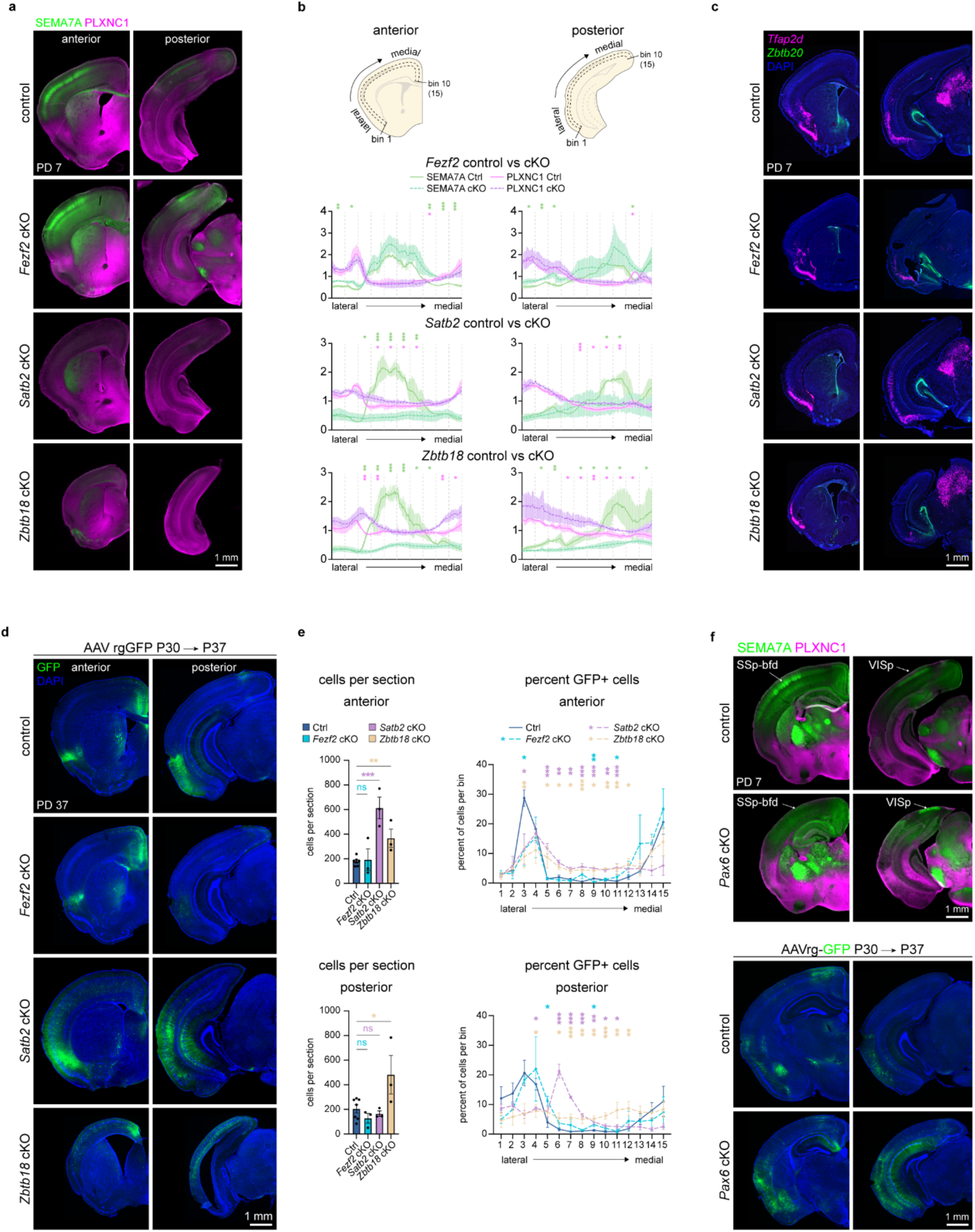
Additional A-P positions analyzed in *Fezf2*, *Satb2*, *Zbtb18* cKO cortices and *Pax6* cKO analysis. **a,** PD 7 control (*Satb2^flox/flox^*), *Fezf2* cKO, *Satb2* cKO, and *Zbtb18* cKO brain sections stained for SEMA7A and PLXNC1. **b,** Top: cartoon depicting regions quantified. Quantifications of signal intensity relative to the mean signal intensity in control versus *Fezf2*, *Satb2*, *Zbtb18* cKOs. The lateral to medial region quantified was divided into 10 bins, and significance was determined based on the average signal intensity in each bin. **c,** RNA-scope experiments probing for *Tfap2d* and *Zbtb20* in PD 7 control (*Satb2^flox/+^*), *Fezf2* cKO, *Satb2* cKO, and *Zbtb18* cKO brain sections. **d,** GFP signal in sections from PD 37 control, *Fezf2* cKO, *Satb2* cKO, and *Zbtb18* cKO mice in which rgGFP-AAV was injected into the mPFC at PD 30. **e,** quantification of average cells per section in indicated genotype and the percent distribution of cells across 15 equal sized bins from the lateral to medial region of the cortex depicted in panel b. **f,** Top: Staining for SEMA7A and PLXNC1 in control (*Pax6^flox/flox^*) and *Pax6* cKO cortices with selected primary areas labeled at PD 7; bottom: PD 37 sections from mice subject to mPFC AAVrg-GFP injection at PD 30. For all quantifications a *t*-test was performed, error bars show standard error of the mean. \**p* < 0.05, \*\**p* < 0.01, \*\*\**p* < 0.001. See **Supplementary Table 19** for detailed statistical information.

**Extended Data Figure 15:**
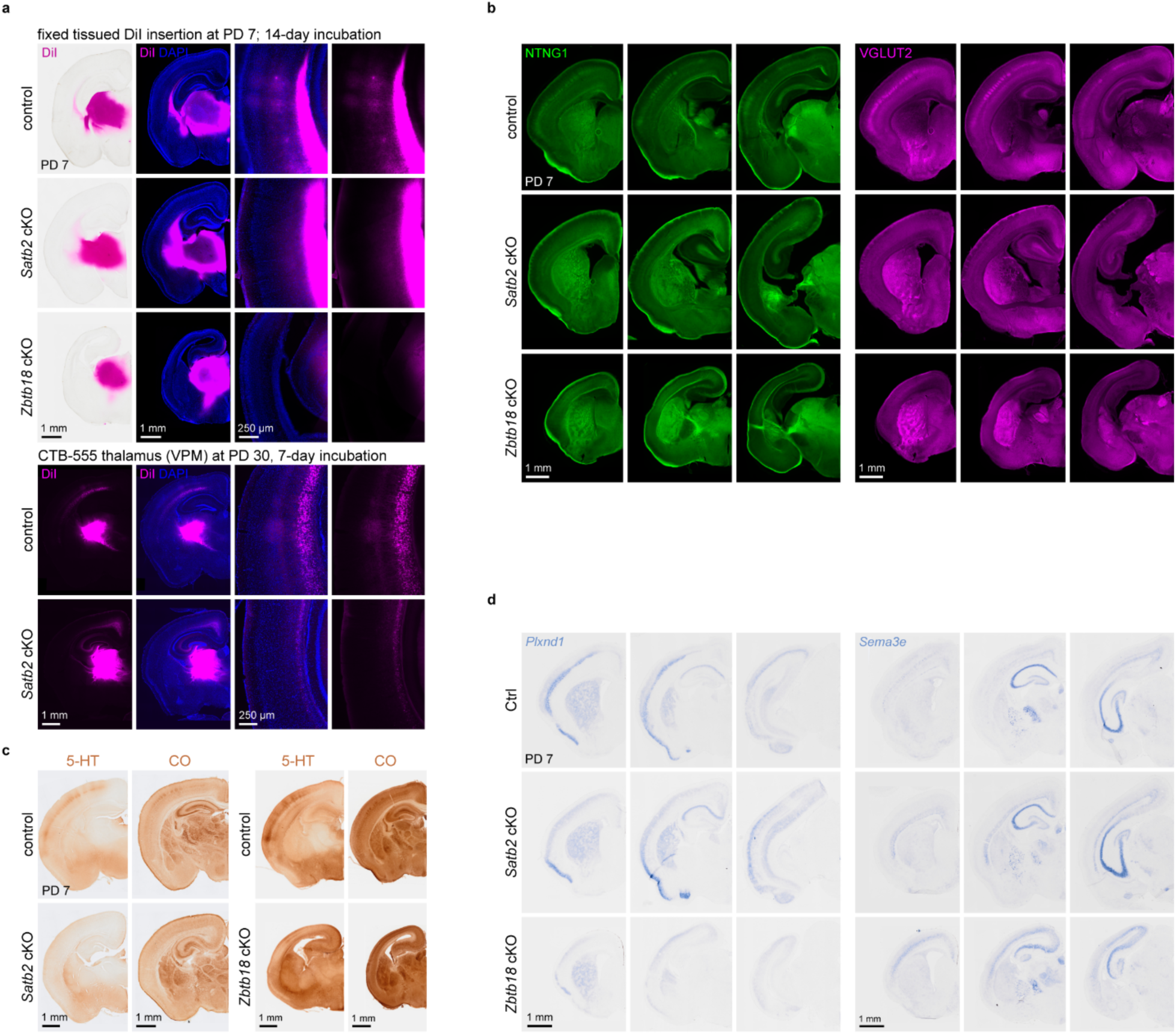
Loss of primary sensory areas and TCAs in *Satb2* and *Zbtb18* cKO cortices. **a,** Top: Images of PD 7 mouse brains in which DiI was placed into the thalamus followed by one week incubation. Brightfield and fluorescence low magnification images are shown next to high magnification images of the cortex. DiI signal can been seen in L4 and 6 in controls only. In *Satb2* cKOs, L6 DiI signal was observed. No cortical signal was observed in *Zbtb18* cKOs. Bottom: additional tracing was performed in *Satb2* cKOs at PD 30 by injecting CtB into the thalamus and collecting the brains at PD 37. CtB signal was seen in retrogradely labeled cell bodies in L6 in control and less so in Satb2 cKOs. Anterograde L4 signal was observed in controls, but not *Satb2* cKOs. **b,** Staining for thalmo-cortical afferent (TCA) markers NTNG1, and VGLUT2 in control, *Satb2*, and *Zbtb18* cKOs. **c,** 5-HT staining and cytochrome oxidase (CO) activity in control, *Satb2*, and *Zbtb18* cKOs at PD 7. **d,** In situ hybridization performed with anti-sense probes to *Plxnd1* and *Sema3e*, two genes known to be required for TCA innervation of the cortex, at PD 7 in control, *Satb2* cKO, and *Zbtb18* cKOs.

**Extended Data Figure 16:**
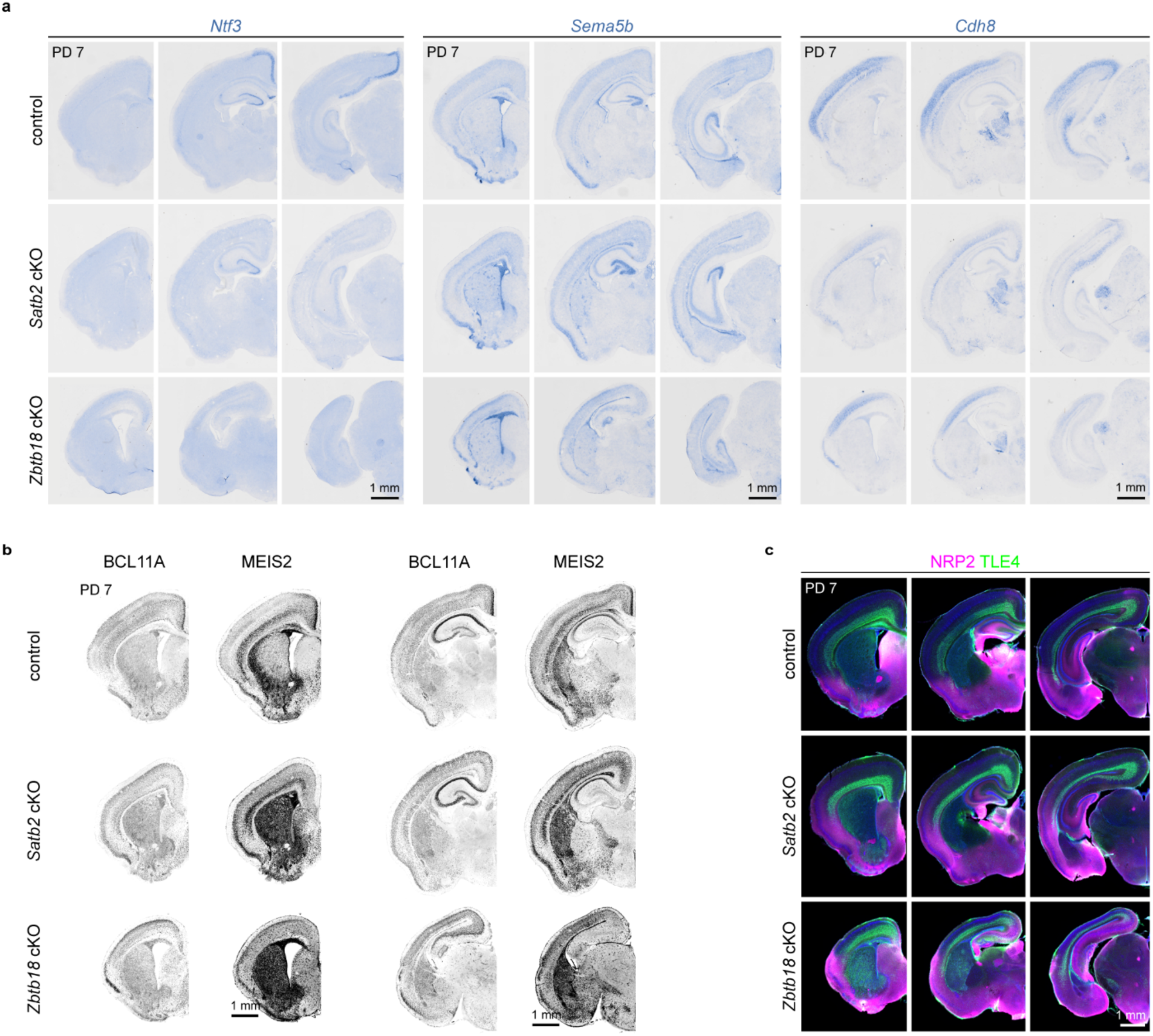
Additional markers in *Satb2* and *Zbtb18* cKOs. **a,** In situ hybridization with antisense probes for *Ntf3*, *Sema5b*, and *Cdh8* using PD 7 control, *Satb2* cKO, and *Zbtb18* cKO brain sections. **b,** Staining for the primary sensory area enriched BCL11A and association and deep-layer marker MEIS2 in PD 7 control, *Satb2* cKO, and *Zbtb18* cKO brain sections. **c,** Staining for NRP2 and TLE4 in PD 7 control, *Satb2* cKO, and *Zbtb18* cKO brain sections.

**Extended Data Figure 17:**
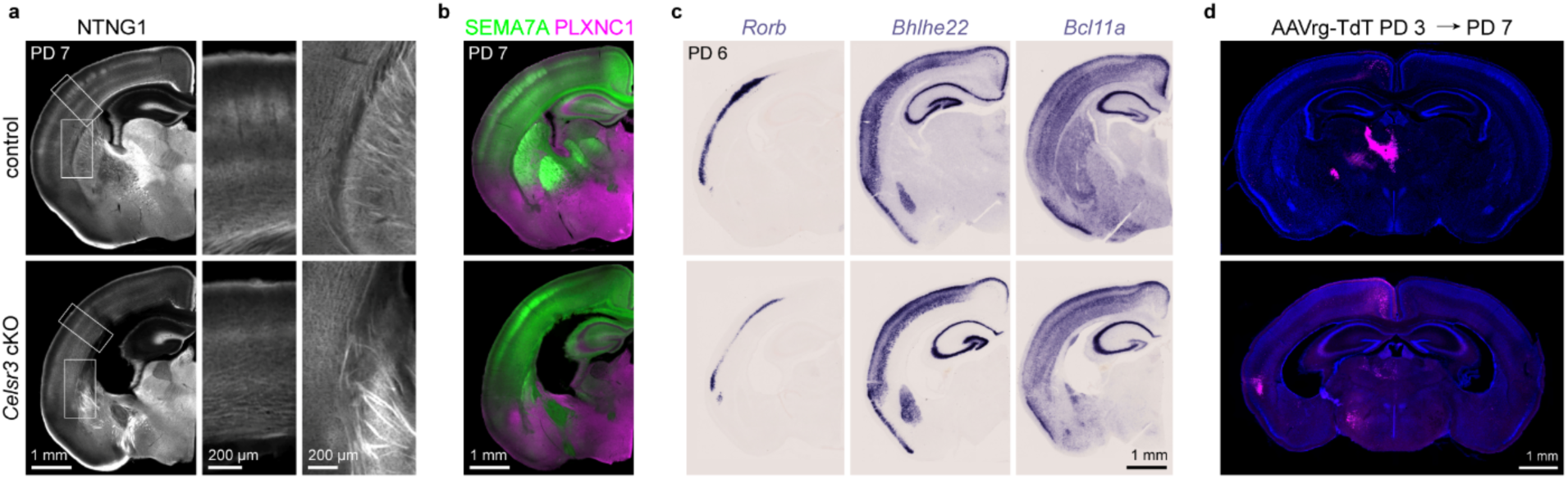
Reduced TCAs and poorly defined areal boarders in *Celsr3* cKO. **a,** Staining for NTNG1 in PD 7 control and *Celsr3^flox/flox^*; *Dlx5/6-Cre* (*Celsr3* cKO) mouse brain sections. **b,** SEMA7A and PLXNC1 staining in control and *Celsr3* cKOs at PD 7. **c,** In situ hybridization using antisense probes for *Rorb*, *Bhlhle22*, *Bcl11a* at PD 6. **d,** Brain sections from PD 7 mice in which AAVrg-TdT was injected into mPFC at PD 3.

**Extended Data Figure 18:**
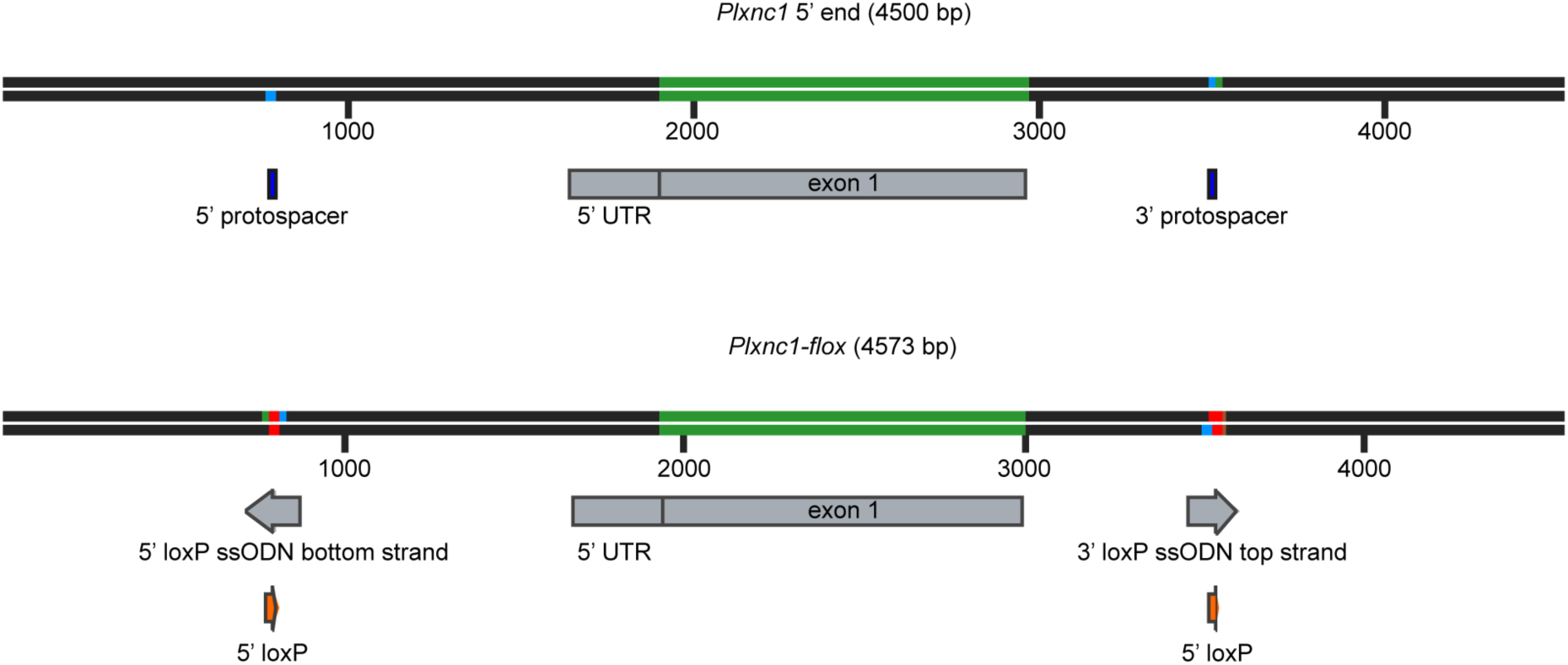
Design of *Plxnc1-flox* allele. Top: structure of the 5’ end of *Plxnc1* showing exon 1, the 5’ untranslated region (UTR) and adjacent intronic regions. Locations of CRISPR/Cas9 target sites (5’ and 3’ protospacers, blue) used for insertion of indicated loxP sites below. Bottom: structure of the edited *Plxnc1-flox* allele, in which loxP sites (orange) have been inserted flanking exon 1 using single-stranded oligodeoxynucleotide (ssODN) templates. This design enables Cre recombinase-dependent excision of exon 1.

## Notes

### Competing Interest Statement

The authors have declared no competing interest.

