## Extended Discussion for "Competing Programs Shape Cortical Sensorimotor-Association Axis Development"

We present experimental evidence and a conceptual framework describing how the development of primary sensorimotor and association areas, and their corresponding networks, is orchestrated by competing identity programs that emerge centrally and pericentrally along the developing neocortical sensorimotor-to-association (S–A) axis, or more broadly, the sensorimotor–association–limbic (S–A–L) axis. These programs arise from opposite poles of the axis and expand toward one another during cortical development.

In mice, pericentral programs emerge neocortically at distinct anterior ventromedial frontal and posterior ventrolateral temporal nodes, situated near the septal and amygdalar poles, respectively, and adjacent to allocortical and ventral pallial structures. Given that our primate analyses were limited to transcriptomic data from 11 neocortical areas, along with histological assessments of a small set of genes within shared frontal and temporal association gene modules (Af and At, respectively), we hypothesize that these programs progressively expand outward along the peri-allocortical border and inward along dominant frontotemporal (F–T) developmental trajectories.

Notably, prior studies from multiple groups have shown that first-order (FO) sensorimotor thalamocortical afferents (TCAs) enter the subplate and neocortical plate within central territories[1], shortly after the pericentral programs begin to emerge and move inwards. These afferents establish distinct, topographically segregated patterning nodes (i.e., sites of interaction between ingrowing FO TCAs and the naïve immature neocortex), where they exert early inductive effects on the molecular, cellular, and connectional properties of the emerging primary sensorimotor areas[2–8]. In doing so, they preclude later-arriving pericentral programs from colonizing regions that have already been committed to forming primary sensorimotor cortex.

The early topographic sorting of growing FO TCAs within the ventral pallium[4,9–14], followed by the topographic segregation of primary somatic sensorimotor, auditory, and visual cortical patterning nodes, each associated with distinct spontaneous or sensory-evoked activity patterns originating from their respective peripheral sensory organs[2–4,7], works in coordination with the induced central molecular programs that are either shared (e.g., SEMA7A) or specific to individual nodes. Together, these mechanisms ensure that each primary network becomes uniquely specified and establishes a modular cortical architecture.

In contrast, association networks become progressively more distributed and less structurally and topographically constrained. They receive weak or no direct input from FO TCAs and instead connect to higher-order thalamic nuclei. These networks also form increasingly long-range, often cross-hemispheric, cortico-cortical connections that engage a spectrum of higher-order sensorimotor, transmodal, and limbic regions[3,15–24]. These features enable integration across multiple sensory modalities, internal states, and memory systems. We present multiple lines of evidence that the patterning of association networks along the neocortical S–A axis cannot be explained by primary patterning anchors alone, but is instead shaped by the interaction of both pericentral and primary area–anchored central identity programs.

Furthermore, we hypothesize that, due to the gradual inward spread of pericentral programs, the early-specified primary nodes exert activity-dependent influences (patterning anchors) on the surrounding prospective unimodal association cortices before these regions are further refined by the delayed arrival of the pericentral programs. Consistent with this spatiotemporal sequence, studies across multiple species have shown that early activity in developing cortical networks follows region-specific patterns shaped by peripheral sensory inputs[2–4,25]. For example, retinal waves serve as the primary driver of neuronal activity in the nascent primary visual cortex but merely modulate ongoing activity in secondary visual areas[26]. As the neocortex matures, the influence of these sensory-petal inputs diminishes, indicating an increasing dominance of intrinsic cortical activity, which begins to compete more directly with signals from the sensory periphery[27–30]. Notably, recent findings demonstrate that modular cortical activity patterns can emerge through self-organizing local excitatory and inhibitory dynamics within developing cortex and can be modified by extrinsic stimulation[31]. Compared to primary sensory areas intrinsic pattering activity may be more influential in the prospective unimodal association areas. This and the additional influence of incoming pericentral programs, may underlie the gradual emergence of graded areal boarders and connectivity features observed in the adjacent unimodal association areas.

During human development, certain aspects of cortical areal and network maturation follow a hierarchical sequence. Myelination begins before birth and proceeds along multiple gradients[32–34]. Notably, there is an early outside-in gradient, where myelination starts in select frontal and temporal (para)limbic regions and extends into adjacent developing transmodal cortices. An inside-out gradient emerges from primary sensorimotor areas, progressing outward toward nearby unimodal association cortices. In contrast, transmodal areas and their long-range connections are not fully myelinated until after birth, continuing well into adolescence and early adulthood, much later than other cortical regions[32,34]. This developmental sequence reflects a hierarchical gradient of functional complexity, in which basic sensory and motor processing capabilities emerge before the higher-order integrative and cognitive functions supported by transmodal networks[32,33].

Correspondingly, electroencephalographic studies have shown that stimulus-evoked potentials across the three major sensory modalities originate from both primary and secondary regions during human late-fetal development[35]. In contrast, association cortex potentials do not appear until early childhood. This early emergence of cortical electrogenesis associated with prospective sensory cortices suggests an early extrinsic impact on the development of those cortical regions[35]. A comparable spatiotemporal pattern has also been observed in human structural and functional MRI studies, where unimodal sensorimotor networks reached their mature positions early in development, while transmodal association networks continued to mature throughout development into early adulthood[17,36–47].

Together, these findings and time windows align with our observation of a slightly delayed yet rapid upregulation and compartmentalization of central 'sensorimotor' GMs, along with an early emergence but slow inward progression of pericentral GMs into the surrounding prospective unimodal association areas in mice, macaques, and humans. We hypothesize that this delay in the inward expansion of pericentral programs allows a specific hierarchical sequence to unfold: central programs, driven by activity-dependent mechanisms, specify the sensoritopic and topological proto-architecture of primary areas, which serve as patterning anchors to then shape key features of higher-order, adjacent, unimodal association cortices and networks [2–5,7,25,27–30,33,48–58]before the arrival of invading pericentral programs which shape key features of higher-order and transmodal association areas and networks.

Our model and experimental data indicate that the full development of higher-order unimodal sensorimotor association features, as well as transmodal association networks, requires additional and distinct developmental mechanisms. We show that these mechanisms are at least partially related to pericentral programs, which originate independently of, and act likely separately from, the mechanisms associated with primary sensorimotor patterning nodes. These pericentral programs exhibit unique transcriptional and spatiotemporal profiles. Rather than replacing the central programs, they compete with them for territory within the early, naïve neocortex. Neither program appears to represent a default developmental state; instead, both emerge from active and passive interactions between nascent neocortical neurons and distinct non-neocortical structures. Our re-analysis of cortical-like organoids, which lack the nodal inductive regions hypothesized by our model, likely represent the true default developmental state because neither the primary nor association specialization emerges.

Our findings support the view that the emerging neocortex begins in a naïve, areally unspecified or default-like state, consistent with the classical concept of a "protocortex"[59]. A key aspect of the MIND model is that both sensorimotor and association features emerge gradually, through distinct molecular and spatiotemporal mechanisms. These emerging pericentral association and central sensorimotor identity programs engage in a competitive interplay, essentially a “space race,” for neocortical territory which eventually shape the S–A axis. This progression from an early fetal naïve/unspecified, to late fetal organization along an S–A axis is reflected in gene expression gradients. In the early fetal brain at period 3 (p3), however, gene expression appears polarized along a dominant F–T axis. It is possible that there are early genetic distinctions between sensorimotor and association areas that we have not observed due to some limitations of our analytical framework. For example, the limited number of cortical areas (11) included, and the absence of ventrally located (e.g., insular) and dorsally located (e.g., central cingulate) regions in our analysis. However, the earliest signs of pericentral programs we observe appear in the most pericentrally/peripherally located fronto-temporal neocortical cortices (medial and orbital prefrontal cortex, and inferior temporal cortex), and begin to move inward, and the focal, FO TCA-induced, central sensorimotor programs emerge slightly later. During the late-fetal periods, we observed a transcriptomic shift in which gene expression variability becomes increasingly polarized along an emerging S–A axis. The simplest interpretation for these observations is that the earlier molecular organization of the naïve neocortex follows an F–T topography which becomes overwritten by the later emergence and propagation of the mutually exclusive pericentral association and central sensorimotor programs, resulting in mature functionally relevant S–A axis organization.

This dynamic molecular landscape suggests that the initial F–T polarization of the naïve neocortex sets the stage for the emergence of these distinct pericentral and central programs that encompass distinct sets of genes encoding both instructive and permissive factors known to influence neural cell and circuit development. Through inductive and exclusionary processes, these competing programs reach an equilibrium that establishes the spatial topography of the S–A axis such that primary sensorimotor areas emerge as focal "islands" embedded within a broader "ocean" of association cortex, which is hierarchically structured from adjacent unimodal association areas to more distributed transmodal networks primarily patterned along an F–T gradient. Collectively, our findings and model propose that the development of the S–A axis is shaped by the interplay of these major opposing programs, alongside activity-dependent, temporal, geometrical, and topological influences and constraints.

**Induction and exclusion principles**

We demonstrate that molecular features of central and pericentral programs, along with their precisely choreographed spatiotemporal dynamics and competitive interplay, generate distinct territories along the S–A axis through dual mechanisms of induction and exclusion. This process orchestrates spatial gradients and compartmentalization of key axon guidance, cell-cell adhesion, synaptogenesis, retinoic acid signaling, Wnt signaling, and autism risk genes. Together, these factors provide graded and spatially coordinated cues that pattern the emerging S–A axis, including the early F–T polarization of higher-order and transmodal association projections.

Specifically, we demonstrate that the complementary expression of PLXNC1 and SEMA7A, a mutually repulsive axon guidance pair known for bidirectional signaling and involvement in the formation of neural circuit topologies, and activity-dependent synaptogenesis and thalamocortical axon branching[60–75], contributes to the antagonistic shaping of inter-areal cortico-cortical axons between primary sensorimotor and association regions.

The principles of induction and exclusion underlying this compartmentalization are consistent with certain previous findings of histochemical, molecular and electrophysiological differences between prospective primary and association regions of the subplate zone or neocortical plate, differences that arise prior to the major onset of intra-neocortical synaptogenesis, or the emergence of obvious areal cytoarchitectonic or connectional differences between those regions[76–92]. One illustrative example of these principles is the anticorrelated expression and complementary repulsive function of SEMA7A and PLXNC1. SEMA7A is expressed most strongly in the primary area of the dominant sensory modality of a given species (i.e., primary somatosensory cortex of mice, and primary visual cortex of primates), specifically in the thalamo-recipient layer 4, which also exhibits the sharpest areal boarders due to outsized sensory-petal influence. These primary areas act as patterning anchors, providing the inductive cues for lower-level SEMA7A expression in the adjacent unimodal association areas. These gradients in SEMA7A expression contributes to differential axonal and dendritic repulsion, allowing for transmodal axons to more readily extend into unimodal association areas where SEMA7A levels are lower.

In addition to layer-specific differences in SEMA7A and PLXNC1 expression between emerging primary and association areas, and their role in shaping distinctive cortico-cortical connection patterns, additional evidence supports the principal of exclusion in the prospective primary visual cortex (V1C). Notably, V1C, and other primary regions typically either lack or exhibit markedly reduced callosal projections[93–96]. Interestingly, the expression of ROBO1, a receptor involved in the formation of callosal projections[97,98] and associated with reading and mathematical abilities[99–101], is downregulated in the prospective fetal human V1C compared to adjacent association areas[83]. Consistent with these findings, *ROBO1* is enriched in the pericentral association programs and in prospective association areas surrounding V1C and other non-primary visual regions of the fetal macaque and human cortex[85,89,102,103]. Similarly, PLXNC1, which is also depleted in both human and non-human primate V1C, is enriched in callosally projecting excitatory neurons (ExNs) of transmodal association areas in the macaque cortex[72], suggesting a role in promoting interhemispheric connectivity selectively outside primary areas. Supporting this notion, tracing performed in infant macaques show that higher-order and transmodal areas of the macaque inferior temporal cortex do not receive transient long-range cortico-cortical projections from V1C. Furthermore, within the ipsilateral hemisphere, the spatial and laminar distribution of labeled cortico-cortical neurons closely resemble those seen following comparable injections in adult macaques[104]. These findings are consistent with our model and support the notion that activity-dependent and molecularly defined exclusion between primary and higher-order sensorimotor areas mediates, in a layer-specific manner, the segregation of primary and association networks throughout development and across processing hierarchies.

Another example of the induction-exclusion principle is the TCA-dependent expression of CYP26B1 in the prospective motor and insular cortex[105], which restricts RA signaling and associated genes, such as *Cbln2*, preferentially to the PFC[83,105–108], supported by our finding of the dorsolateral frontal expansion of PLXNC1 expression and PFC-like association intra-cortical connectional features in short-tail opossum (*Monodelphis domestica*), which lacks a true primary motor cortex[109–113]. Taken together, these findings suggest that induction and exclusion function as fundamental, yet opposing, mechanisms by which pericentral and central programs shape the S–A axis and contribute to the establishment of cortical processing hierarchies.

**The emerging concept of pericentral "association" identity programs**

Our findings demonstrate that the proper development of key molecular and long-range cortico-cortical connectional features of the association cortex requires additional mechanisms and principles distinct from, but interacting with, those involved in primary sensorimotor areas. These mechanisms appear not to involve early sharp compartmentalized area-specific gene expression, unlike in the developing primary motor and sensory areas. Rather, spatiotemporal gradients of transcriptional programs or waves initiated at specific nodal sites at the neocortical edge propagate towards the center but are excluded from territories occupied by the arrival of sensory TCAs and the induction of primary sensorimotor-specific gene expression programs. Many of the key genes within the pericentral GMs, such as *Plxnc1*, *Pcdh10*, and *Pcdh17*[82], are enriched across the immature layers of the prospective paralimbic and transmodal association cortices. To a lesser extent, they are also expressed in the immature deep, but not upper, layers of the prospective sensorimotor cortices in neonatal mice. Interestingly, both *Plxnc1* and *Pcdh10* are also expressed in the neocortical proliferative region[82], though not in any obvious gradient at this developmental period. Furthermore, both plexins and protocadherins have been shown to functionally interact with other proteins found in the shared or individual pericentral Af and At gene lists, such as the MET receptor tyrosine kinase, neuropilin axon guidance proteins, and components of the Wnt signaling pathway, all of which have been previously implicated in cortical development and neurodevelopmental disorders[60,62,114–116]. These findings suggest that combinatorial interactions among these proteins may underlie the diverse effects of pericentral gene programs.

We also observed that pericentral programs share certain genes like *PLXNC1*, and contain program-specific genes, reflecting their different nodal initiation sites (i.e., frontal or temporal). At least in case of *PLXNC1* and other select genes, we showed that the F–T polarization and inward propagation generate layer-specific gradients across prospective association areas. We hypothesize that such a spatiotemporal pattern of articulation supports the formation of distributed, graded architecture rather than sharply defined association areas and sub-networks. Through this process, specialized cognitive modular networks underlying distinct functions may gradually emerge and become increasingly refined over the course of an individual's development. We speculate that this developmental framework facilitates the formation of a highly distributed architecture and graded differences in the laminar origin and termination between inter-areal connections that support the confluence of multiple cognitive processes to perceive a specific thing.

Furthermore, this model provides mechanistic explanations for how the cortical cyto- and myelo-architectonic gradients and long-range intra-cortical connectional trends previously identified extending from neocortical borders with (para)limbic periallocortex toward primary areas located more centrally are generated,[117–120] and how some regions such as posterior parietal cortex become transmodal while intercalated between unimodal sensorimotor areas. We hypothesize that pericentral programs interact with self-organizing sensoritopic and activity-dependent mechanisms across fields and layers of prospective unimodal association cortices, which progressively start to acquire certain intra-cortical connectional features shared with higher-order association, but not primary sensory areas, such as direct connections with the transmodal cortices and prominent callosal connections. We hypothesize that these features are formed by distinct layer and cell-type specific neuronal types, as even *Plxnc1* and other pericentral program genes display both area- and layer-specific expression along the S–A axis.

Experimental studies involving defects in peripheral sensory organs and rerouting of their associated TCAs, or transplantation of immature neurons from the predominantly central donor region of the neocortex have demonstrated that sensorimotor traits of primary areas are largely determined by the FO TCAs innervating the host site[121–127]. These findings support the inductive role of FO TCAs and their associated central programs. In contrast, immature neurons from prospective perirhinal transmodal or paralimbic cortices retain association-like characteristics when transplanted into central regions, where primary sensorimotor areas typically form, suggesting an early commitment to an association or limbic fate[128], reflecting the earlier emergence of peripheral programs. Our experimental evidence and model predict that differences in molecular composition and spatiotemporal progression between pericentral and central programs drive these distinct areal commitments, depending on whether immature neurons are located more centrally or in pericentral regions neighboring allocortical/limbic cortex. Furthermore, certain shared molecular and spatiotemporal features of allocortical/limbic and neocortical transmodal ExNs, driven by pericentral programs, may explain why allocortical neurons involved in olfactory sensory processing exhibit projection profiles that more closely resemble those of neocortical association areas than primary sensory areas, and why they preferentially target transmodal association regions while avoiding primary sensorimotor areas[15,20,21,129–131]. Notably, the claustrum, considered a peri-paleocortical structure[129,132], is enriched for *Plxnc1* and select other pericentral genes during development. Although the claustrum maintains topographically organized reciprocal projections to the entire cortex, the majority of its connections are with limbic and fronto-temporal associative cortices and motor cortex, with relatively less prominent inputs from primary sensory cortices[133,134]. In addition, the claustrum receives input from several subcortical limbic structures, particularly the mediodorsal nucleus of the thalamus, the basolateral amygdala, and the hippocampus[133,134], structures we also found to highly express *Plxnc1* and certain other pericentral genes.

The opposing developmental programs, along with inductive and exclusion principles, appear to be evolutionarily conserved and may extend beyond the cortex to structures such as the thalamus. In the developing thalamus, SEMA7A and PLXNC1 exhibit anti-correlated expression in first-order (FO) “sensorimotor” (SEMA7A$^+$) versus higher-order (HO) “association” (PLXNC1$^+$) nuclei, mirroring their expression patterns in the cortex. Our gene inactivation and co-culture experiments disrupted sensorimotor and association networks, confirming their cell-autonomous roles in cortical development. Moreover, the reciprocal expression of these genes across cortical regions, species, developmental stages, and cell types points to an evolutionarily conserved mechanism that predates the emergence of complex cortical networks. In particular, the conserved mutual exclusivity of *Plxnc1* and *Sema7a* expression in the telencephalon of both birds and mammals, together with a corresponding distinction between primary sensory and association connectivity, including PFC-like intracortical association features present in both marsupials and eutherians, supports the view that transmodal networks and the mechanisms guiding their development are phylogenetically ancient rather than recent evolutionary innovations. Accordingly, our findings and model share some similarities and important differences with an alternate model which proposes that the rapid evolutionary expansion of transmodal association cortices in primates effectively "untethers" those regions from strong constraints of molecular gradients and early activity-dependent sensory cascades[19]. Our model posits that the expanded geometry of the primate cortex allows for a greater expanse of cortex to be invaded by evolutionarily conserved pericentral identity programs which result in a shifted S–A equilibrium towards greater absolute and relative association cortex in expanding brains. Rather than novel evolutionary innovations, our evidence strongly suggests that the core developmental framework is conserved across mammals, and even, some non-mammals, while key molecular effectors may have undergone selective evolutionary modifications. For instance, *Plxnc1* has undergone a partial duplication unique to parrots, which has been proposed to contribute to their advanced vocal and imitative capacities[135]. In the human lineage, HAR26 and 2xHAR.178, among the fastest-evolving genomic regions since the divergence from chimpanzees, regulate *PLXNC1* expression in iPSC-derived cortical ExNs[136].

Taken together, these findings and our framework provide a unified model of how evolutionarily conserved pericentral programs shape key features of association networks progressing spatiotemporally from prospective (para)limbic, to transmodal, to unimodal cortices. They also open the door to new questions and avenues of inquiry.

**Mechanisms regulating and mediating pericentral programs**

Inductive interactions mediated by diverse mechanisms and signaling pathways are essential for cell differentiation and regional patterning in the telencephalon[137–140]. Through Gene Set Enrichment Analysis of pericentral shared and individual dataset GMs, we identified significant enrichment for genes encoding proteins involved in canonical Wnt signaling (Af and At modules in both shared and individual datasets) and retinoic acid (RA) signaling (Af module in the Li et al., 2018 dataset[141]). These findings implicate Wnt and RA pathways as potential regulators or effectors of the developmental programs active in these regions. The enrichment of the "cellular response to retinoic acid" category in Af GMs aligns with our previous findings, which showed increased expression of genes involved in RA synthesis, signaling, and downstream targets, such as *CBLN2* and *MEIS2*, in the midfetal frontal cortex[83,106]. In the present study, we further demonstrate that the expression of key pericentral GM genes, most notably *Plxnc1*, is regulated in part by RA signaling in the mouse medial prefrontal cortex (mPFC) at birth, a developmental stage equivalent to midfetal human cortex. These results underscore the broader potential roles of RA and Wnt as paracrine morphogens and signaling molecules in the development and patterning of association cortices.

Previous genetic studies have shown that RA is not required to regulate early patterning of the forebrain but that it functions in a paracrine fashion later during generation of early neurons of the rostral ventral pallium where its synthesizing enzyme ALDH1A3 is expressed by radial glial cells and other cell types of the lateral ganglionic eminence (LGE) and septum[142–144], the most anterior structures of the ventral pallium that are also adjacent to the antero-ventral-most portion of the emerging cortical plate. Supporting this, we find that both PLXNC1 and ALDH1A3 are expressed in the LGE and septum, adjacent to the ventrally located allocortical plate, where PLXNC1 expression first emerges in the dorsal pallial cortical plate.

Previous studies have demonstrated that only very low levels of RA are present in the early nascent neocortex itself[106,142,144–146], suggesting that source RA may be diffusing from neighboring ventral pallial structures, generating a diffusion gradient that helps initiate the expression of *Plxnc1* and maybe other pericentral genes. RA is also produced by the cortical meninges[142,144,147], and, at a later developmental time, by the anterior ventromedial frontal cortex, the region from which the Af program emerges within the neocortex[105,106]. Here presented and previous analyses[105,106,142]. *RARE-LacZ* RA signaling reporter mice revealed that signaling emerges first in the early (around PCD 11.5) LGE and striatum, and later in development appears in meninges, and in the antero-medial frontal cortex, hippocampus, and postero-lateral amygdalohippocampal cortex temporally, which to some extent overlap with spatiotemporal articulation of the pericentral programs. Notably, this later emergence of RA in the immature cortex coincides with expression of key RA regulated genes such as *Cbln2*, *Meis2* and *Plxnc1*, and the onset and gradual inward progression of the pericentral programs. Importantly, the incompleteness of this overlap also strongly suggests the role of other factors in the induction and articulation of those programs.

Of potential relevance to this scenario, mice that ectopically overexpress FGF8 in the early neocortex, or those lacking FGF8, which show loss of anterior ventral pallial structures including the septum, LGE, and medial ganglionic eminence (MGE), exhibit altered expression of *Lmo4*, a gene enriched in the early medial mPFC and included in the pericentral Af GM[148–153]. Notably, the septum and LGE are critical early sources of anterior RA, even though neither FGF8 nor FGF17 appear to be produced by mPFC neurons. Furthermore, the transcription factors ZIC1, ZIC3, and ZIC4 are enriched in progenitors and other cell types within the developing septum, and to a lesser extent in other ventral pallial structures and the anteromedial frontal cortex across mice, macaques, and humans[83,154,155]. Regulatory gene network predictions also suggest that ZIC transcription factors may function upstream of *Fgf8* and *Fgf17* in these developing structures[154]. Genetic deletion of both *Zic1* and *Zic3* in mice leads to defects in the medial pallium[155].

Together, these findings suggest a mechanistic link between the previously described *Fgf8*-, *Fgf17*-, or *Zic*-dependent development of the septum and other anterior ventral pallial structures[148–153,156], which are the earliest sources of RA and other factors in the anterior ventral pallium[106,142,144–146], and the induction of certain pericentral Af genes, such as *Plxnc1*, and their role in PFC development. However, it is unclear if this is a molecularly direct or indirect link due to developmentally affected anterior ventral pallial structures in mice hypomorphic for or lacking *Fgf8*, *Fgf17* or *Zic* genes. Notably, RA deficiency affects FGF and SHH signaling in the ventral forebrain[157]. However, it is worth noting that apparently neither the ventral VZ expression of *Meis2* or *Fgf8* require RA[143], further supporting a scenario for the involvement of additional factors in regulating these and other relevant genes. Thus, based on these observations, RA signaling alone is unlikely to fully account for the complex and prolonged spatiotemporal dynamics underlying the induction and articulation of pericentral programs.

A limitation of our study is that there are likely additional molecular and cellular mechanisms regulating the initiation and articulation of the Af and At programs which remain unidentified. The lists of genes in the shared GMs were curated using stringent criteria across three independently generated data sets of human (2 datasets, microarray and RNA-seq) and macaque (1 RNA-seq dataset) through different methods. This limited approach hindered our ability to expand our lists in a rigorous manner and fully resolve cell type–specific signatures, particularly those exclusive to immature ExNs. One mechanism we considered involves the well-established reciprocal interactions between thalamic and neocortical development. While the influence of FO TCAs is essential for inducing central programs and sensorimotor arealization, our data suggest that such a direct and inductive relationship may not hold in the case of HO association TCAs, cortical pericentral programs, and early transmodal patterning. Specifically, our analysis of PLXNC1 expression and cortico-cortical projections to the mPFC reveals that key molecular features of pericentral programs, as well as transmodal-like connectivity patterns, can emerge even in the absence of TCAs in the neocortex, as observed in *Satb2* and *Zbtb18* conditional knockout (cKO) mice, or when TCAs are severely reduced, as in *Celsr3* and *Gbx2* cKO mice. A previous study using full-body *Gbx2* knockout mice[150] found that certain aspects of intra-neocortical projection patterns were not disrupted in the absence of thalamic input when analyzed at postnatal day 0 (PD 0). This finding is consistent with our observations in *Gbx2* conditional knockout (cKO) mice, in which we observed an expansion of temporal and (para)-limbic association projections into the mPFC along the dorsoventral (D–V) axis in the days that followed. Specifically, by PD 37 in *Gbx2* cKO mice, when associative connections between the mPFC and temporal or paralimbic association regions are typically established, these long-range projections into the mPFC had expanded into medial cortical territories that are normally occupied by primary auditory and somatosensory areas. These results suggest that thalamic input is not required for the initial formation of intra-neocortical projections, but is necessary for their exclusion from, or refinement within, specific cortical domains.

Although the timing of birth of FO versus HO thalamic nuclei differs markedly between mice and macaques, the overall spatiotemporal pattern of pericentral program development appears to be conserved across species. Nevertheless, HO thalamic nuclei exhibit a broader, yet still specific, pattern of areal innervation and, as such, likely influence the development of certain molecular, cellular, and physiological features of the higher-order association cortex.

Lastly, we also noticed that the general spatiotemporal pattern of the pericentral programs in the neocortex follows the general pattern of progression of cell differentiation and maturation, such as synaptogenesis, dendritogenesis, and long-range intra-cortical connectivity; it is, however, inversely correlated with the pattern of neocortical myelination which first appears in primary sensorimotor regions and last in transmodal regions. Whether opposing programs described here contribute to or are reinforced by those general maturational trends remains to be explored.

Taken together, we hypothesize that the induction and articulation of pericentral programs are governed by a combination of passive and instructive cues, including RA. Future studies aimed at dissecting the regulatory mechanisms controlling *Plxnc1* and other key pericentral GM genes in the developing forebrain may provide critical insight into this developmental process.

**The concept of central “sensorimotor” identity programs**

Our current understanding of the mechanisms underlying neocortical patterning primarily stems from experimental studies on intrinsic and extrinsic factors involved in sensorimotor arealization. Intrinsic mechanisms focus on the concept of a molecular gradient-organized "protomap" within progenitor cells[158], where specific secreted signaling molecules and transcription factors with graded expression patterns regulate the position and size of primary sensorimotor areas[127,148,159–161]. The reciprocal interactions between sensorimotor thalamic and neocortical development represent extrinsic mechanisms[1]. This close relationship is exemplified by experiments demonstrating the instructive, activity-dependent roles of sensory input from peripheral organs and their thalamic relay nuclei in establishing primary area-specific properties[2,4,5]. The sensorimotor TCAs maintain a topological register that organizes primary topographic maps, which then serve as activity-dependent and neuronal adjacency-based anchors for the patterning of higher-order networks[3,19,58]. This mechanism preserves cortical adjacency and sensoritopic relationships while meeting local functional requirements across various levels of the sensorimotor hierarchy. These properties appear to be determined regardless of neuronal position within the neocortical surface or the gradient-organized protomap, within the context of what has been posited the early pluripotent "protocortex"[59].

Furthermore, these studies have revealed the inductive capacity of the peripheral sensory signals and FO TCAs in specifying the location and size of prospective primary sensorimotor areas, as well as inducing key aspects of layer- and area-specific gene expression, cytoarchitecture and cortico-cortical connectivity[29,49–53,55,56,58]. For instance, primary sensory areas, particularly layer 4, exhibit strong sensory specialization. These areas adopt a sensory-petal organization, forming a somatotopic, retinotopic, or tonotopic map that mirrors the layout of the peripheral sensory organ, as demonstrated by classic studies[121,162,163]. Thus, sensory-petal FO TCA targeting and induction do not affect an entire primary area en bloc. Instead, it gives rise to distinct laminar patterns. For example, in primates, the border between areas 17 (primary visual) and 18 (secondary visual) is sharp in layer 4 but appears more gradual or even absent in other layers. This suggests that induction operates by conferring properties onto individual neurons rather than uniformly across progenitor populations or the cortical plate. Whether subpallial structures are involved in this process remains unclear. It is likely that, due to this mode of action, progenitor cells are not directly involved in the induction and exclusion processes.

Additionally, the absolute size of sensory or motor areas can expand or contract by widening or shrinking the cortical representation of a sensory input, driven by peripheral development of enhanced or reduced sensory or motor functions. For instance, reductions in

LGN volume, optic nerve diameter, and occipital cortex grey matter have been observed in albinism likely due to retinal abnormalities and decreased foveal receptors[164,165]. Alternatively, the relative size of sensory or motor areas can change due to a change in the overall size of the cortex. One illustrative observation is that the relative size of the optic tract, LGN, and V1C are highly coordinated within an individual and do not vary greatly between individuals, yet total cortical size varies orders of magnitude more between individuals[166]. Therefore, individuals with larger cortices, on average, will have a relatively smaller V1C, while the opposite is true for individuals with smaller cortices. The proceeding examples highlight that the ultimate driver of the cortical area devoted to a sensory system is the number of sensory receptors in a given sensory organ and their specialization. This point is further illustrated by the observations that eye volume and visual cortex area are correlated[167], and majority of the occipital cortex responds to central vision, a narrow region in the visual space, corresponding to the photoreceptor dense fovea[168].

The functional consequences of reduced peripheral input to the human cortex are illustrative in congenital blindness, where the territory corresponding to primary visual cortex develops in the absence of peripheral visual input. Rather than remaining silent, the occipital cortex in these individuals is consistently activated during nonvisual tasks such as braille reading and auditory language processing[169–171]. Because the occipital cortex activates bilaterally during braille reading, which is performed with a single dominant hand, it is unlikely that somatosensory TCAs are rerouted to the occipital cortex to a significant extent and that this is a primary sensory response[170]. Some have concluded that large portions of the visual cortex in the congenitally blind appear to be functionally integrated with higher-order association networks and higher cognitive functions[170,172]. This may reflect reduced central V1C programming and expanded pericentral programming. However, this is merely speculative as there is also no direct evidence demonstrating increased anatomical transmodal connectivity or pericentral gene expression within occipital regions typically corresponding to V1C in the congenitally blind. These observations are based on functional coupling. Nevertheless, these findings underscore the developmental plasticity of the naïve neocortex and the remarkable functional changes, adaptations, and repurposing of occipital cortex, possibly towards higher-order processes, observed in the human brain in the absence of peripherally driven visual input during development.

The concept of central “sensorimotor” programs incorporates some of these key previous findings and concepts, predominately building on previous research highlighting the instructive role of the FO sensorimotor TCAs in guiding early neocortical neurons to acquire features of primary areas[7,9,73,88,91,92,96,105,121–126,158,173–187]. We incorporate this understanding in our integrative model by identifying two mechanisms (i.e., induction and exclusion) linking sensorimotor TCA-driven gene expression to the emergence of both primary sensorimotor and unimodal association area-specific cortico-cortical connectivity. Furthermore, the principles of induction and exclusion direct and constrain experience-driven modifications throughout the sensory and motor regions. Induction is exemplified by sensorimotor TCA promoting the formation of primary areas and enriching SEMA7A expression within them, while excluding PLXNC1, an axon guidance molecule with anticorrelated expression that acts as a repulsive counterpart in this process and is expressed by inward-moving pericentral association programs. Consistent with the principle of induction, SEMA7A expression is highly specific to primary areas, where previous studies have shown that it is induced and regulated by FO TCAs in an activity-dependent manner, controlling dendritic growth and synaptic connectivity[63,65,67,68,71,73,74]. Building upon these foundations, we reinforce the notion that these processes amplify the expression of some of the genes that might be expressed at low levels in nascent neocortex and drive the nested and elevated expression of those and other select genes across primary sensorimotor areas.

**Topographic and functional organization of intra-cortical networks**

The mature cerebral cortex exhibits a hierarchical organization involving S–A networks, and the structure of these networks varies across mammalian species. As such, the biological basis of conscious perception, complex cognition, and voluntary behavior depends, at least in part, on the topographic and topological patterns of long-range intracortical connectivity along this axis[15,16,18–21,23,32,118–120,188–210]. At one end of this hierarchy lie the unimodal primary sensory and motor areas with dense local networks that interact with the environment and body by processing early sensory inputs and executing motor behaviors, respectively[48]. At the other end are higher-order association areas distributed across the prefrontal cortex, posterior parietal cortex, and large parts of the temporal cortex that either surround peripherally or interpose centrally between the primary and unimodal sensorimotor association areas[15]. These transmodal areas are interconnected by long-range anteroposteriorly oriented axons, forming highly distributed and parallel processing networks that integrate information across modalities to support complex cognitive functions including language, memory, attention, social cognition, and abstract reasoning[15,21,193]. Transmodal networks interact less directly with sensory input or motor output but are highly interconnected with the limbic and distributed association networks of the three-layered allocortex, which are crucial for processing and regulating functions such as emotion and memory[15,193,211,212], with which it shares certain molecular and hodological similarities[24,83,85,190,213,214].

**Establishment of large-scale inter-areal networks along the S–A axis**

Functional maturation is unevenly distributed across large-scale intracortical networks and follows a prominent pattern of hierarchical progression along the S–A axis[15]. These maturational changes are associated with the earlier development of sensorimotor abilities and the later emergence of complex cognitive and social functions. This function must be based in structural organization stemming from developmental mechanisms shaping the S–A axis. However, as described in the introduction, despite significant advances in our understanding of primary and unimodal sensorimotor cortices, fundamental questions remain regarding the formation and evolution of the S–A axis, particularly the transmodal cortices. These unresolved issues include: (1) Are transmodal cortices patterned by the same mechanisms that shape unimodal cortex?; (2) How do transmodal neurons establish long-range connections across vast distances while avoiding primary areas?; (3) Why do distributed transmodal networks feature fronto-parieto-temporal transmodal hubs and frequently engage PFC?; (4) Why do association areas exhibit graded transitions, in contrast to the sharply defined boundaries delimiting primary areas?; and (5) Has the evolutionary expansion of transmodal cortex in primates necessitated distinct developmental mechanisms?

Our findings and model address the above questions and leave open room for further analysis. (1) Patterning of the S–A axis is regulated by competing molecularly and spatiotemporally distinct pericentrally and centrally emerging programs that emerge and progress from opposite sides of the developing axis. Together, these opposing programs impart distinct molecular characteristics that shape the archetypal features of the S–A axis. (2) The mutual repulsion observed between primary and association areas is a key feature that facilitates long-range associative connections across vast distances which strategically avoid primary areas. The role of *SEMA7A* and *PLXNC1*, along with other axon guidance and cell-cell adhesion molecules, is to, in part, mediate repulsion between sensorimotor and association areas. Parietal association cortex, which expresses *PLXNC1* but not *SEMA7A*, is integrated with the frontal and temporal association hubs despite being intercalated between primary sensorimotor areas. Our model predicts that this is due to the spread of pericentral programs towards the neocortical center and lack of central program induction in this area, allowing the acquisition of association area features. (3) The F–T axis is seeded by early-born deep-layer ExNs and supports the stepwise inward expansion of association networks along the fronto-temporal axis, following the progression of pericentral programs. (4) Our model predicts that transmodal networks or connectivity hubs emerge as gradients within a distributed architecture. Opposed to the sharp delineation of focal areas driven by sensoritopic TCAs, the fronto-temporal emergence and propagation of pericentral programs supports the creation of gradients and distributed architecture rather than the sharp delineation of specific sub-networks. This supports the observation that specialized cognitive networks and modules gradually develop and refine over the course of an individual's development. Along with this, there is no evidence of area- or network-specific gene expression associated with particular transmodal networks. Rather than a single gene underlying a distinct process, like language, we propose that groups of genes within pericentral programs contribute to association pattering, highlighting a shared genetic basis underlying diverse cognitive functions. (5) The opposing patterning programs, along with inductive and exclusionary principals, appear evolutionarily conserved. These observations also support the notion that transmodal networks and mechanisms underlying their development are ancient as opposed to recent evolutionary inventions. These findings and our model indicate that disproportionate expansion of transmodal cortex in primates does not necessitate distinct developmental mechanisms. Furthermore, unlike in mice, the primary sensory thalamic nuclei in macaques are generated later than the medially located association nuclei. This delayed generation may allow the invading pericentral programs to occupy more cortical territory in the absence of FO sensorimotor thalamocortical axons. Consequently, the expansion of transmodal areas in primates does not need a separate molecular mechanism or to be disconnected from primary sensorimotor anchors; rather, they leverage the same pericentral programs that operate in other mammals and the principles of induction and exclusion in competition with the central programs. Thus, there is no need for primate-specific mechanisms to explain the expansion of the transmodal cortex with the increasing size of the neocortex.

**Central programs shape sensorimotor networks**

Our findings reveal that SEMA7A regulates the nested pattern of cortico-cortical connectivity within sensory cortices and, through its interaction with PLXNC1, restricts the acquisition of certain connectional features characteristic of the association cortex. The pericentral programs are not evenly distributed along the peripheral circumference but instead emerge at specific nodes, driving polarization along F–T trajectories spreading inwards along this axis toward the central territories not yet fully influenced by the sensorimotor TCAs. However, this progression is interrupted by areas and layers rich in molecules of the central programs, such as SEMA7A, as well as by other factors like CYP26B1 in the prospective antero-lateral motor cortex, which limit their influence. This is particularly evident in the thalamo-recipient layer 4 of primary sensory areas, which exhibit the highest levels of SEMA7A expression, and in layers 2 and 3 which express slightly lower levels of SEMA7A. Concurrently, we observed expression of some pericentral effectors including *Pcdh10*, *Pcdh17*, *Plxnc1,* and Meis2 in the deep-layers of central regions which express little or no SEMA7A. The absence of primary areas in SATB2 and ZBTB18 cKOs might be the result of defects in the guidance and fasciculation of TCAs in the ventral forebrain and/or the expression of early central genes, such as SEMA7A, which may be involved in the growth of TCAs into the cortical plate. In the case of their deletion, TCAs fail to invade the neocortex and, consequently, do not form the necessary connections. Additionally, we observed that the intensity of SEMA7A expression varies across different primary sensorimotor areas and between homologous areas in different species. Notably, its expression appears to be highest in the dominant sensory modality, which generally exhibits the most well-differentiated cytoarchitectonic features and distinct sharp areal borders with surrounding association areas. For example, the somatosensory barrel fields (S1C) in mice and the primary visual area (V1C) in primates. Based on previous results and our findings, we hypothesize that sensory TCAs induce an activity-dependent mechanism that drives the modality-specific enrichment of SEMA7A expression. Consistent with this, the primate V1C and rodent S1C have the thickest primary thalamo-recipient layer, layer 4. This enrichment, in turn, facilitates the formation of highly differentiated cytoarchitectonic features and cortico-cortical segregation at the borders with neighboring higher-order areas. Furthermore, we observed a strong correlation between lower density of primary sensory TCAs in mouse S1C and lower intensity of SEMA7A immunoreactivity in different mutant mice that lack obvious well differentiated primary areal cytoarchitectonic features and borders.

**Pericentral programs shape association, including transmodal, networks**

In mice, the onset of the Af program occurs around the antero-medial frontal cortex neighboring the septal complex and anterior olfactory nucleus (septal pole) while the At programs emerge around the postero-lateral amygdalohippocampal temporal cortex (amygdalar pole) in alignment with the observation that the anterior olfactory nucleus and amygdala are among the earliest cortical structures to be differentiated[215,216]. While this onset coincides with early neurogenesis and maturation of these regions at the ends of the geometric fronto-temporal axis, neither programs progress inwards with age along the dominant ventro-dorsal (V–D) axis (ventral insular paleocortex to mediodorsal archicortex) of neurogenesis[217]. We propose that this early and global polarization along the fronto-temporal axis and propagation of opposing central and pericentral programs, help shape many key features of the intra-cortical networks, such as gradients, matching of areal and laminar patterns of high-order feedback and low-order feedforward connections. The concept that early neocortical spatio-temporal patterning of association areas progresses inward along the topographic fronto-temporal axis and while the primary areas emerge subsequently as focal islands located more centrally is consistent with the spatio-temporal progression of cytoarchitectonic differentiation of the human neocortical plate previously described by Kahle in the 1960s[218]. He described what he called developmental "graduations" in the cytoarchitectonic differentiation of the fetal human neocortex. These graduations first emerge at the frontal and temporal polar borders adjacent to the ventral allocortex and gradually spread inward toward the center. It is in this central region that, later in development, the first signs of cytoarchitectonic differentiation associated with the primary sensorimotor areas appear. Complementary spatiotemporal cytoarchitectonic patterns were also observed using various modalities, further reinforcing that, even in the large human brain with its protracted developmental timeline, regional patterning and specification of the prospective higher-order association areas in the frontal and temporal cortex emerge early in fetal development[219–221].

However, the nodal patterns and temporal progression of pericentral and central programs cannot also be fully explained by the timing in the birth (temporal proximity or matching, i.e., born together wired together)[222,223], spatial proximity, or the timing of invasion of different TCAs. This is due to our observation that the spatiotemporal appearances of the pericentral and central programs follows a similar pattern in primates and mouse, but the correspondence in the timing in the neurogenesis of major sensorimotor and association thalamic nuclei and their respective neocortical areal targets differ greatly. In general, in mice, the FO sensorimotor thalamic nuclei tend to be generated earlier during development than the HO association nuclei. Conversely, in macaques, and likely in humans as well, the developmental timeline is somewhat reversed, with HO nuclei emerging slightly earlier or concurrently with FO nuclei. The delayed birth of the primary sensory nuclei compared to mouse could explain why primary sensory areal differentiation appears relatively later in non-human primate and human neocortex compared to the F–T intracortical cytoarchitectonic "graduation" trends. On the other hand, Kahle and others have also observed that the early cytoarchitectonic differentiation of ventrolateral paleocortex and insular meso/neocortex as well as complementary dorsomedial archicortical differentiation seems to be more spatio-temporally restricted. This contrasts with the fact that the anti-hem signaling region and the paleocortex are largest on the ventrolateral side while the embryonic hem signaling region and associated archicortex are elongated along the dorsomedial and dorsoventral edge of the cortex, and the main V–D neurogenetic trend of the neocortical plate. Also, the role of hem and anti-hem have been largely associated with the formation of archicortex and paleocortex, receptively. We also hypothesized that the cytoarchitectonic gradients described reflect in large part the developmental trends in the patterning of the neocortical plate rather than evolutionary trends. Paleocortical ExNs, which are phylogenetically and ontogenetically older than neocortical ExNs, tend to form diffuse projections within distributed limbic structures and preferentially target pericentrally located transmodal neocortical regions while largely avoiding primary sensorimotor areas, consistent with those neurons abundantly expressing *Plxnc1*.

In the absence or reduction of primary areas, "association-like" neurons targeting the mPFC span nearly the entire cortex, maintaining alignment along the fronto-temporal axis, which serves as a key axis of long-range association connectivity. This axis is seeded by early-born deep-layer ExNs and supports the stepwise inward expansion of association networks along the fronto-temporal axis mainly from ventrolateral (and to some extent dorsomedial) regions. We hypothesize that these principles influence adult features like areal and laminar connectivity, dendritic organization, cytoarchitectonic gradients, and gyrification[107,117–120,224].

**Insights into cortical geometry and gyrification**

The initial fronto-temporal polarization trend of the prospective association neocortex is facilitated and maintained by multiple factors, while the V–D axis (ventrolateral paleocortex to dorsomedial archicortex) seems to be less prominent in the context of our model. The geometry of cortical curvature and gyri is thought to influence the spatial expression of putative patterning molecules through a hierarchical reaction-diffusion mechanism, thereby imposing geometric constraints on early developmental programs[225,226]. Furthermore, the spatiotemporal expression of CYP26B1, an enzyme that degrades RA in the frontally regulated program, provides an insight into how molecules with polarized expression along the V–D axis may promote polarization of signaling pathways and genes along the F–T axis. CYP26B1 is highly enriched in the prospective insular paleocortex and periallocortex expanding towards the prospective motor-somatosensory regions as well as the dorsomedial archicortex and mesocortex during prenatal development. Furthermore, the pericentral programs emerge early on at frontal and temporal polar nodes or points of interaction with allocortex. While anterior (septal complex and anterior olfactory nucleus) and posterior (i.e., cortical amygdala, piriform-amygdalar area and postpiriform transition area) paleo- and peri-cortex are likely bordering regions that express *Tfap2d* around PCD 13.5 in mice when we first observed *Plxnc1* expression in the neocortical plate. The *Zbtb20* expression domain in the archicortex is being just established as it is generated after the first most ventral paleocortical structures. Nevertheless, the frontal and temporal nodes are also where most anterior and posterior transitional zones of the archicortex also neighbor the two paleocortical poles. As far as we are aware with the evidence, the frontal node is likely the peri-archicortex of mainly the antero-medial subcallosal and subgenual areas while the temporal node consists mainly of the anterior part of parahippocampal gyrus (i.e., perirhinal and entorhinal cortex), which would correspond to the postero-lateral region of the temporal cortex in the mouse.

The geometric bending of the frontal pole and more prominently the temporal poles around early prenatal insular axis in species like anthropoid primates, curves the fronto-temporal axis. Initially during fetal development, the insula is exposed and matures faster and earlier than the surrounding cortical areas[227]. This early maturation causes the insula to serve as a stable anchor point around which the surrounding frontal, parietal, and temporal areas grow, elongate, and bend, which encapsulates the insula resulting in the generation of the surrounding operucula (opercularization) and formation of the sylvian fissure. The opercular gyri covering the insula contain numerous cortico-cortical axons connecting frontal and temporal transmodal cortices[228,229] and appear early in fetal development in humans and macaques[224,230]. This growth, elongation, and bending around the insular axis causes the temporal node/pole to move anteriorly in comparison to V1C[227]. The C-shaped distortion resulting from this process is likely caused by multiple factors, including mechanical tension along the F–T axis, differential areal growth rates, axon tension, and physical constraints imposed by the cranium. F–T elongation and cortical distortion result in the emergence of a “false” anatomical occipital pole, in addition to the “true” topographic frontal and temporal poles[231]. In this respect, V1C is geometrically dorsomedial and just posterior to the center of the neocortical sheet. Therefore, in such a position, the axonal systems originating from the V1C forming the dorsal (also known as the “where pathway”) and ventral visual streams ("what pathway") in the unimodal association cortices of primates also elongate along the F–T axis. Furthermore, the non-polar location of the visual cortex is apparent in cetaceans, which, relative to humans have massively distorted F–T axes. Specifically, bottlenose dolphins have massive temporal lobes and relatively small frontal lobes, and sensorily rely predominantly on auditory processing[232,233]. In these brains, compared to primates, the visual cortex is positioned in a more dorso-lateral position adjacent to the larger auditory cortex[232]. Consistent with the F–T polarization gradient, the mesoscale connectome of the mouse neocortex predominantly follows an anteromedial (mPFC) to posterolateral orientation of long-range distributed association projections[202–204,234]. In contrast, the output of the primary visual cortex has short-range outward visuotopically organized projections into surrounding neighboring association areas[235].

It has previously been proposed that mechanical tension along both short- and long-range cortico-cortical axons contributes to various aspects of cortical morphogenesis, including cortical folding[236]. In line with this model, we propose that the F–T polarization helps to guide the earliest frontal and temporal long-distance cortico-cortical axons and generates mechanical force that contribute to a specific cortical folding pattern by inducing anisotropic tension along the F–T axis. This mechanical tension, combined with the inherent C-shaped distortion of the cortex in some species like humans, promotes a longitudinal (F–T oriented) folding pattern and the formation of gyri, such as the cingulate gyrus and the superior, middle, inferior frontal, and temporal gyri. It also contributes to the development of bent gyri, such the angular and supramarginal gyri. Indeed, the gyrification index, a measure of cortical folding, is highest in the human prefrontal and parieto-occipito-temporal association cortices, reflecting their dense long-range interhemispheric and callosal cortical-cortical connectivity[224].

The MIND model also provides potentially important implications as to why the main gyri of the prefrontal, cingulate, perisylvian, and temporal cortices align along the F–T axis. The topography of cortico-cortical axonal pathways plays a key role in the formation of gyri and we hypothesize that this pattern is seeded in the early patterning or polarization of early-born deep-layer cortico-cortical neurons of the frontal and temporal poles. The axons from these neurons are pioneers of the longest fiber systems within the cortex that connect certain association and limbic regions. The orientation of the main gyri of the association cortices, therefore, are the consequence of the formation of the earliest axons of the cingulate medially, uncinate fasciculus laterally, and frontal and temporal long-range association axons of early born deep layer neurons guided by the At and Af programs expanding along the F–T axis. These gyri are interrupted orthogonally by the centrally located precentral and postcentral gyri associated with primary motor and sensory areas, respectively. These gyri are strongly connected by U shaped fibers originating at least in part form later born upper layer neurons forming the mirror representation of the motor and sensory homunculi or in the case of the calcarine fissure, the upper and lower visual fields[228,229]. While the central sulcus and calcarine fissure appear earliest, this likely represents the faster activity driven maturation of short-distance sensorimotor projections. The protracted development of the distributed association networks likely results in the delayed emergence of their associated gyri, despite the axons of these networks being the first established and having elongated topography. Their slower maturation is most evident by the delayed appearance of fully myelinated axons in transmodal cortices which do not appear until well after early postnatal development in humans[32,34]. Thus, our model also provides some explanatory values as why major transmodal long-range connections and associated gyri are elongated along the F–T axis and why it is the frontal and temporal lobes that are prominent and places of developmental and evolutionary expansions. Dedicated studies are needed to test and clarify these points.

**Insights into the evolution of the neocortex**

Our findings and model provide potentially important insights into the processes underlying the evolution of the neocortex. Nervous systems did not evolve in isolation but rather in the context of the body, behavior, and ecological niche. In this regard, the areal map and networks of the neocortex encapsulate both the body and the environment. Primary and unimodal association sensorimotor cortices appear to be under stronger adaptive pressures, allowing for specialization, whereas transmodal association networks may be more innate and even hardwired to some extent but will change in a complementary manner to varied unimodal areas, including expansion and contraction. Interestingly there is a negative allometric scaling of eyes and the visual LGN across primates and apes, with humans having both relatively and absolutely the smallest primary visual cortex among great apes[237]. These linked patterns of absolute and relative complementary expansions of transmodal networks that may lead to emergent properties of scaling distributed networks and representations.

Like the principle of Turing's reaction-diffusion model[225], size matters: when FO sensorimotor TCAs innervate a smaller neocortical sheet, they occupy a proportionally larger territory and create broader exclusion zones. These zones limit the inward expansion of pericentral association programs. Furthermore, the cortical neoteny in large-brained mammals, in particular the waiting of TCAs in the subplate zone, delays the ingrowth of TCAs into the cortical plate further after the onset of pericentral programs which appear to happen much earlier. This developmental delay will allow the pericentral programs to occupy more territories before being limited by the induction of primary areas. Thus, our model offers a simple mechanistic explanation for the variation in the size, modular organization, and location of cortical areas and networks, both primary sensorimotor and association, across species. Furthermore, it suggests that the expansion of human association cortices represents an adaptation based on strong adaptive pressures to enlarge the size of the neocortical sheet and decrease the size of territories occupied by the primary sensory and motor areas.

These differences are shaped by factors such as overall cortical sheet size, the timing of cortical and thalamic neurogenesis, the heterochronicity between central and pericentral developmental programs, and peripheral influences. We show that in the absence of primary areas, or without neocortex-enriched markers such as SATB2 or ZBTB18, the neocortex does not default to an allocortical fate. We also observed a subtle shift in the boundary between the neocortex and the peri-paleocortex/piriform cortex in SATB2 and ZBTB18 cKOs. This suggests that these two transcription factors, enriched in the neocortex and expressed at lower levels in the archicortex, play a regulatory role in establishing the boundary between the (peri-) paleocortex and the neocortex. This finding complements previous work highlighting the central role of the transcription factor NR2F1 (also known as COUP-TF1) in progenitor cells in defining the border between the neocortex and the medial entorhinal cortex, a transitional peri-archicortical region[238]. Despite these shifts, much of the dorsal cortex in SATB2 and ZBTB18 cKOs retains key characteristics consistent with a neocortical identity. For instance, the neocortex in both mutants exhibits widespread connectivity and molecular profiles typical of higher-order and transmodal association areas. Along with prior studies[128,132,153,158,161,213,214,238–247], this supports the scenario where the identities of the allocortex and neocortex, as well as the boundaries between them, are specified by mechanisms acting both at the level of molecular-gradient "protomap" in cortical progenitors and through the compartmentalized and graded expression patterns in nascent postmitotic neurons. Furthermore, this supports the view that the neocortex evolved en bloc, crowning the allocortical ring, and that the dominant growth axis for long-range association axons is F–T, rather than latero-medial or V–D. The early long-range association networks are seeded by early-born deep-layer ExNs in frontal and temporal polar regions and expand progressively inward along the F–T axis mainly from ventrolateral (and to some extent mediodorsal) cortical regions. In contrast, the V–D axis appears to be dominant of cortical neurogenetic gradients.

A key distinction among mammalian neocortices is the significant variation in the presence and size of primary sensorimotor areas versus transmodal regions in the frontal, parietal and temporal lobes. Consistent with the anchoring role of primary sensorimotor areas in neocortical patterning, the oldest living mammals, such as monotremes and marsupials, possess relatively small brains with a large olfactory (piriform) allocortex and a small neocortex composed predominantly of unimodal sensorimotor areas. In contrast, phylogenetic comparisons suggest that in large-brained placental mammals, such as primates, transmodal areas and distributed networks have greatly expanded within an enlarged neocortex and recent evolutionary specializations. This dramatic expansion in primates has been proposed to have "untethered" transmodal zones from the patterning influence of primary sensorimotor areas, conferring unique properties that contribute to distinct structure-function relationships along the S–A network hierarchy[19].

A contrasting phylogenetic perspective on arealization is embodied in the "Dual Origin of the Neocortex" hypothesis, which posits that primary sensorimotor areas are the most evolutionarily recent due to their highly differentiated cytoarchitectonic laminar structure[24,206,248–254]. According to this theory, the neocortex evolved gradually from the piriform cortex (paleocortex) and hippocampus (archicortex), with concentric rings of increasing laminar complexity emerging from paralimbic areas and extending toward the primary sensorimotor regions. The hypothesis further divides the neocortical surface into two distinct regions: the dorsomedial division, derived from the archicortex, and the ventrolateral division, derived from the paleocortex. While the cytoarchitectural laminar gradients in the adult neocortex identified and utilized in this hypothesis are largely accurate, the concept lacks grounding in contemporary understandings of neocortical evolution, as it conflates simpler laminar architecture with phylogenetic antiquity and more complex lamination with evolutionary recency[210,255]. Furthermore, this dual evolutionary hypothesis is inconsistent with recent findings demonstrating a distinct transcriptomic signature across the ring-like organization of the mesocortex, a transitional region between the peripheral allocortex and the central neocortex[106,214,256]. Nevertheless, the "dual origin" concept was revisited following the discovery of systematic relationships between macroscale cortico-cortical connection patterns and spatially ordered changes in laminar differentiation, exemplified as cortical gradients. Specifically, as laminar complexity increases from paralimbic areas toward primary cortices, the origin of PFC-directed projections shifts from predominantly deep to predominantly upper layers. These and other gradients have provided foundations for multiscale models of cortical networks and hierarchy.

Based on our findings, model, and interpretations, there are some similarities and resemblances to the Dual Origin hypothesis, but important differences. Our findings suggest that certain primary and association cortical features are conserved between mammals and birds, supporting the concept that the neocortex evolved en bloc within the allo-/meso-cortical ring[210,255–257]. The neocortex can be thought of as a single structure that emerged in evolution, like an elongated (along F–T axis) elliptical dome crowning the hemispheres above the allo-/meso-cortical ring[257], which exhibits a unique S–A topography shaped by the evolutionary adaptations acquired by a given organism to suit its ecological niche and intrinsic mechanisms consistent with our pericentral programs. They key elements we implicated in S–A axis development (pericentral and central programs) are present in all mammals we analyzed, as well as in the chicken. The central sensorimotor programs depend on the ingrowth of FO TCAs, and in chicken this input is minimal, restricted to a narrow region of the dorsal pallium known as the interstitial nucleus of the hyperpallium apicale (IHA)[258], which contains transcriptionally homologous neurons to mammalian neocortical thalamo-recipient layer 4 and 5 IT ExNs[259]. Consistent with this, the IHA expresses *Sema7a*, and the surrounding regions of the dorsal pallium in chicken express *Plxnc1*. Taken together, these findings suggest that both primary and association features of cortical organization are ancestral to living mammals, and thus likely already present in their last common ancestor. Consequently, simpler cortical lamination should not be interpreted as a proxy to phylogenetic antiquity in all cases.

A notable innovation of anthropoid primates is that they possess a PFC with granular layer 4 that dominates the frontal lobe, with humans having the largest PFC both in absolute and proportional terms, balanced by a large temporal lobe. This enables better integration and gating of multisensory information within the PFC, facilitating top-down control of cognition, attention, and emotions. In contrast, in other analyzed placental mammals, the PFC occupies a smaller area on the medial surface, and the frontal lobe is primarily dominated by motor areas involved in controlling movement and basic behavioral responses. The expansion of the PFC in anthropoid primates underlies the evolution of complex cognitive abilities unique to these species. Our previous research has shown that manipulation of RA signaling, specifically through prospective motor and insular expression of CYP26B1, an enzyme that degrades RA, regulates the areal expansion of the PFC and several genes associated with frontal pericentral programs. This provides experimental evidence of how modulating gene expression gradients can lead to the expansion of the frontal pericentral program and PFC. Our model and experimental findings further indicate that changes in sensorimotor gene expression, such as *CYP26B1*, or the negative allometry of sensory thalamic nuclei and subsequent contraction of primary areas, are an alternative mechanism for increasing the relative and absolute size of the association cortex. Thus, the enlargement could occur either due to a primary expansion of an association cortex, such as the PFC, and/or reduction of primary areas, such as the primary motor cortex. Indeed, the second scenario is supported by our finding of the dorsolateral frontal expansion of PLXNC1 and PFC-like association connectional features in *Monodelphis domestica*, which lacks a true primary motor cortex.

In cetaceans (dolphins, whales, and porpoises) and elephants, which possess the largest neocortex, the frontal cortex is relatively smaller compared to human, with the PFC primarily restricted to the medial and orbital frontal cortex[232,233]. Dolphins, for instance, have a smaller frontal lobe and lack a fully developed olfactory system, while having a massive temporal lobe, reflecting their reliance on sound and biological sonar for navigation[232,233].

Importantly, primate-specific developmental mechanisms are not strictly necessary to explain the expansion of transmodal frontal and temporo-parietal regions. The broader spatial extent of frontal and temporal pericentral programs can emerge naturally as a result of overall brain enlargement and the relative reduction of primary areas, processes regulated by evolutionarily conserved core mechanisms that may or may not undergo significant modification within specific clades to account for unique brain features. For example, both the primary visual thalamic nucleus (LGN) and its primary cortical target, the primary visual cortex (V1C), scale with negative allometry in primates[237]. As the thalamus and neocortex increase in size, these structures occupy a progressively smaller proportion of the neocortex in larger brains, a trend that is especially pronounced in apes, including humans. Thus, the expansion of the transmodal cortex with increasing brain size can be explained without invoking primate-specific developmental mechanisms. Consistent with this scenario, primates, in particular humans, possess a large neocortex[48], ventro-centrally located motor-sensory CYP26B1[106], and smaller FO sensory thalamic nuclei and primary sensorimotor cortical areas[48,237]. Nevertheless, our observations indicate that while the general process is conserved, primates exhibit clade- and species-specific changes in the expression of genes associated with these pericentral programs.

In general, progressive ontogenetic events should not be assumed to reflect evolutionary trends or steps. The same applies to our model, data, and interpretations. Our findings do not suggest that the central region, default state, or different association fields represent stages in the evolution of the mammalian neocortex, which developed later in mammals than the periphery. Nor do they imply that sensory areas emerged by transforming ancestral association areas or displacing existing ones. The naïve neocortex or default state should not be considered the ancestral or generalized cortex of olfactory (paleocortical) origin, as has been proposed for early stages of neocortical evolution[24,260]. The first neurons in the neocortical plate share many developmental programs with the allocortex, as they are all derived from the dorsal pallium. However, the naïve neocortex or default state is not allocortical in nature, as it expresses transcription factors such as SATB2 and ZBTB18, which are enriched in neocortical ExNs to a greater extent than in the allocortex.

Additionally, the idea that the neocortex emerged within the allocortical ring and developed in a maturational sequence from the periphery to the core remains speculative. Lastly, the default program or state likely represents a more generalizable phenomenon in neural development. This is supported by evidence such as the role of default state inhibition in neural induction and observations that PLXNC1, SEMA7A, and various PCDHs exhibit similar temporal sequences and spatial segregation in other early forebrain structures, including the thalamus.

**Insights into the developmental and genetic basis of cognitive modularity**

Our model and developmental framework suggest that the establishment of transmodal networks or connectivity hubs within a distributed architecture, rather than as the sharply delineated units seen in primary sensorimotor areas, is, in part, driven by the gradual inward progression of F–T polarized pericentral gradients. This process describes how specialized cognitive networks and modules, traditionally considered distinct units within the brain responsible for specific functions, gradually develop and refine over the course of an individual's development.

Furthermore, there is no evidence of area- or network-specific gene expression that is exclusively associated with particular transmodal or cognitive networks. We speculate that this developmental framework facilitates the formation of distributed association networks, which support the integration of multiple cognitive processes required to perceive and interpret specific stimuli. Rather than a single language-specific gene, our model proposes that groups of genes within pericentral programs contribute to both transmodal and unimodal association patterning, highlighting a shared genetic basis underlying diverse cognitive functions.

However, the prevailing notion that transmodal systems develop later, particularly in relation to language, is primarily based on functional emergence, delayed myelination, and structural brain imaging studies, or functional imaging techniques. These methods provide only indirect assessments of circuit maturation, rather than direct insights into axonal and synaptic development at the cellular level.

**Potential insights into neurodevelopmental and neuropsychiatric conditions**

Transmodal association areas are disproportionately affected in a range of neurodevelopmental disorders, as evidenced by both structural and functional changes. Large-scale neuroimaging efforts such as the Enhancing NeuroImaging Genetics through Meta-Analysis (ENIGMA) consortium has aggregated neuroimaging data across thousands of individuals and found widespread anatomical differences in psychiatric conditions including schizophrenia[261], and autism spectrum disorder (ASD)[262] with the largest effect sizes typically observed in association regions. In our analyses, we find that genes linked to S–A development are enriched for those associated with ASD risk and intelligence quotient (IQ). These select genes likely regulate early regional patterning and connectivity of the neocortex. This enrichment implies that these developmental processes may be altered in ASD and related conditions. Consistent with this hypothesis, convergent evidence indicates that early cortical development and F–T associative connectivity are disrupted in children with ASD[263–266]. Resting-state functional magnetic resonance imaging (fMRI) studies report increased functional connectivity in prefrontal and temporal transmodal association areas, particularly within the default mode network, and decreased connectivity in sensory-motor and visual regions in individuals with ASD[267–269]. These patterns of functional dysconnectivity are strongly correlated with the behavioral symptoms observed in ASD[269], supporting a link between early network-level dysregulation and clinical phenotypes.

Interestingly, many high confidence ASD-associated genes identified in our region-specific GMs, end up being more broadly expressed across the mature neocortex and are involved in fundamental cellular processes such as synaptogenesis and transcription. Despite their widespread adult expression, alterations in these genes and pathways have been implicated in specific cognitive and behavioral features of ASD, some of which may originate within frontal and temporo-parietal transmodal and (para)limbic networks[263–268,268–270]. As emphasized before[271,272], the early regional differences in the timing of the formation of these cortico-cortical axons and synaptogenesis, protracted maturation and development, and delayed myelination of these networks extends the window of vulnerability to both genetic and environmental perturbations. Our model further posits that the F–T axis is seeded by early-born, deep-layer ExNs that follow the inward progression of pericentral association programs to establish the initial long-range pioneering axons of distributed association networks. Notably, during mid-fetal development, when these axonal projections are growing, certain high-confidence ASD risk alleles are enriched in deep-layer ExNs that normally give rise to these connections [265,273], offering a potential mechanism by which early disruptions in these projection neurons could produce outsized effects on early large-scale cortico-cortical connectivity and related cognitive and behavioral phenotypes.

While these conceptual links are compelling, they must be interpreted with caution, as there is currently no direct evidence linking ASD risk alleles identified in our gene modules to the specific connectional phenotypes discussed above. Further studies will be required to establish direct links between specific risk variants and the architecture and function of S–A networks.

**Limitations, conclusion and remaining questions**

There are several limitations of this study. First, we identified and defined pericentral and central GMs based on bulk tissue-level RNA-seq data from a spatially and temporally limited set of cortical areas in two species. This approach lacks sufficient cell type-specific resolution and fails to capture the full spatiotemporal dynamics, particularly along the D–V axis of the cortex and in numerous other cortical regions. Further study is needed to more precisely determine which cell types express true pericentral or central genes and refine those GMs. Second, our limited transcriptomic spatiotemporal resolution and the histological assessment of a small set of genes within stringently conserved GMs provide only a coarse view of the spatiotemporal articulation of both pericentral and central programs. While the concept of central programs is supported by a larger body of prior literature, the same cannot be said for the pericentral programs. Third, additional evidence is required to clarify which genes within these GMs are driving the programs, which are involved in general maturational processes, and which are indirectly or non-causally included in these modules. Fourth, our understanding of the molecular and cellular processes regulated by either pericentral or central programs remains limited. Although we identified functional enrichment for some pathways, such as Wnt signaling, these were not explored in the current study. Moreover, although prior work has shown functional enrichment for RA signaling in frontally enriched mid-fetal genes (i.e., PFC and motor cortex combined), we only observed enrichment in the Af gene list from Li et al., 2018. We suspect this discrepancy is due to methodological differences and variations in sample selection. Nevertheless, key RA signaling-related genes, such as *Cbln2* and *Meis2*, are present in the most stringent and conserved Af GMs. Fourth, our experimental validation of the role of GM-associated genes in shaping the topographic organization of association axons and in differentiating primary from association areas has been limited to *Plxnc1* and *Sema7a*. While these genes show loss-of-function phenotypes, they appear to account for only a small fraction of the connectivity patterns observed along the S–A axis.

In assembling this extended discussion, we have selectively synthesized a broad body of literature, integrating diverse findings, some emphasized, others less so, into a coherent narrative. While grounded in established principles and experimental evidence, our study also introduces novel insights and mechanisms. Most importantly, the model we propose unifies previous and new findings into a cohesive explanation of how the S–A axis is patterned. It resolves prior conceptual tensions and provides a robust, adaptable framework for understanding cortical development and evolution. Though simplified and not fully capturing the complexity of the cortex, the model accommodates variability across developmental periods, pathological conditions, and evolutionary contexts. As such, it offers a generative foundation for future research into cortical development, evolution, function, and disorders.
